# A Workflow for Improved Analysis of Cross-linking Mass Spectrometry Data Integrating Parallel Accumulation-Serial Fragmentation with MeroX and Skyline

**DOI:** 10.1101/2024.02.13.580038

**Authors:** Juan Camilo Rojas Echeverri, Frank Hause, Claudio Iacobucci, Christian H. Ihling, Dirk Tänzler, Nicholas Shulman, Michael Riffle, Brendan MacLean, Andrea Sinz

## Abstract

Cross-linking mass spectrometry (XL-MS) has evolved into a pivotal technique for probing protein interactions. This study describes the implementation of Parallel Accumulation-Serial Fragmentation (PASEF) on timsTOF instruments, enhancing the detection and analysis of protein interactions by XL-MS. Addressing the challenges in XL-MS, such as the interpretation of complex spectra, low abundant cross-linked peptides, and a data acquisition bias, our current study integrates a peptide-centric approach for the analysis of XL-MS data and presents the foundation for integrating data-independent acquisition (DIA) in XL-MS with a vendor-neutral and open-source platform. A novel workflow is described for processing data-dependent analysis (DDA) of PASEF-derived information. For this, software by Bruker Daltonics is used, enabling the conversion of these data into a format that is compatible with MeroX and Skyline software tools. Our approach significantly improves the identification of cross-linked products from complex mixtures, allowing the XL-MS community to overcome current analytical limitations.

## INTRODUCTION

Recent advances in mass spectrometry (MS) have revolutionized our understanding of how proteins function due to different techniques in structural MS. Among these techniques, cross-linking mass spectrometry (XL-MS) has emerged as a particularly valuable tool.^1–3^ XL-MS provides unique insights into protein conformations and protein-protein interactions.^4,5^ Despite the great potential of XL-MS, there are currently still a few technical and analytical challenges constraining its full exploitation in protein interaction studies: (I) A significant challenge is the low abundance of cross-linked (XL) products in samples. At concentrations orders of magnitude lower than non-cross-linked species, XL-products demand sensitive MS techniques with a high dynamic range for detection and quantitation in a background dominated by more abundant species and often requiring XLs specific enrichment strategies.^1,2,5^ (II) Another challenge in XL-MS is the interpretation of complex fragment ion spectra generated XL-products thatoften present overlapping series of fragment ions and, in the case of MS-cleavable cross-linkers, diagnostic fragment ions generated by the cross-linker. As a result, the unambiguous identification of cross-links necessitates specialized analytical strategies and computational data anaylsis tools.(III) The ambiguity of XL site adds an additional layer of complexity, since the presence of multiple potential XL sites introduces uncertainties in determining the precise interaction interfaces.^6^ (IV) Datadependent acquisition (DDA) methods, while commonly used in XL-MS, exhibit limitations in effectively sampling the peptide matrix.^7^ In DDA, mass spectrometers select precursors for fragmentation based on their intensity, leading to a bias towards more abundant species. Consequently, XL-products that are typically present in the low-intensity range might be overlooked, resulting in a potentially significant underrepresentation of important protein interaction events.

To counter these limitations, targeted and sensitive data acquisition methods, such as parallel reaction monitoring (PRM), have been implemented where all fragment ions from a limited list of known XL products are monitored.^8^ More recently, data-independent acquisition (DIA) techniques have also been explored as an alternative. DIA facilitates systematic and unbiased sampling across the entire mass range, improving the coverage of low-intensity XLs. Its unbiased nature enhances both sensitivity and reproducibility, crucial for detecting low-abundance species and transient interactions. However, XL MS2 (or product ion) spectra are chimeric themselves which challenges the integration of DIA into XL-MS. DDA spectral libraries remain necessary for accurate XL identification. Recently, aided by DDA spectral libraries, it was shown that DIA can boost the sensitivity and reproducibility of detection of XLs.^9,10^

With the implementation of Parallel Accumulation-Serial Fragmentation (PASEF)^11^ on timsTOF instruments (Bruker Daltonics) the sampling efficiency of the available ions has been improved significantly, resulting in several successful implementations of PASEF for XL-MS analysis.^12,13^ However, the inherent limitations of biased sampling in DDA persists, necessitating critical validation of raw MS data with visualization tools like Skyline.^14^ Despite these advancements, the integration of techniques, such as trapped ion mobility spectrometry (TIMS), with DDA-PASEF introduces additional complexity in the data recorded. This complexity often results in limited software support for native DDA-PASEF data, posing a barrier to the widespread adoption of these techniques in cross-linking MS studies.

This technical note details a workflow to process DDAPASEF data using the native Bruker software infrastructure. First, the LC-TIMS-MS/MS data was processed and simplified into peak lists in mascot generic format (.MGF), compatible with most database search engines. Subsequently, these files were used to perform identification of XL-products by MeroX^15^ and coupled with a standardized protein XL-MS document structure (Proxl XML)^16^ to generate spectral libraries for subsequent analysis of the raw MS data with Skyline. This case study, involving bovine serum albumin (BSA) crosslinked with disuccinimidyl dibutyric urea (DSBU) illustrates the processing specifics necessary for XL-MS studies using timsTOF instruments. The methodologies outlined here aim to equip the cross-linking community with the necessary tools to overcome current limitations in DDA applications and create a framework for the quantitative analysis of PRM and DIA XLMS data with the native infrastructure of Skyline which promotes transparency and standardization in data sharing; both critical principles required to enhance the detection, analysis, and understanding of protein interactions.^17^

## MATERIALS AND METHODS

### Cross-linking and digestion

Bovine serum albumin (BSA) cross-linking with DSBU was carried out as previously described.^12^ A 10 μM BSA solution in 50 mM 2-[4-(2-hydroxyethyl)-piperazine-1-yl]ethanesulfonic acid (HEPES) (pH 7.5) underwent cross-linking at room temperature for 1 hour using a 50-fold molar excess of DSBU, freshly dissolved in DMSO at 25 mM. Alongside, a negative control without DSBU was prepared. Post cross-linking, three 80 μg aliquots from each sample were dried using a SpeedVac concentrator. Sample preparation, including protein digestion, was performed using Suspension-Trapping (S-Trap, Protifi) according to the manufacturer’s guidelines.

### LC-TIMS-MS/MS Data Collection

Dried peptides were reconstituted in 320 µL of 30% (v/v) acetonitrile (ACN) with 0.1% (v/v) trifluoroacetic acid (TFA), adjusting to a concentration of 0.25 µg/µL. From this, a 20 µL aliquot (5 µg) was mixed with 250 fmol of Pierce indexed retention time (iRT) standards and further diluted to 200 µL with 0.1% TFA. For analysis, 40 µL (1 µg of BSA-DSBU digest and 50 fmol of iRT peptides) was loaded onto an UltiMate 3000 RSLC nanoHPLC system (Thermo Fisher Scientific), coupled to a timsTOF Pro mass spectrometer (Bruker Daltonics). Peptides were trapped on a C18 precolumn (precolumn Acclaim PepMap 100, 300 μm × 5 mm, 5 μm, 100 Å, Thermo Fisher Scientific) and separated on a self-packed Picofrit (New Objective) nanospray emitter (360 μm OD × 75 μm ID x 150 mm L, 15 μm Tip ID) with C18-stationary phase (3.0 μm, 120 Å, Dr. Maisch GmbH). The precolumn was washed for 15 minutes with 0.1% (v/v) TFA at 30 μl/min and 50°C. Elution and separation were conducted at a flow rate of 300 nL/min using a 90-minute linear gradient of water−ACN (3% to 50% B), where A is 0.1% (v/v) formic acid and B is 0.1% (v/v) formic acid in ACN.

After chromatographic separation, peptides were ionized using electrospray ionization (ESI) at a capillary voltage of 1500 V, with drying facilitated by N_2_ gas at 180 °C and a flow rate of 3.0 L/min. The ions were then analyzed using trapped ion mobility spectrometry (TIMS) in a dual cell setup before tandem mass spectrometry (MS/MS) detection. TIMS-MS/MS data acquisition utilized DDA-PASEF with ion accumulation and ramp time set to 200 ms. Three mobility-dependent collision energy ramps were employed (see Table 1).

**Table 1.**
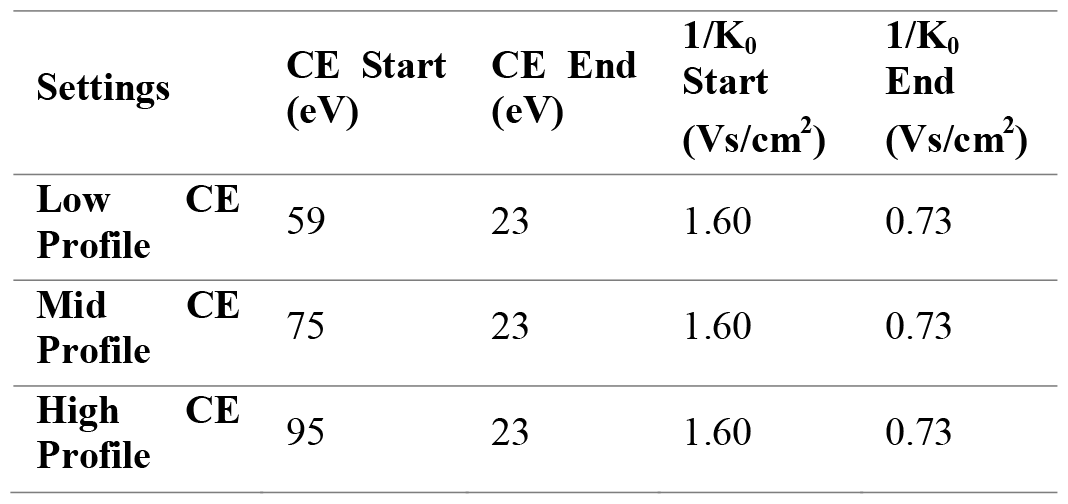
Ion mobility dependent collision energy profiles.

Collision energies were linearly interpolated between specified 1/K_0_ values, remaining constant above or below these values. The PASEF precursor target intensity was set to 100,000, with a minimum intensity threshold of 1,000. Each acquisition cycle, lasting 2.47 s, triggered 10 PASEF MS/MS scans. Precursor ions with m/z ranging from 100 to 1700 and charge states from 3+ to 8+ were chosen for fragmentation. Active exclusion, set for 0.5 min with a mass width of 0.015 Th and 1/K_0_ width of 0.100 V·s·cm^-2^, included early re-targeting if precursor intensity improved by 4x.

### Data Conversion, Cross-linked Peptide Identification and Data Validation

Post-acquisition, DDA-PASEF data was processed using DataAnalysis (v5.3; Bruker Daltonics) to create peak lists of fragment ion spectra in MGF files. Fragment ion spectra with precursor ions collected within a 0.75 min window, having a monoisotopic m/z deviation within 0.015, and 1/K_0_ values within 0.025 V·s·cm^-2^, were combined.

Identification of cross-links was conducted using MeroX (v. 2.0.1.7)^15^ and SwissProt BSA sequence (Uniprot ID: P02769). BSA-DSBU cross-link annotation settings included: fully specific proteolytic cleavage at Lys and Arg (up to 3 missed cleavages, peptide lengths 5-50 amino acids), posttranslational modifications (PTMs) (alkylation of Cys by iodoacetamide: fixed, oxidation of Met: variable), cross-linker specificity (Lys, Ser, Thr, Tyr, N-terminus), XL-fragments (essential: Bu +C_4_H_7_NO, BuUr +C_5_H_5_NO_2_; optional: Δmass = 0), search algorithm: RISEUP mode: up to two missing ions, a-, b-, y-ion series, precursor mass accuracy (10 ppm), fragment ion mass accuracy (15 ppm), 10% intensity prescore cutoff, 25% false discovery rate (FDR) cut-off, and minimum score cut-off of 15. After searches were done, files were combined and the global FDR was set to 5%.

MeroX results from each analysis were merged into a single dataset and exported as .csv files. An in-house R script was used to extract a peptide and precursor-specific ion mobility library from these grouped results. See R-notebook deposited in https://panoramaweb.org/XL-MS_MeroX_Skyline.url and Supporting Information. For each matched precursor ion, mean 1/K_0_ and range were calculated based on all reported 1/K_0_ values from the cross-link spectra matches (XSMs).

These data were formatted into a Skyline ion mobility spectral date to report cross-linking results in ProXL format, a standlibrary and also exported as .csv files.

### Peptide ID Import and Validation in Skyline

MeroX results, converted to Proxl XML files (https://github.com/yeastrc/proxl-import-merox), along with MGF files, were used to create spectral libraries of XL products in Skyline. Pierce iRT standards, measured separately, provided retention time calibration for the BSA-DSBU + Pierce iRT peptide analysis. The compiled ion mobility library was integrated into the document for extracting ion chromatograms (EICs) of the first three isotopes of each precursor ion from raw DDA-PASEF files, using 10 ppm extraction windows. XLs meeting criteria – 5% FDR at the XSM level, detection across all sample replicates, and absence in negative controls – were confirmed as true. Their retention times, adjusted using Pierce iRT standards in Skyline’s retention time calculator, were then indexed. Raw data are publicly available with ProteomeXchange identifier PXD047307.

## RESULTS AND DISCUSSION

### Data Analysis Workflow

This workflow utilizes DataAnalysis (Bruker Daltonics) tools to process native LCTIMS-MS/MS data produced by timsTOF instruments (Figure 1). It begins with mass recalibration and compound detection, associating each compound with fragment ion spectra for a specific *m/z* within a set retention time range and *m/z* error tolerance. Mass spectra are summed and peak picked to derive average precursor and fragment ion spectra. Additionally, for PASEF data, the average precursor scan is used to determine the ion’s charge state, monoisotopic peak *m/z*, and the ion mobility center (1/K_0_). The latter is done by creating an ion mobilogram across the same retention time range. Fragment ion TOF scans across PASEF ramps, falling within the selected *m/z*, retention time, and 1/K_0_ ranges are associated accordingly. The resulting summed and peak picked fragment ion spectra are exported as a simplified MGF file where the precursor ion’s 1/K_0_ is annotated in the scan title. By leveraging Bruker software’s native capabilities, this approach efficiently processes TIMS-MS/MS data, exploiting improvements in sensitivity and selectivity using PASEF, and circumvents limitations in other software tools like MeroX that cannot process raw LC-IMS-MS/MS data. Although demonstrated for XL-MS, this portion of the workflow discussed here is broadly applicable to proteomics, as most DDA database search engines can utilize these MGF files.

**Figure 1.**
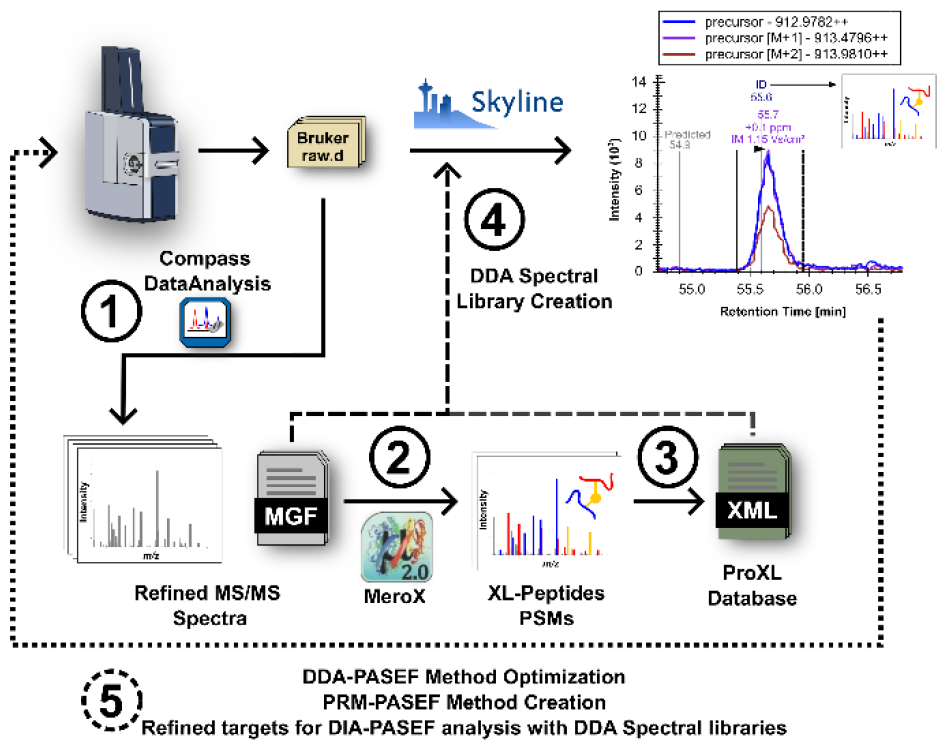
Data analysis workflow. Raw LC-TIMS-MS/MS data was pre-processed with DataAnalysis to create peak lists in MGF format. The MGF files were used as input for crosslinked peptides identification with MeroX 2.0. MeroX results were converted into ProXL format. The ProXL files and MGF files were used as input for creating spectral libraries with Skyline.

Skyline’s support for XLs has significantly advanced since previous XL-MS implementations.^7,8^ Extracting peptide ion mobility libraries from MeroX results now enables streamlined analysis of native DDA-PASEF and DIA-PASEF data in Skyline. Previously, XLs were imported into Skyline as artificial linearized entities with mass shifts from cross-linkers represented by artificial modifications or use of the “molecule” interface of the software. These methods, however, led to ambiguity in cross-linking site reporting and limited use of Skyline’s native peptide fragment ion calculation infrastructure. Recent community discussions have highlighted the need for standardization in XL-MS reporting.^17^ Therefore, Skyline has undergone several improvements to align with these standardization goals. Enhancements include better native support for calculating cross-linked peptide fragment ions, including those from MS-cleavable cross-linkers, and a mandate to report cross-linking results in ProXL format, a standardized XML format that already supports multiple XL –MS database search engines.

MGF files are utilized for XLs identification using MeroX, which are then parsed into ProXL XML files. These ProXL files guide the BiblioSpec function in Skyline to map MGFstored fragment ion spectra to XL proposals, creating a crosslinked peptides spectral library. This library, along with indexed retention times (iRTs) facilitates analysis of XL -MS data with targeted and untargeted workflows that have al eady been developed for other bottom-up proteomics applications. Accompanying this technical note is a detailed tutorial on applying this workflow to a well-characterized XL-MS benchmark system^18^, BSA (see Supporting Information or https://panoramaweb.org/XL-MS_MeroX_Skyline.url). While compatible with all LC-MS/MS platforms, additional information is provided for processing data from tims TOF instruments.

### Beyond Spectrum-Centric Validation in XL-MS

Traditionally, XL-MS has employed a ‘spectrum-centric’ approach^19^, where peptide identification relies on a reliable XL spectrum match (XSM). With this approach only peptide sequences that are derived from a match to a fragment ion spectrum are considered to be present in a particular sample. When replicates are available, it has been shown that IDs shared in multiple replicates tend to be the most reliable and can control false discovery rate (FDR) efficiently^20^. However, this raises questions about the reliability of XSMs that, despite passing global FDR filtering, are found only in some of the technical replicates.^15^ When considering the incomplete sampling limitations of DDA, it is incorrect to assume that a peptide is not present in a sample if no XSM exists. The corresponding precursor ions, although present, might simply not have been selected for fragmentation. This consideration is particularly relevant for low abundance signals, such as those of XLs, amidst intense background. Therefore, the XL-MS community has developed further validation approaches that leverage available protein models to use cross-link distance constraints as false positive discriminators^21,22^. Although this approach proves its usefulness for large proteome-wide datasets, it is limited by the availability and accuracy of protein models and is ineffective for intrinsically disordered proteins or regions with flexible conformation ensembles that defy standard modeling and distance analysis^23^.

This technical note proposes a complementary validation approach for XLs identified with data acquired in DDA mode, adopting strategies from ‘peptide-centric’ methods^19^ used in DIA and targeted data analysis. Although the list of XLs that are tested for detection still depends on spectrum-centric search engines such as MeroX, here we show how to use the complete LC-(TIMS)-MS/MS data collected to test detection and alignment of identifications proposed from any replicate measurement. This approach assumes each peptide elutes once during its chromatographic separation, collecting precursor and fragment ion spectra from which a diversity of data can be inferred apex retention time, peak shape, co-elution of isotopes, precursor ion’s mass error, isotopic pattern, ion mobility, and XSMs. All of this information is weighed together to make a decision on whether the peptide was detected or not either through automatic chromatographic peak peaking models or manual evaluation. For this purpose, Skyline has been chosen due to its versatility in visualizing LC-TIMS-MS/MS data and promotion of transparency in data analysis by enabling sharing results to public databases like Panorama^24^.

This paradigm shift is particularly useful in simplified in-vitro systems with limited cross-link diversity for accurate FDR measurement at the peptide and inter-protein levels. It also compensates for the stochastic nature of DDA while at the same time allows aligning results acquired with different instrument settings or acquisition modes. An example is illustrated in Figure 2, where an UpSet plot^25^ is used to contrast IDs from four technical replicates, measured with a low collision energy profile (Low CE), with a list of validated precursor ions of XLsdes. This analysis reveals that while only 42% of identified precursor ions carried XSMs across all technical replicates (Figure 2; top panel), EICs of peptides with XSMs in a single replicate still showed matching chromatographic peaks, ion mobility, and retention time. This is exemplified with the [M+3H]^3+^ ion of the KQTALVELLK-KFWGK(DSBU@1,1) XL (Figure 2; middle panel). Notably, the EICs of this ion were distinguishable despite their low signal-to-noise (S/N) ratio. Only, through the summation of multiple TOF scans across the PASEF ramps collected across this chromatographic peak an interpretable fragment ion spectrum was still available for one of the replicates (Figure 2; bottom panel).

**Figure 2.**
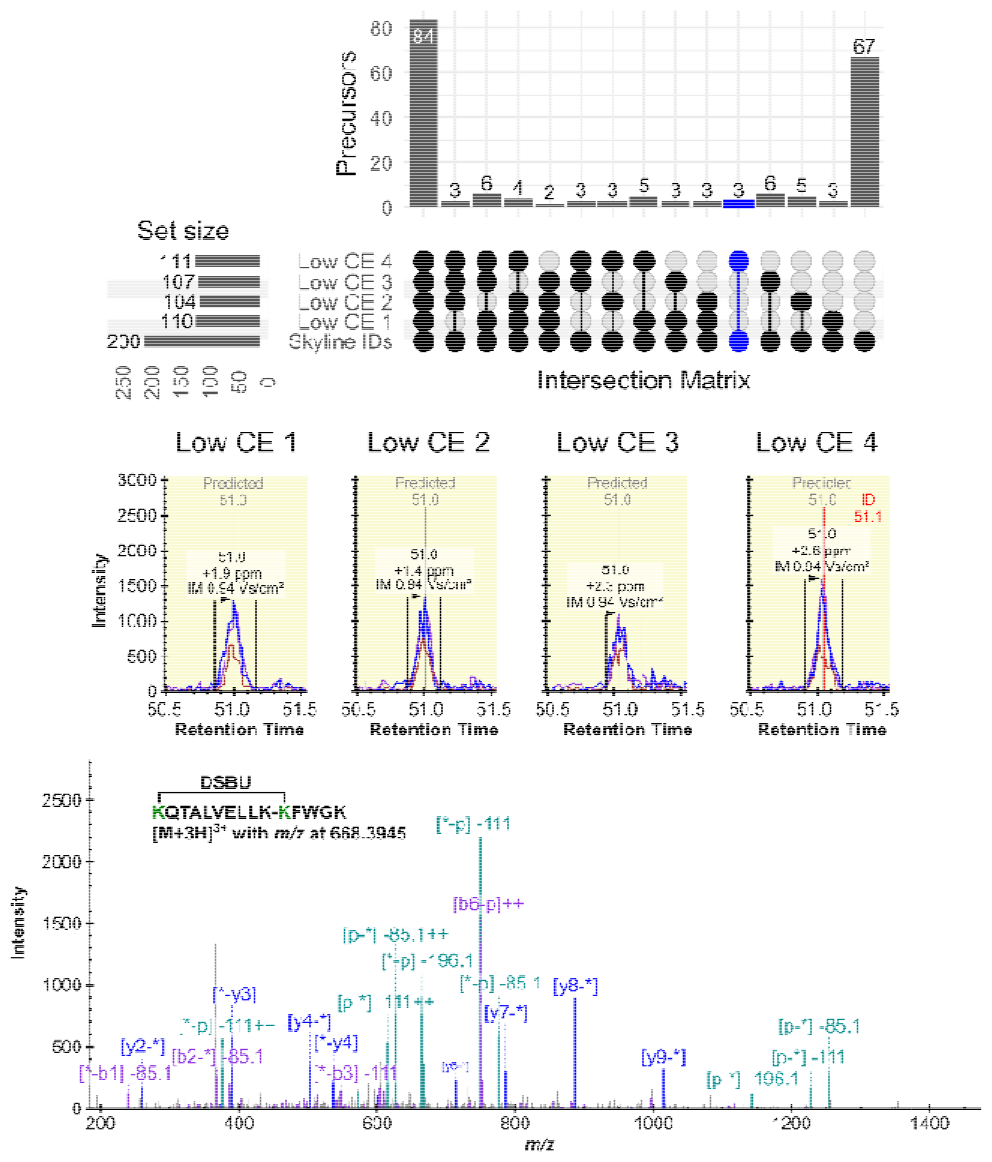
Evaluation of identification reproducibility of precursor ions associated to. UpSet plot representing the overlap of identified precursor ions (top panel) across replicate injections of a pooled digest of cross-linked BSA with DSBU. The identified precursors per replicate are contrasted with a list validated using results from all measurements in this dataset. Blue highlight corresponds to the precursor ions carrying XSMs only in “Low CE 4” sample. Extracted ion chromatograms of the [M+3H]^3+^ ion of the KQTALVELLK-KFWGK(DSBU@1,1) XL (middle panel) which was identified by MeroX only in replicate “Low CE 4” and corresponding XSM (bottom panel).

Additionally, the UpSet plot reveals that a subset of 67 precursor ions, not identified in the four technical replicates, were identified exclusively in samples subjected to Low CE replicates. These ions required increased collision energies for effective fragmentation, generating interpretable spectra. In contrast, ions identified with the Low CE profile often resulted in low-scoring or over-fragmented spectra in measurements done with higher collision energy settings. These results highlight the benefit of characterizing samples with distinct collision energy profiles to expand the coverage of identifiable peptides in a sample. Although exposure of precursor ions to distinct collision energy profiles was required to achieve ideal fragmentation, by design this approach does not produce good XSMs in all replicates. Here is where careful matching between runs is facilitated by the consistent detection of precursor ion data and their retention time, indexed against iRT standards. Looking forward, implementing DDA gas phase fractionation^26^ or off-line fractionation^27^ could be used to enhance spectral library comprehensiveness with the workflow proposed here.

Moreover, by using the iRTs of XLs it is possible to capitalize on the typically low signal yields of cross-linking reactions, utilizing them as indicators of authentic cross-link presence. It involves contrasting the iRT of XLs in negative controls where no cross-linker has been added to ensure the cros -link ID is not assigned to other matrix signals. This method is especially valuable for discerning unique, low-intensity species linked to XSMs that were marginally accepted by i entification software. Once confirmed as unique to the crosslinking reaction, these ambiguous, low-intensity ions c n be prioritized for re-analysis using targeted methods. While not addressed in this study, this approach could also be adapted for entirely untargeted analyses to detect unique cross-linking chromatographic features, with ID assignment as a subsequent step.

For example, in the study of the Y_2_R-NPY system, the peptide-centric method provided a reliable means to detect crosslink sites on the N-terminus of Y_2_R. These sites belong to a region predicted to be intrinsically disordered, showcasing the ability of this approach to reveal interactions that have been intractable to other structural proteomics techniques.

Despite the limitations of DDA, it still heavily used in XL-MS due to the clearer origin determination of fragment ions which simplifies the interpretation of the inherently complex fragmentation pattern of XLs. With this technical note and the accompanying tutorial material (Supporting Information) we show how to leverage some of these limitation by matching identifications between LC-MS/MS runs, a common practice in other proteomics fields^28^. However, we present it in a platform that facilitates visual critical analysis of mass spectrometry data to avoid false transfer of IDs and encourage a shift towards peptide centric evaluation of XL-MS data.

### Framework for DIA-XL-MS Data Analysis in Skyline and Integrative Data Processing

Recent workflows have shown the processing of XL-MS results acquired with DIA using DDA spectral libraries for a peptide-centric data analysis^9,10^. The presented workflow leverages Skyline’s visualization tools for DIA data interrogation. As a proof of concept, a Skyline document examined DIA-PASEF data from a separately prepared BSA-DSBU digest, measured weeks apart. This dataset can be found in: https://panoramaweb.org/XLMS_MeroX_Skyline.url. The use of validated iRTs from DDA measurements was key to aligning results, accommodating retention time shifts from changes to the LC system, i.e., change of trap and analytical column. iRTs’ role in dataset alignment tests uniformity across different sample preparations, addressing challenges for reliable quantitative XL-MS studies^1^.

DIA data’s promise for enhancing reproducibility and peak detection selectivity, particularly through peptide-specific fragment ions, is juxtaposed with the challenge of identifying low-intensity XLs when fragment ions are the sole focus. DDA remains indispensable for generating spectral libraries to interrogate DIA data, a process streamlined by Skyline’s vendor-neutral platform and its compatibility with diverse data sources once formatted to ProXL. This integrative approach, strengthened by recent validation as an effective spectrum-centric filter for cross-links and evidenced in studies like Matzinger et al., 2022^20^, enhances the robustness and authenticity of identified cross-links. The comprehensive workflow presented, utilizing accessible tools such as DataAnalysis, MeroX, ProXL, and Skyline, represents a paradigm shift in the cross-linking community’s data analysis methodologies. It not only demonstrates the collaborative potential of instrument-vendors and open-source platforms but also sets a replicable and reliable course for unveiling the complex interactions within the proteome, underscoring the collective advancement towards more nuanced proteomic exploration.

## CONCLUSION

Advancements in XL-MS, driven by peptide-centric validation and interrogation of quantitative queries, signify progress in analyzing protein interactions. The integration of PASEF for DDA and DIA pipelines sets a new standard for identifying and quantifying cross-links. Despite the challenges in DIAPASEF data analysis for XLs, this technical note outlines effective strategies for overcoming these obstacles. The creation of comprehensive DDA spectral libraries from multiple search engines in Skyline and empirical retention time indexing are crucial in this dynamic field, ensuring rigorous and innovative analysis.

## Supporting information

Tutorial Workflow

## ASSOCIATED CONTENT

### Supporting Information

The Supporting Information contains a tutorial entitled “Tutorial: Cross-linking mass spectrometry with timsTOF DDA-PASEF using MeroX and Skyline” (PDF).

## AUTHOR INFORMATION

**Authors**

**Frank Hause** – Department of Pharmaceutical Chemistry & Bioanalytics, Institute of Pharmacy, Martin Luther University Halle-Wittenberg, D-06120 Halle (Saale), Germany; Center for Structural Mass Spectrometry, D-06120 Halle (Saale), Germany. Institute for Molecular Medicine, D-06120 Halle (Saale), Germany.

Claudio Iacobucci – Department of Pharmaceutical Chemistry & Bioanalytics, Institute of Pharmacy, Martin Luther University Halle-Wittenberg, D-06120 Halle (Saale), Germany; Center for Structural Mass Spectrometry, D-06120 Halle (Saale), Germany; Institute for Molecular Medicine, D-06120 Halle (Saale), Germany; 4Department of Physical and Chemical Sciences, University of L’Aquila, I-67100 L’Aquila, Italy.

Christian Ihling - Department of Pharmaceutical Chemistry & Bioanalytics, Institute of Pharmacy, Martin Luther University Halle-Wittenberg, D-06120 Halle (Saale), Germany; Center for Structural Mass Spectrometry, D-06120 Halle (Saale), Germany.

**Dirk Tänzler** – Department of Pharmaceutical Chemistry & Bioanalytics, Institute of Pharmacy, Martin Luther University Halle-Wittenberg, D-06120 Halle (Saale), Germany; Center for Structural Mass Spectrometry, D-06120 Halle (Saale), Germany.

**Nicholas Shulman** – Department of Genome Sciences, University of Washington, Seattle, United States of America

**Michael Riffle** – Department of Biochemistry, University of Washington, United States of America

**Brendan MacLean** – Department of Genome Sciences, University of Washington, Seattle, United States of America

## Author Contributions

All authors have given approval to the final version of the manuscript.

## Notes

The authors declare no competing financial interest. Raw data are publicly available with ProteomeXchange identifier PXD047307.

## ACKNOWLEDGMENT

JCRE was funded by the DFG (CRC 1423, project number 421152132); AS acknowledges financial support by the DFG (RTG 2467, project number 391498659 “Intrinsically Disordered Proteins-Molecular Principles, Cellular Functions, and Diseases”, INST 271/404-1 FUGG, INST 271/405-1 FUGG), the Federal Ministry for Economic Affairs and Energy (BMWi, ZIM project KK5096401SK0), the region of Saxony-Anhalt, and the Martin Luther University Halle-Wittenberg (Center for Structural Mass Spectrometry). MR acknowledges financial support by the University of Washington’s Proteomics Resource (UWPR95794). BM acknowledges funding by NIGMS National and Regional Resources (R24) Program. The authors thank Dr. Lindsay Pino, and Alessio Di Ianni for their time and constructive input invested into this work.

