## Supplementary material for "A Workflow for Improved Analysis of Cross-linking Mass Spectrometry Data Integrating Parallel Accumulation-Serial Fragmentation with MeroX and Skyline": Tutorial Workflow

**Tutorial: Cross-linking mass spectrometry with timsTOF DDA-PASEF using MeroX and Skyline**

^1^Department of Pharmaceutical Chemistry and Bioanalytics, Martin-Luther-University Halle-Wittenberg; ^2^Center for Structural Mass Spectrometry, Martin-Luther-University Halle-Wittenberg; ^3^Institute for Molecular Medicine, Martin-Luther-University Halle-Wittenberg; ^4^Department of Physical and Chemical Sciences, University of L’Aquila, Via Vetoio, Coppito 1, 67100 L’Aquila, Italy;^5^Department of Genome Sciences, University of Washington, Seattle, WA; ^6^Department of Biochemistry, University of Washington, Seattle

This tutorial explains the steps posterior to mass spectrometry data acquisition on a timsTOF Pro instrument of a digest containing cross-linked peptides. The current material refers only to the analysis of DDA data acquired with Parallel Accumulation-Serial Fragmentation (PASEF). The digest consists of bovine serum albumin (BSA) cross-linked with disuccinimidyl dibutyric urea (DSBU) at a ratio of 50-to-1 with respect to cross-linker to protein concentration at room temperature for 1 hr. Digestion was performed using a suspension trapping (S-Trap) method following the ProtiFi (manufacturer) standard operational procedure. This digest has been spiked with Pierce™ Peptide Retention Time Calibration Mixture (ThermoFisher Scientific) and the sample load on the LC-MS system corresponded to 1 µg of BSA (with respect to protein concentration prior to digest) and 50 fmol of the iRT peptides.

The tutorial will be focused on the analysis of 13 DDA-PASEF measurements of the digest mentioned above corresponding to 5 instrument technical replicates from a digest pool with the same collision energy (CE) profile, two digest pool replicates analyzed with different CE profiles, three sample preparation replicates, and three negative control sample preparation replicates. The negative controls correspond to the same batch of BSA used for the cross-linking (XL) experiments that were processed in parallel, but no cross-linker was added to the sample.

All materials for this tutorial can be found in: <https://panoramaweb.org/XL-MS_MeroX_Skyline.url>

The tutorial will use advanced features from Skyline that have dedicated tutorials to them. Although we try to make the tutorial as self-explanatory as possible, we advise users to have a look at the following tutorials:

MS1-filtering: <https://skyline.ms/wiki/home/software/Skyline/page.view?name=tutorial_ms1_filtering>

iRT tutorial:

<https://skyline.ms/wiki/home/software/Skyline/page.view?name=tutorial_irt>

Ion mobility tutorial:

<https://skyline.ms/wiki/home/software/Skyline/page.view?name=tutorial_ims>

Make sure to download Skyline (<https://skyline.ms/project/home/software/Skyline/begin.view>) and MeroX v2.0.1.7^1^. You can find the most recent version of MeroX in the “Supplementary Files: MeroX Software” folder within the Panorama submission referenced above.

### Processing and binning MS/MS scans with Bruker Compass DataAnalysis

PASEF allows for Multiple MS/MS scans from distinct precursor ions to be obtained within each trapped ion mobility scan (i.e. ramp). These MS/MS scans correspond to the inverse reduced ion mobility (1/K_0_) range of the detected precursor during the prior precursor scanning PASEF ramp. The same precursors are re-selected for fragmentation in subsequent PASEF ramps until a certain target intensity criteria (defined in the acquisition method) is satisfied before placing the precursor in a dynamic exclusion list. Additionally, the precursor ions can be removed early from the dynamic exclusion list if these are detected with an improved signal-to-noise (S/N) ratio.

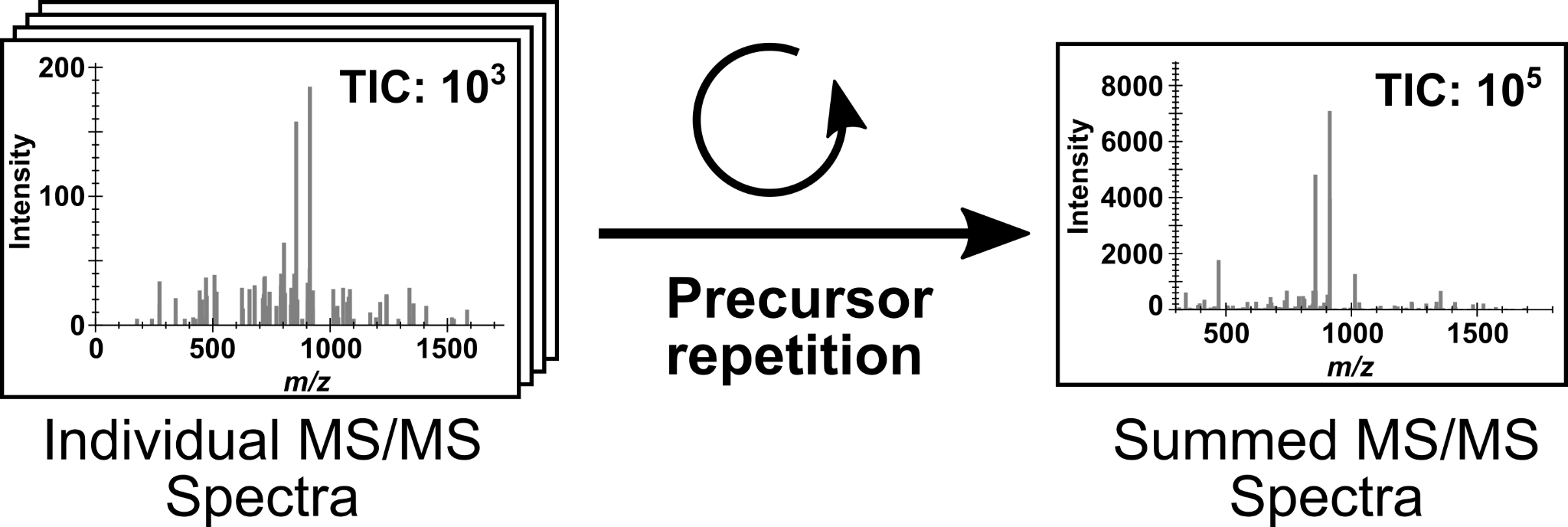

Summing up fragmentation spectra results in improved S/N of true fragment ions. However, this process requires correct peak picking in the retention time (t_R_) and 1/K_0_ dimensions to associate correctly only the fragment ions generated from their corresponding precursor ions. For the data of this tutorial, Bruker’s *Compass DataAnalysis* (provided with all Bruker instruments) was used to summarize all the relevant fragment ion scans into a single summed spectrum and written into a mascot generic format (*.mgf*) peak list. Each entry within the *.mgf* file corresponds to a “compound”, that is a precursor ion with a defined retention time, inverse ion mobility, and quadrupole isolation range.

#### Method parameters

The precision with which DataAnalysis can identify scans associated with a specific precursor depends on several settings for the PASEF peak picking. To access these settings in DataAnalysis do the following:

- Open Bruker Compass DataAnalysis
- Click on **Parameters…** within the **Find** drop-down menu
- Under the **Find** menu click on **PASEF**

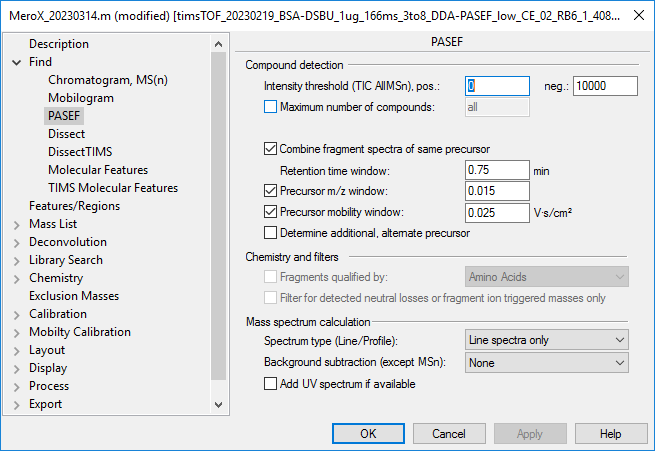

The settings displayed here are saved within the method file used to process the raw files. The crucial settings to pay attention are:

- **Combine fragment spectra of same precursor** [Ticked]
- **Retention time window**
- **Precursor *m/z* window** [Ticked]
- **Precursor mobility window** [Ticked]

The correct settings will depend heavily on the chromatographic and MS settings used during acquisition. Using settings with overly wide ranges might result in binning together closely eluting isobaric species or isomeric species that are resolved in the combined LC-IMS dimensions. While using settings with too small ranges will result in signal splitting. It is advised to visualize and profile initial results acquired with default settings (all of the above unticked) on a quality control sample or more easily detectable peptides (e.g., unmodified peptides in a digest containing cross-linked peptides). For the purpose of this tutorial we will use the following settings that describe well the cross-linked peptides within the samples analyzed:

- **Combine fragment spectra of same precursor** [Ticked]
- **Retention time window:** 0.75 min
- **Precursor *m/z* window** [Ticked]: 0.015
- **Precursor mobility window** [Ticked]: 0.025 V∙s / cm^2^
- Click **OK** when parameters have been filled

NOTE: at the end of this tutorial you will see some of the analysis performed to check if these initial parameters were appropriate for the current dataset.

#### (Optional) Internal mass recalibration

The mass accuracy of the data acquired can be improved by updating the calibration equation of each measured file to account for small drifts in *m/z* estimation caused by environment changes (e.g. temperature, humidity changes). To perform internal mass recalibration it is necessary that the calibration solution signal is still detectable through the chromatographic run. For the data presented here, the volatile ESI-L Low Concentration Tuning Mix (Agilent Technologies) was used for initial instrument calibration and post-acquisition internal mass recalibration. This volatile mixture can still reliably display three abundant ions (with *m/z* at 622.028961, 922.009799, and 1221.990637) after 5 days of placing 20 µL on top of the air filter located on top of the ESI inlet of a timsTOF Pro in the configuration presented below.

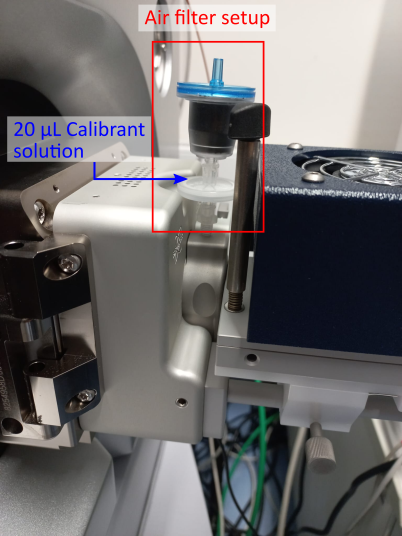

You can either define a “Segment” in your acquisition method at the beginning or end of your LC-MS/MS run or identify a region of your chromatographic run that consistently displays the calibrant signals. These sections will be used by DataAnalysis to perform an “Automatic Internal Calibration”. To define these settings do the following:

- Click on **Parameters…** within the **Calibrate** drop-down menu
- Under the **Calibration** menu on the **Internal Calibration**
- For **Calibration group** select **ESI**
- For **Calibration list** select **Tuning Mix ES-TOF CCS 3 Masses (filter)**; if not present define the list as follows:

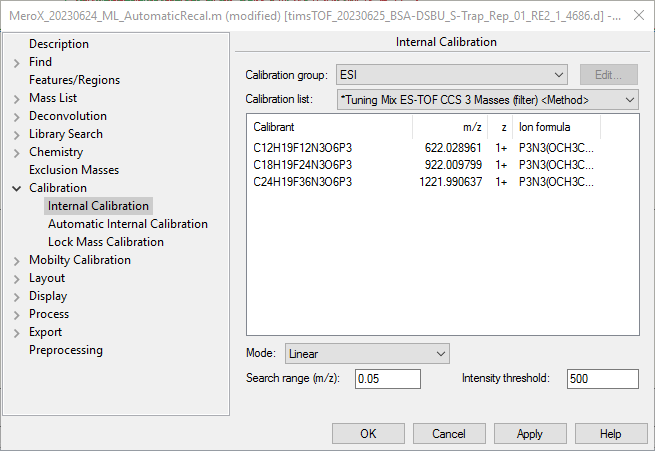

- Set calibration **Mode** as **Linear**
- Error tolerance **Search range (*m/z*): 0.05**
- **Intensity threshold** of **1000** after averaging the calibrant extraction window

Now indicate the chromatogram regions that should be used for detecting the calibrants.

- Within the **Calibration** parameters click on **Automatic Internal Calibration**
- The acquisition method did not have defined calibration “segments” DataAnalysis is directed to look for the calibrants within **Start: 0 min** and **End: 20 min**.
- Additionally, retain the extracted ion chromatograms (EIC) and compounds spectra of the calibrants by ticking **Retain calibration spectrum and chromatogram**
- Click **OK**

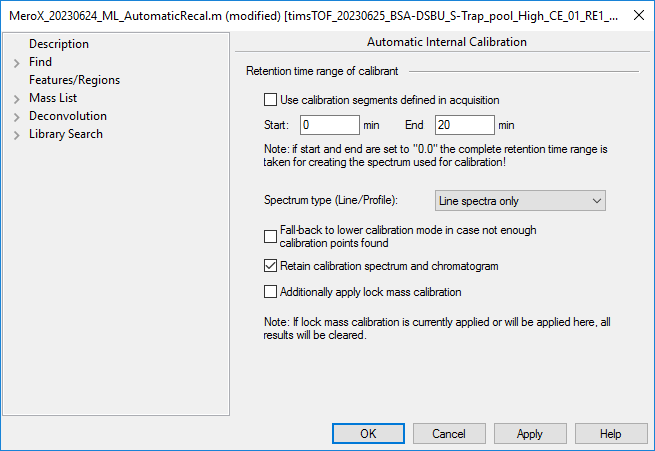

To manually perform the internal calibration within the DataAnalysis graphical user interface (GUI) do the following:

- Open the raw DDA-PASEF files
- Make sure the file of interest is highlighted in the **Analysis List**
- Click on **Automatic Internal** within the **Calibrate** drop-down menu
- Once the file has been processed, make sure to **Save** (Ctrl+S) so the new calibration is retained
  - IMPORTANT: If you would like to keep your files with the original calibration used during acquisition make sure these steps are performed only on a copy of the original file

#### Automatic processing with visual basic script

All processing can be performed automatically using visual basic scripts within DataAnalysis. Bruker method files are made of 1) a set of parameters that DataAnalysis should interpret and 2) the sequence of events with which you want to process the raw files defined in the script. For the analysis of this data, a “default” proteomics processing method provided with the instrument was edited with the parameters described above and the script was edited to ensure that the *.mgf* output was downstream compatible with MeroX^1,2^, proXL^3^ and Skyline^4,5^. You can find the processing script as a text file in the same folder where you can find this tutorial and embedded within the raw Bruker .d folders when opened in DataAnalysis.

To have a look at the script do the following:

- Click on **Script…** within the **Method** drop-down menu or **Ctrl+F2**

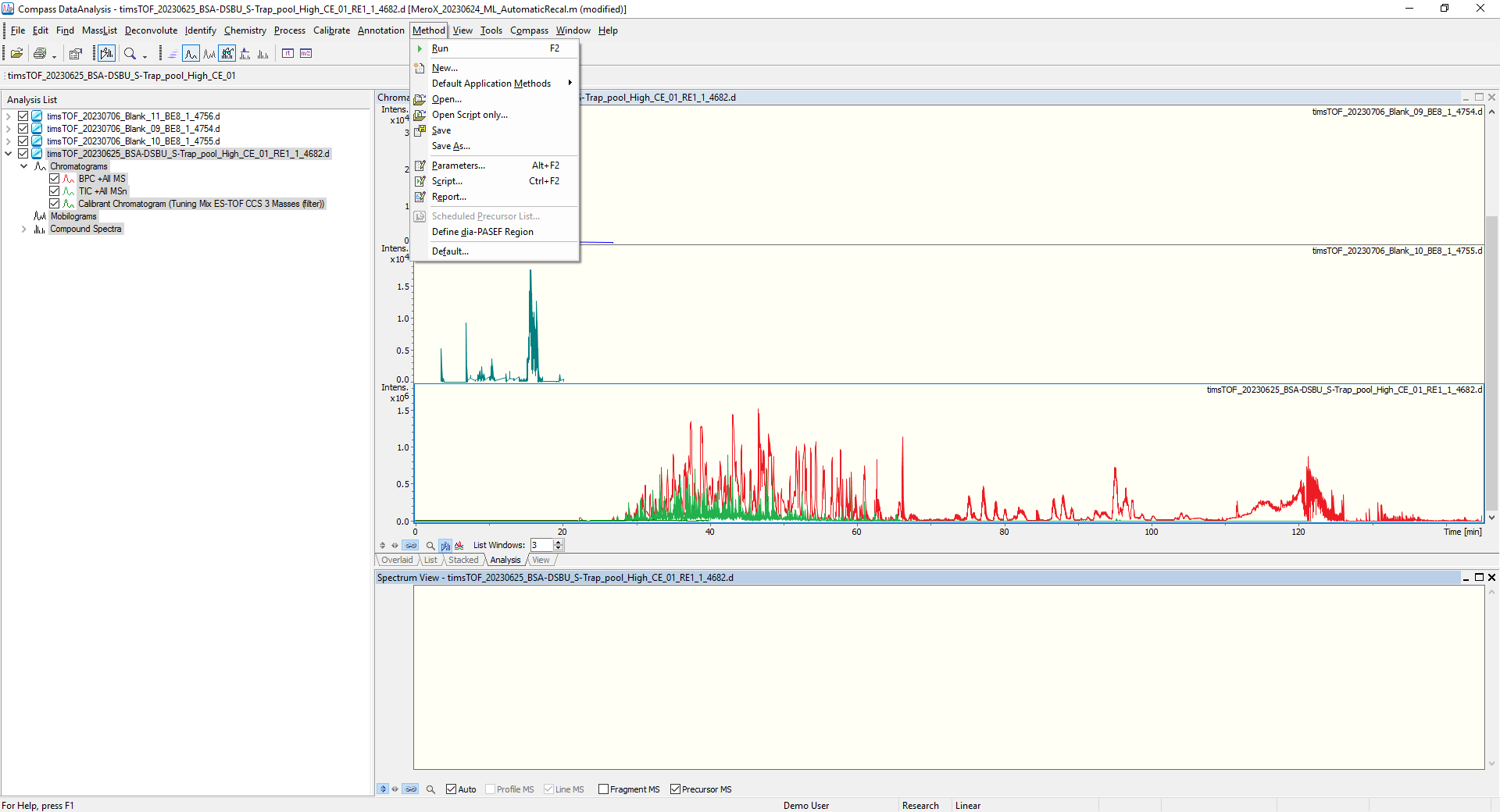

Documentation of the “functions” available for automatic data processing within DataAnalysis can be found in the **Help** (**F1**) manual. The general steps performed by the script are the following:

1) Define working directory for reading and output
2) (Optional) Perform internal calibration
   a. Analysis.RecalibrateAutomatically
   b. By default, the automatic recalibration is commented out of the script. To enable this part of the processing remove the quote mark.

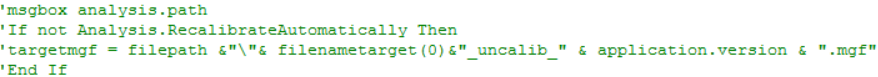

Keep in mind that this should only be done if you have defined correctly the automatic calibration parameters and identified a region that will consistently show the necessary calibration signals. If the internal calibration fails, then output *.mgf* will have a “_uncalib_” text mark appended to the file name.

3) Automatic peak picking and compound generation from PASEF data using the parameters defined above
   a. analysis.processautoMSn
4) Write a *.mgf* file within the data analysis folder (i.e., inside the Bruker .d folder)
   a. The scan title and additional metadata required for correct parsing for downstream analysis is extracted with the remaining part of the script. This native data mining portion of the script was graciously provided by Bruker.

NOTE: when processing data automatically with this script, the generated compound spectra are deleted (“Clear Results”) after processing. Keeping the compound spectra within the analysis document severely impairs the processing capacity of the software. If you are interested in seeing the generated compounds by DataAnalysis we advise to perform the compound creation within the GUI by using the **Compounds-AutoMS(n)/PASEF** function within the **Find** menu. However, we also recommend that you perform this one file at the time to minimize computing strain on the pc/software.

The output of these processing steps, either through the GUI (one file at the time) or with automation right after acquisition through *HyStar* or *AutomationEngine*, are fragmentation spectra with well annotated meta data indicating the raw scans used to build them, the center of ion distribution in the inverse ion mobility (1/K_0_) dimension and the apex retention time of the peak detected by DataAnalysis. Additionally, isotope correction is employed and the written precursor ions correspond to the monoisotopic peak detected after summing the precursor scans around the chromatographic peak and that correspond only to the 1/K_0_ of the selected precursor.

NOTE: The steps above apply generally for creating files compatible with most spectrum-centric peptide database search software. Many of the spectra data processing steps required prior to analyte identification have already been performed by DataAnalysis and should be taken into account when defining your search settings, i.e. precursor m/z correction and isotope correction using scans from the MS profile spectrum, retention time correction to the apex of the precursor’s chromatographic peak, and detection of the center of the ion population in the 1/K_0_ dimension.

### Cross-linked peptides search with MeroX

#### Database search settings

All steps outlined before illustrate how you could perform data preprocessing with your own experiments. This pre-processing has already been performed for the DDA-PASEF runs of this tutorial and the .mgf peak lists can be found in the “Raw Data” tab of the Panorama folder (<https://panoramaweb.org/XL-MS_MeroX_Skyline.url>) inside the “MeroX Results” folder.

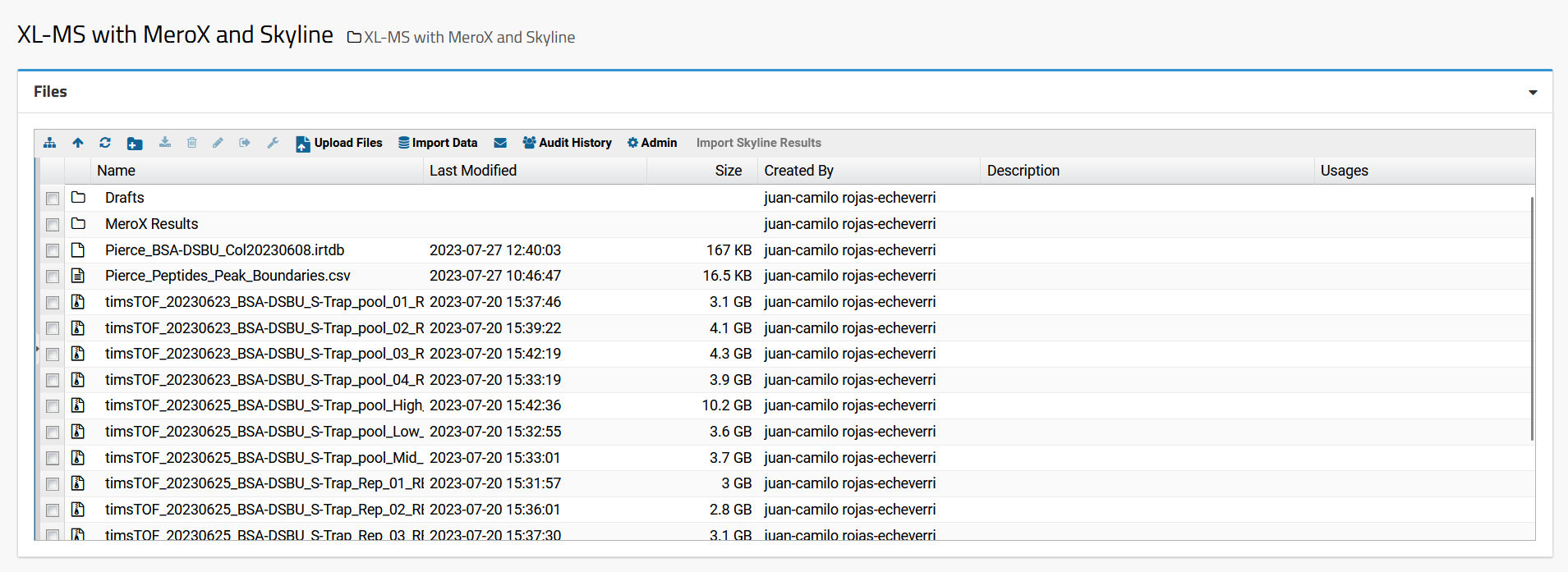

##### Cross-linker definition

The search settings used here are for a single protein (i.e. BSA) cross-linked with DSBU considering reactivity to Lysine (K), Serine (S), Tyrosine (Y), Threonine (T), and peptide N-terminus at both cross-linking positions. Only peptides generated with full trypsin cleavage specificity with up to 3 missed cleavages were considered.

The search settings are embedded within the MeroX results files and this can be extracted using the MeroX GUI and used as a template for editing based on the user’s experimental setup.

- Open MeroX v2.0.1.7.exe windows executable file by double clicking on it
- Click on **Settings**

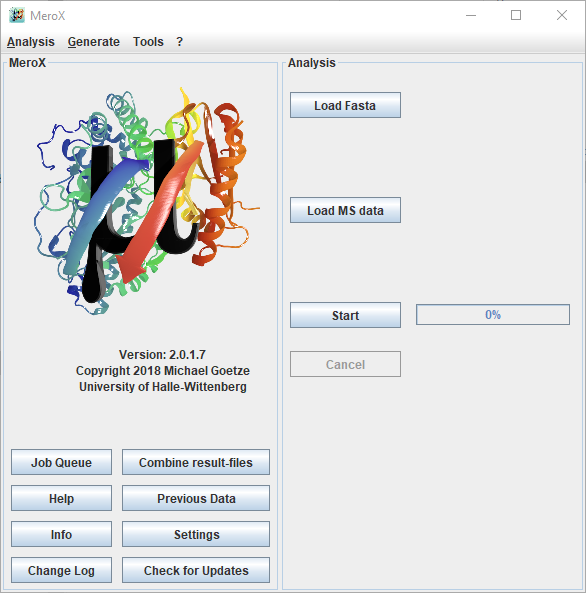

- Then click on **Load** and select a MeroX results file (.zhrm compressed files) that contains the search settings you want to replicate

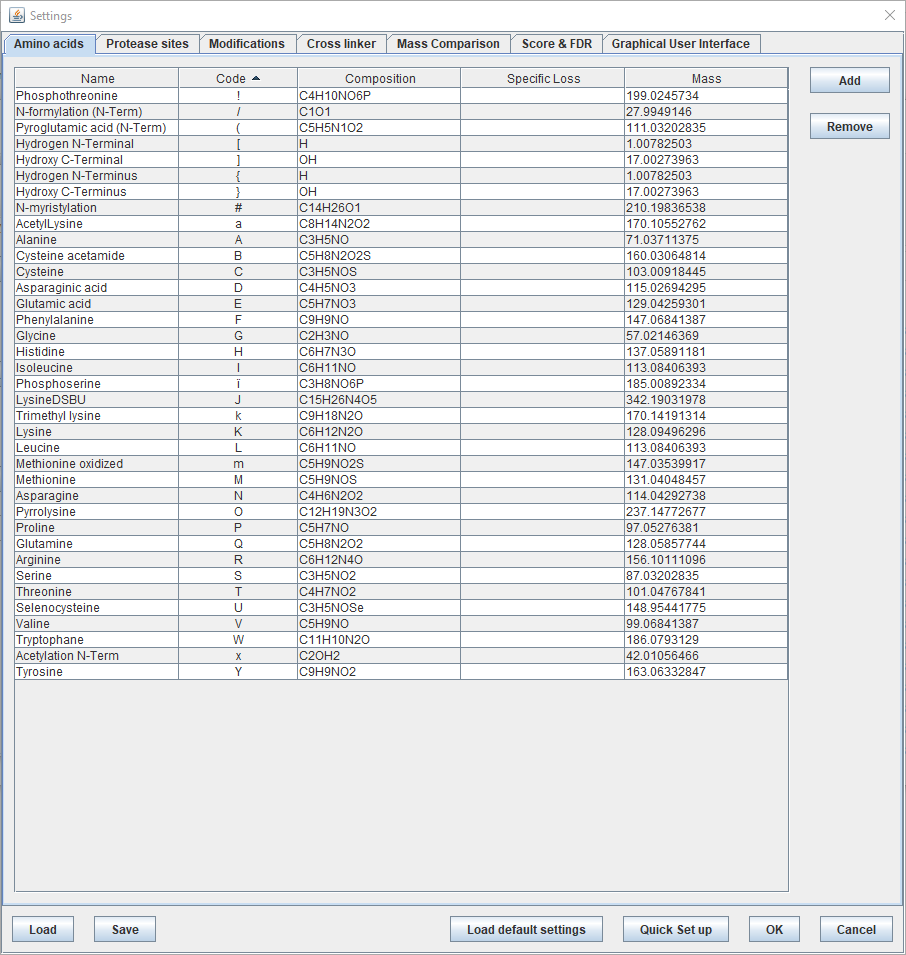

For all the specific settings used for these searches we advise the user to explore all the settings tabs after loading the settings of the search results provided with this tutorial and explore the online documentation in <https://www.stavrox.com/help.htm>. For the sake of this tutorial, we will only focus on a few settings that are relevant for the data pre-processing performed and subsequent data validation. First, let’s have a look at the cross-linker settings:

- On the **Settings** window click on the **Cross linker** tab

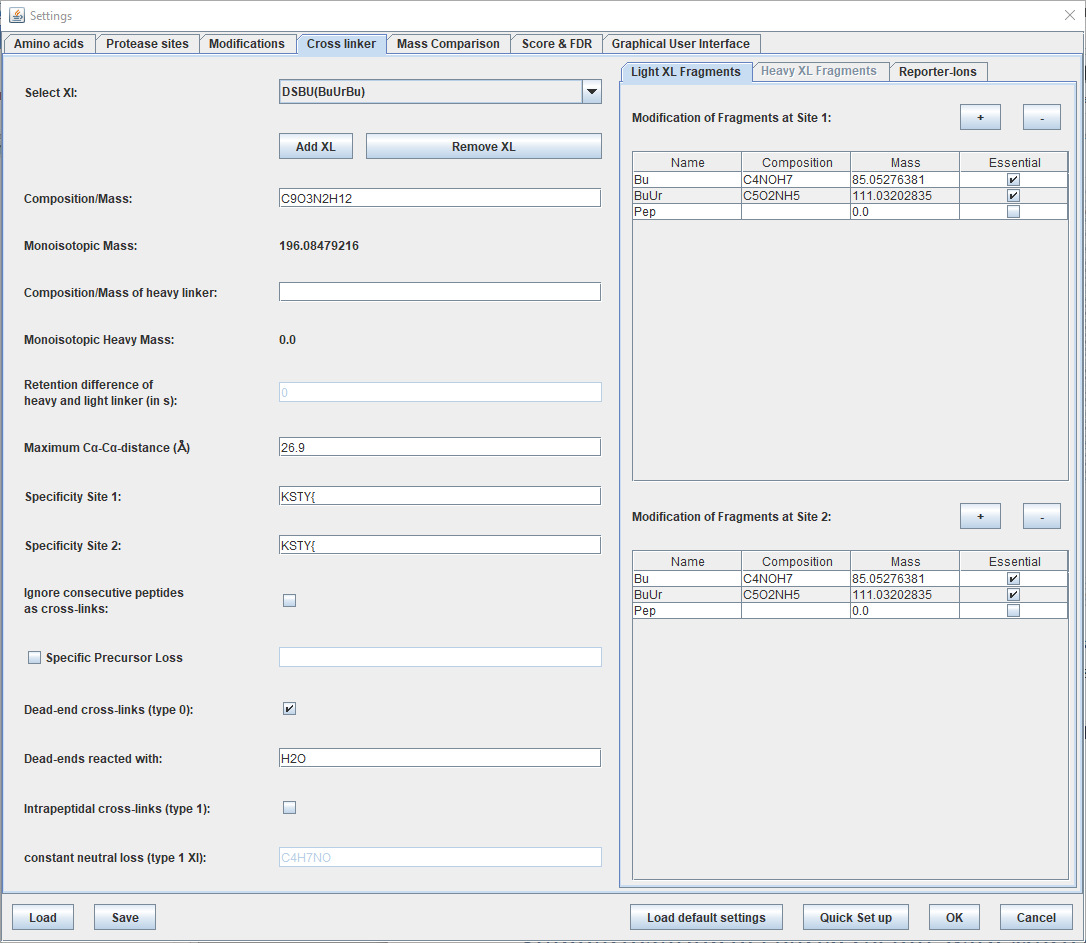

Here the cross-linker composition, specificity, it’s “dead-end” (i.e. DSBU reaction with only one amino acid) products, and the MS-cleavable fragments are defined. The detected precursor ion’s *m/z* have to match a mass increase of +196.08479216 u (C_9_O_3_N_2_H_12_) induced by the reaction product DSBU with two nucleophiles. This mass increase has to also be considered in the fragment ion series for fragments where the two peptide chains remain connected.

NOTE: The fragments induced through beam-type collision induced dissociation (CID) in the timsTOF collision cell correspond to a-, b-, and y-ion series. However, a-ion fragments were only used to support the main sequence coverage provided by b-ions during manual validation.

MS-cleavable cross-linkers such as DSBU, generate additional fragment ions that can be used to further confirm a cross-linked peptide and can also be used to reduce the search space of a specific dataset^6^. Here three types of mass shifts were considered for calculating and matching additional fragment ion series. Ion series containing the mass shifts of +85.05276381 u (**Bu**; C_4_NOH_7_) and +111.03202835 u (**BuUr**; C_5_O_2_NH_5_) correspond to ions retaining only partial fragments of DSBU. When observed, these fragments generate characteristic “doublets” with mass difference of 25.97926454 u that can be used to identify the mass of the individual peptide chains that have been cross-linked.

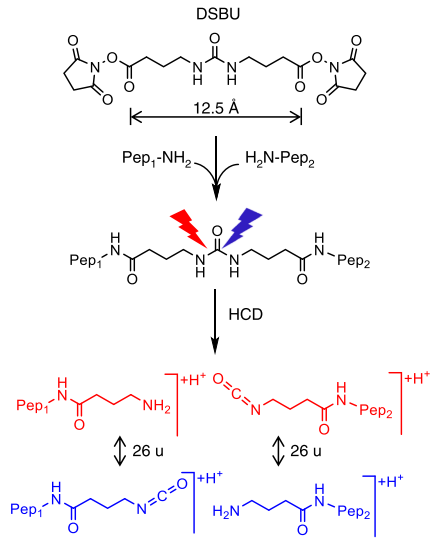

Lastly, the fragment ion series without any mass shift from the canonical mass of each amino acid is considered (**Pep**). These fragments can occur when there is a complete loss of DSBU from the cross-linked site.

By **ticking** the **Essential** box for the **Bu** and **BuUr modifications** only cross-linked peptide proposals that contain these MS-cleavable fragments are considered. On the other hand, the **unticked** setting for **Pep** means that these fragments are calculated and matched, but not used to exclude possible matches.

These mass modifications will be considered again when **defining** the **cross-linker modification** in **Skyline**.

##### Mass accuracy and pre-processing settings

Now let’s have a look at the mass accuracy settings:

- On the **Settings** window click on the **Mass Comparison** tab
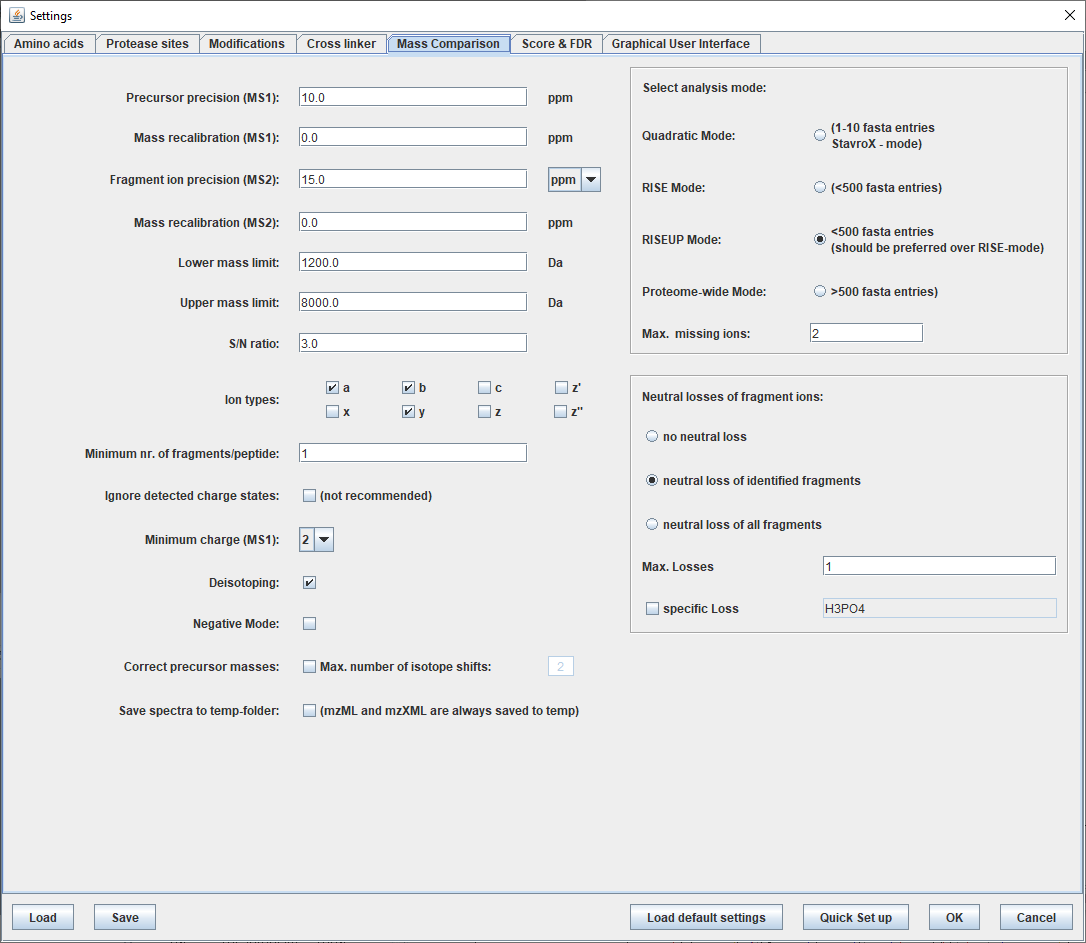

To reduce search space and control false discovery rate the mass accuracy should be reduced as much as the corresponding data set allows. For this we suggest doing initial searches with broad **Precursor precision (MS1)** and **Fragment ion precision (MS2)** and plot the mass error distribution of the proposed ids. The precursor mass error in parts-per-million (ppm) is reported in the MeroX results and the distribution of the data used for this tutorial correspond looks as follows:

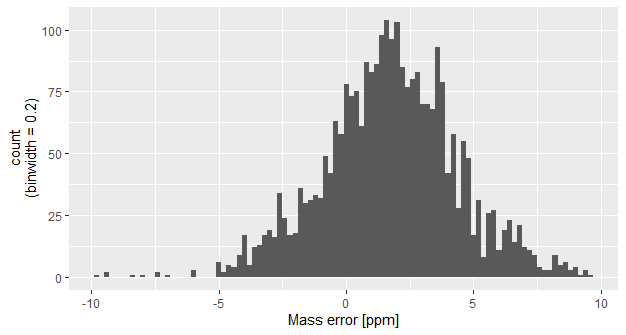

Assuming a normal distribution, 99.7% of matched precursor ions are within 1.75 ±7.9 ppm (i.e. mean ±3 ∙ standard deviation). MeroX allows shifting the deviation with **Mass recalibration** for both precursors and fragment ions, but in order to match MeroX results to the raw data in Skyline this was avoided. Instead, safe error boundaries of **±10 ppm** for **precursor ions** detection while allowing slightly bigger error tolerance of **±15 ppm** for **fragment ions** to extend the detection range of low intensity ions that might have impaired accuracy due to the lower population of ions that are profiled.

If time constraints limit or prevent manual inspection of proposed cross-linked peptides then the minimum number of fragment ions per peptide should be **at least 3**. Here we used a very lenient value of **1**. This choice was done based on the knowledge that manual inspection of cross-linked peptides was performed in downstream analysis in Skyline while integrating the LC and IMS characterization of their respective precursor ions.

DataAnalysis has already performed precursor mass correction which is why the **Correct precursor masses** box has been left **unticked**. However, during this processing, the summed and peak picked fragment ion spectra were not deisotoped. Therefore we still let MeroX perform **Deisotoping** (**ticked**) to reduce spectrum complexity and improve the sensitivity for the monoisotopic fragment ions prior to spectrum interpretation.

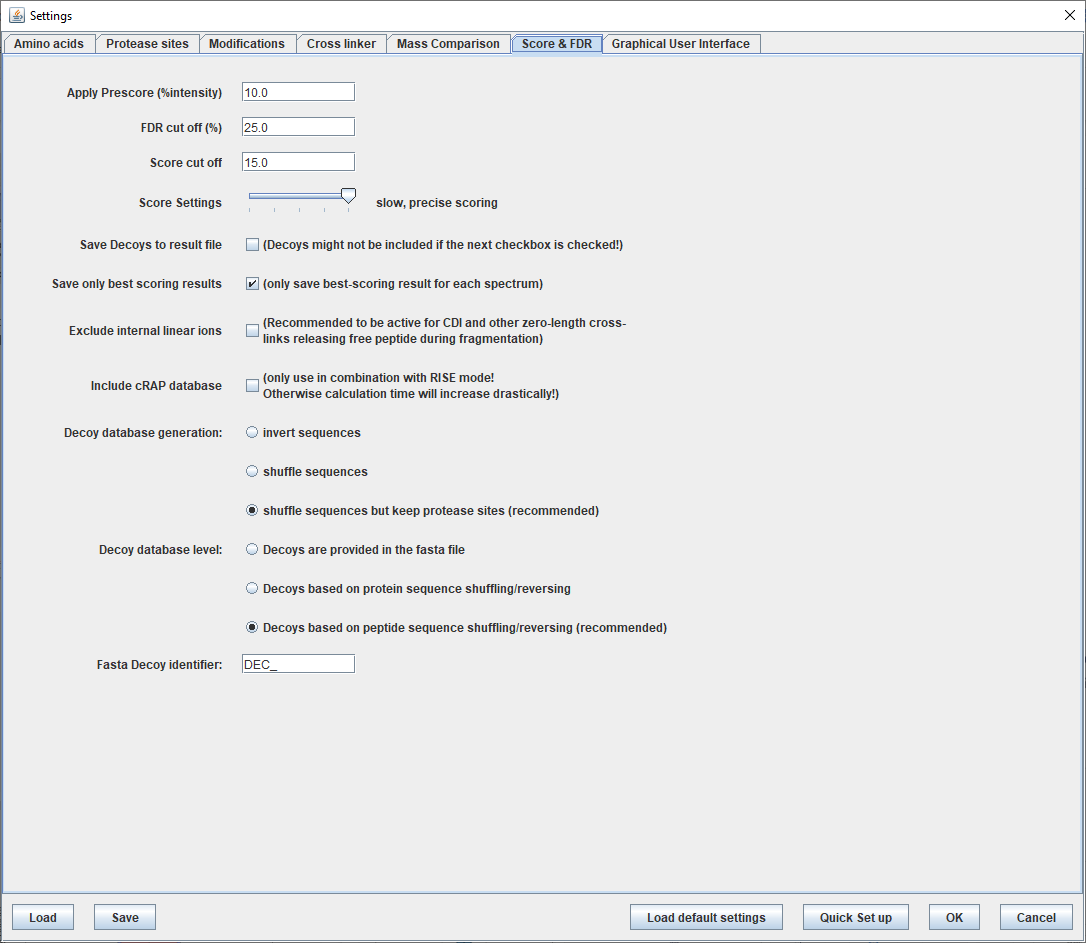

##### Score and false discovery rate settings

MeroX performs some pre-scoring based on the percentage of the total ion intensity explained where spectra matches with many unassigned peaks are filtered out and not considered further by the algorithm. This can be controlled with the **Apply Prescore (%intensity)** and was set at **10%** here.

MeroX allows the user to control the false discovery rate (FDR) estimated using the MeroX composite score explained in detail in previous work^2^. This can also be performed after the database search has been performed. The **FDR cut off (%)** setting here simply limits the results that will be visualized in the MeroX result file. Since manual validation was performed for this dataset we opted on using a very lenient 25% FDR cutoff to explore the results and how the data-preprocessing with DataAnalysis might affect the results. In ideal scenarios, where all fragment ion spectra of a respective ion were binned together, only one peptide spectrum match (PSM) per file is expected. If the dataset contains few unique cross-link peptides, then it is possible that there won’t be enough data points to correctly model the false positive distribution and discriminate it from the true distribution. For single-file results, it might be more useful to rely on the MeroX spectral score. The negative scenario described above can be minimized by determining the MeroX cut-off using the combined results of multiple LC-MS/MS experiments. Here, we allowed MeroX to consider cross-link proposals with a MeroX **Score cut off** of **15** or more.

NOTE: the score cutoffs used here are very lenient because manual annotation of spectra was performed afterwards and, more importantly, because the results were further validated with Skyline that allows considering additional criteria that MS/MS spectrum-centric database search software can perform, i.e. chromatographic peak shapes, co-elution of precursor/fragment ions, gas phase ion mobility profile, etc.

#### Performing database search

To perform the XL-peptide database search with MeroX do the following:

- Open MeroX v2.0.1.7.exe windows executable file by double clicking on it
- Load the search settings provided with this tutorial by clicking on **Settings**
- Click on **Load** and select the **timsToF_DSBU_25p_FDR.mxf** settings file and click **Open**

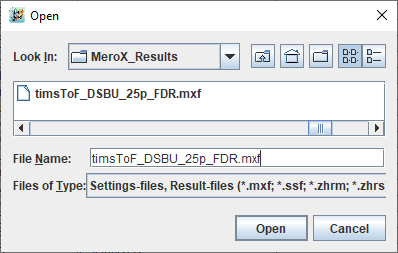

- Click **OK**
- Now let’s load the protein database by clicking on **Load Fasta**
- Select the **BSA_uniprot.fasta** and click **Open**

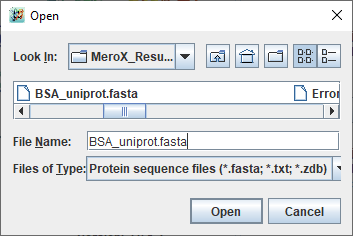

- Now let’s load one of the peak list files exported from DataAnalysis by clicking on **Load MS data** and select the following file:
  - **timsTOF_20230625_BSA-DSBU_S-Trap_pool_High_CE_01_RE_1_4682.mgf**
  - The name convention used here stands for *instrument*_*acquisition date*_*protein*-*cross-linker*_*digestion protocol*_*collision energy profile*_*technical replicate*_*auto sampler position*_*LC-MS/MS injection record*
  - The *.mgf* output from DataAnalysis that can be found within the .d folders has appended the DataAnalysis software version. We advise to make a copy of these files and delete the version number to avoid any issues with file name recognition in downstream analysis
- Click on **Start**
- MeroX will show a summary of the settings to be used in the search; double-check the values and click **OK**

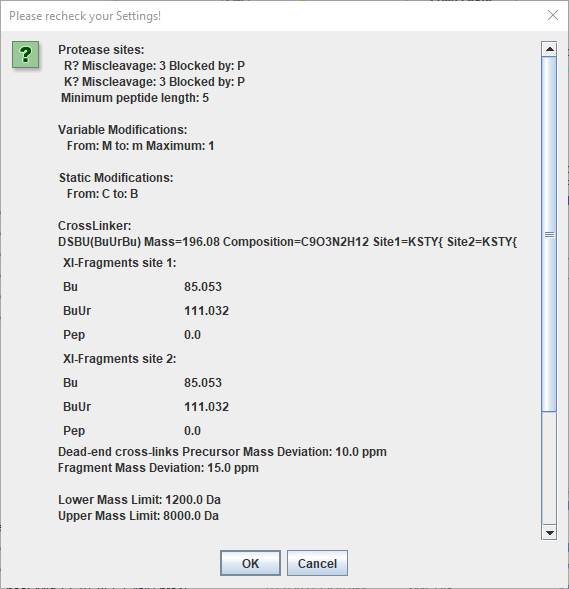

Once the search is finished two windows will open:

- The results table containing all the proposed cross-linked peptides and the corresponding PSM that supports the identity

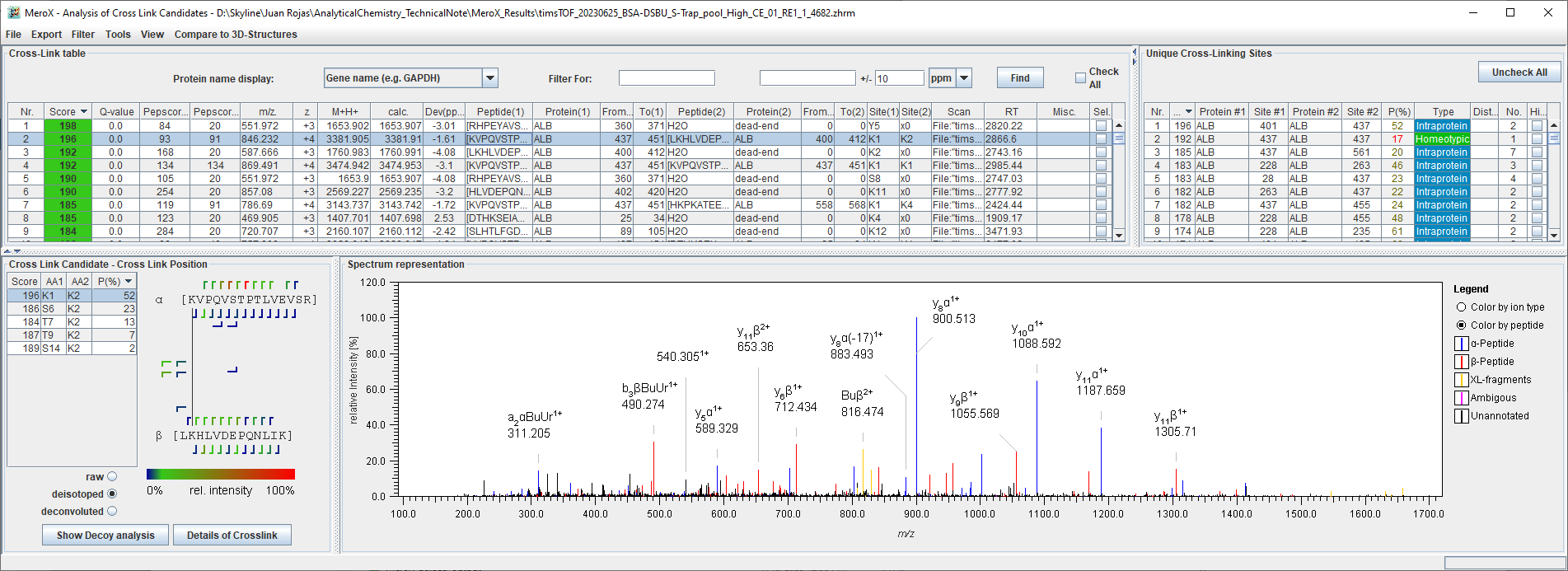

- And the Target-Decoy analysis corresponding to the searched file; we’ll have a look at this later in the tutorial

For now, just close both windows to avoid cluttering your desktop. Repeat the search steps for the remaining *.mgf* files until you have a total of 6 MeroX .*zhrm* results files.

#### Combining MeroX result files

Now let’s merge the MeroX results from all the replicates:

- Open MeroX v2.0.1.7.exe windows executable file by double clicking on it
- Click on **Combine result-files**
- Select all the 6 .zhrm files and click **Open**
- Assign a name for the combined files, e.g. **combined_BSA-DSBU_S-Trap**
- Click on **Save**
- Now open the new combined results by clicking on **Previous Data**
- Select the **combined_BSA-DSBU_S-Trap.zhrm** and click **Open**

A new file has been created containing the PSMs from all measurements. This action allows performing FDR estimation considering the whole dataset with respect to the MeroX score and the q-value is recalculated using all PSMs; that is the rank of each PSM with respect to all matches to the true and decoy database.

The document with the combined results contains PSMs with very low MeroX scores due to the lenient score and FDR settings used in the search. To have a FDR filtered document for visual inspection of cross-links let’s have a look first at the target-decoy analysis.

- On the results window click on **Show Decoy Analysis**

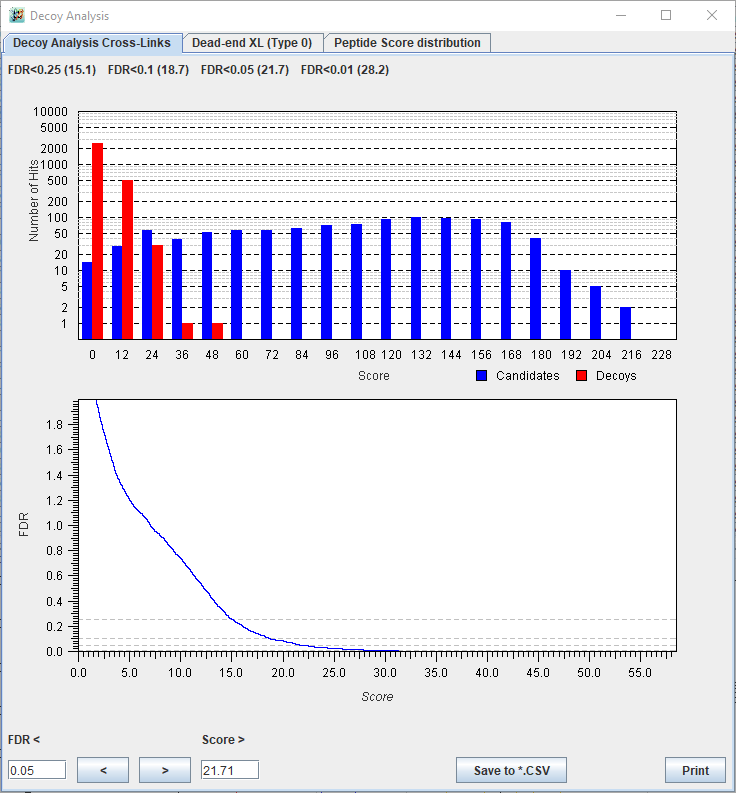

You’ll notice that the FDR is estimated for the different types of peptides that can be observed. Our focus here is FDR of cross-link peptides which can only be intra-protein cross-links in the system used for this tutorial; an additional tab would appear for inter-protein cross-links for multiprotein systems. On the top of each tab are summarized the MeroX scores that would be required to separate true hit from decoy hits at distinct FDR; e.g. applying a MeroX score cut-off of 28.7 would give us a dataset with 1% (0.01) false discovery rate. To apply this filter to the results file do the following:

- Take note of the MeroX score cutoff you want to apply; **28.7** for this tutorial
- Close the **Decoy Analysis** window
- Under the **Filter** menu click on **Advanced Filtering**
- Click on **Add new Filter**
- On the **Column** drop-down menu choose **Score** and set it as value has to be smaller than 28.7

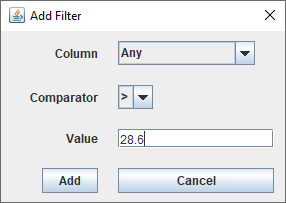

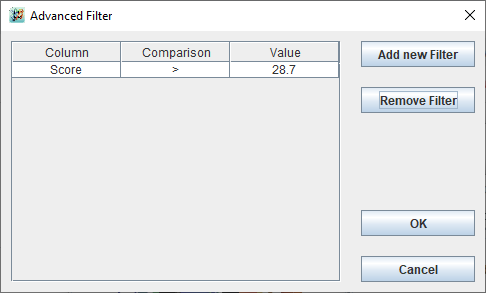

- Click **Ok** to remove all PSMs with a score lower than 28.7

BSA is a well characterized system that produces enough unique intra-protein cross-links that can be visualized with short tryptic peptides favorable for ionization. This diversity seems to be enough for useful FDR filtration, but this might not be the case for other protein systems where only few detectable cross-linked peptides are present. In the latter scenario the true vs. decoy PSM distribution cannot be distinguished reliably, making the FDR filtering strategy not viable. Instead, it might be better to rely on the MeroX spectral cutoff score as a guide on which matches to trust. A score of 70 ≤ has been recognized as a good ballpark score for reliable identification and recently^7^ a similar score was found to accurately estimate the FDR when analyzing a synthetic library of cross-linked peptides. PSM with scores of 70 can still be true, but we advise users to be very critical on the manual inspection of these matches and to consider adding these as priority targets for sample re-analysis (e.g. using DDA-preference lists or PRM) to improve spectral quality.

#### Extracting ion mobility library for Skyline

DataAnalysis performs peak picking in the retention time and ion mobility dimensions for all precursors exported to the *.mgf* files. This information is written in the **TITLE** line for each precursor ion and below is an example (visualized with Notepad++ v8.4.7) of the format created by the DataAnalysis processing script used for this dataset:

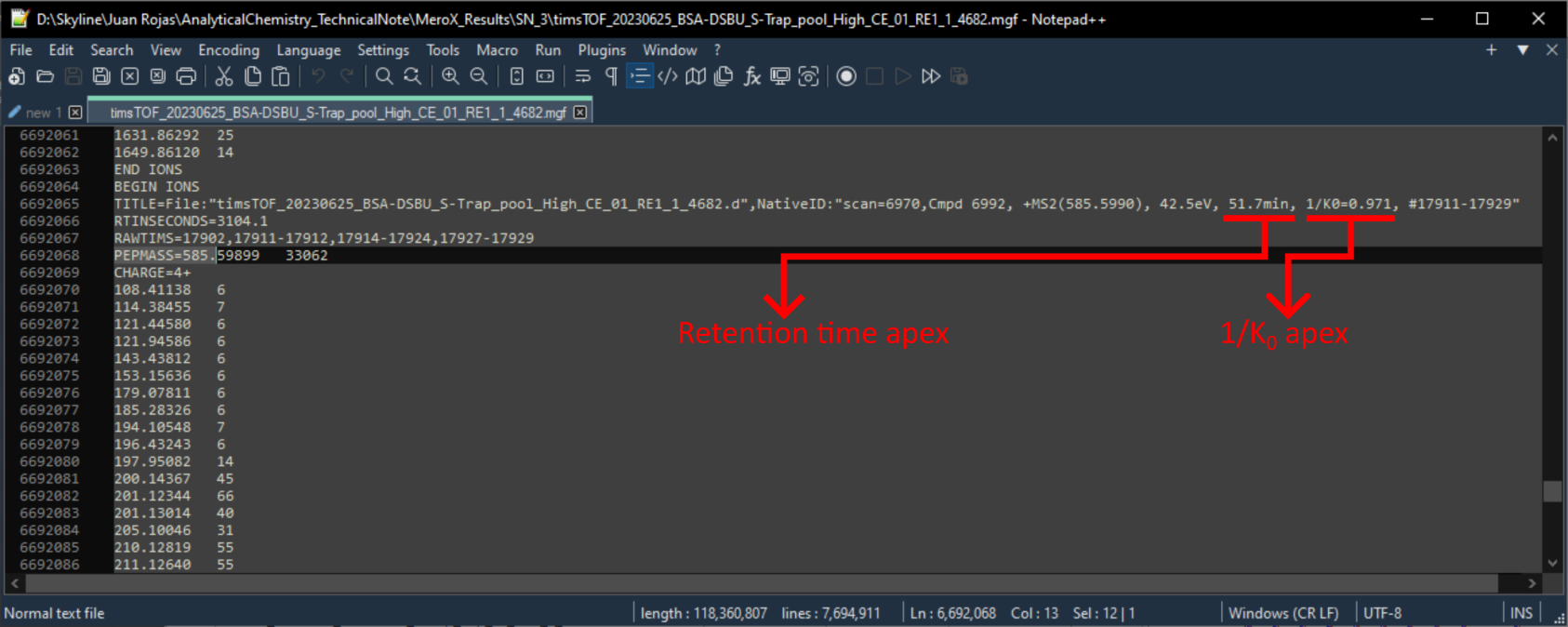

This information is passed to the **Scan** column of the MeroX results:

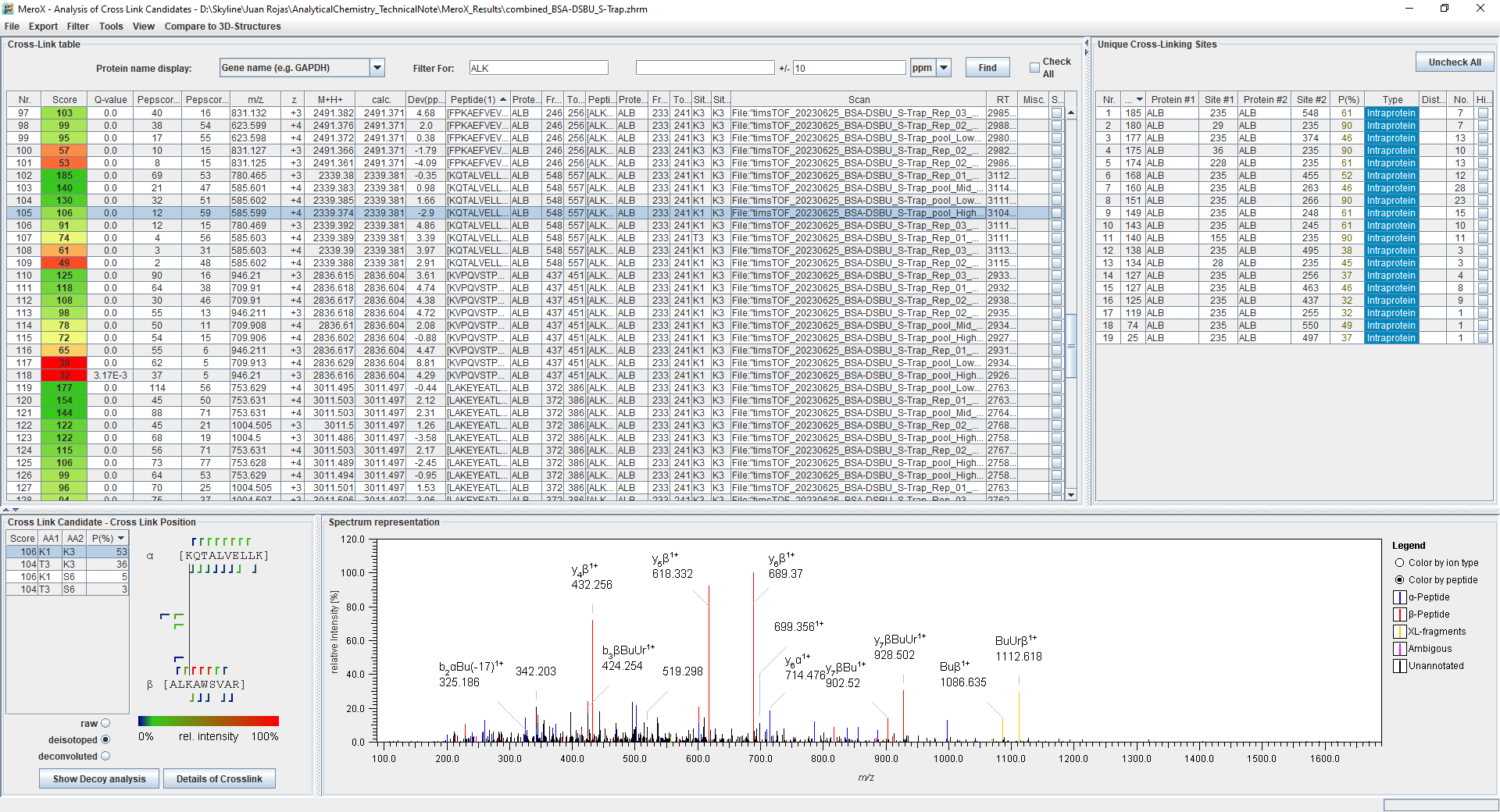

These results can be exported to a comma separated values (*.csv*) file by clicking on **… to CSV** inside of the **Export** menu. Users are encouraged to mine the **1/K_0_** associated with each precursor as they find convenient, but for this tutorial you will have a text file formatted in the syntax that Skyline expects from an ion mobility library; file: *MeroX_IMS_library.txt*.

NOTE: This was generated with a custom R code. The R code for the IMS library extraction and all further data processing with R is documented with an accompanying R Notebook that you can find in the “Supplementary Files: R Script and Data Integration“ folder located in <https://panoramaweb.org/XL-MS_MeroX_Skyline.url>.

The IMS library was created by extracting the reported 1/K_0_ from all PSMs corresponding to a specific precursor and then determining the average 1/K_0_ of these PSMs. Additionally, the range of all the values extracted was calculated to detect the possibility of multiple gas phase conformers of a specific precursor ion. These multiple gas phase conformers can be pointed out by plotting the calculated average ion mobility vs. the range of the values averaged:

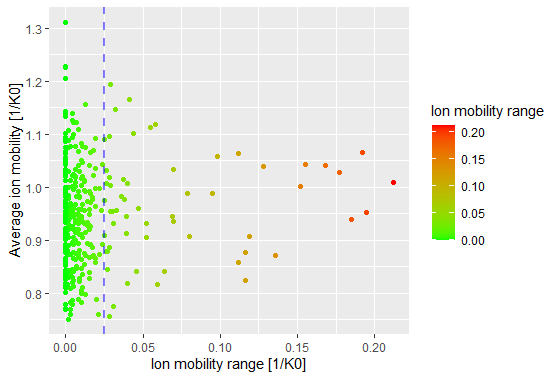

Most estimated average ion mobility values have very small averaging ranges corresponding to small deviations during peak picking. These ranges are well below 0.025 1/K_0_ (blue dashed line) which correspond to the precursor mobility window allowed for grouping fragment ion spectra (see **Method parameters** section). Values that are beyond this range could happen due to precursor splitting or because two different gas phase conformers were sampled. The former occurs if the mobility range allowed for grouping spectra was smaller than the precursor ions distribution in the mobility dimension. The ion mobility averages of precursors with large sampling range require correction. Assuming there were enough points for all gas conformers then these could be clustered and extract the average values from each cluster of conformers, but unfortunately some of the conformers were only sampled once in this dataset. Therefore, we’ll show how to correct these values manually later in the tutorial.

### Standardization of search results with proXL

To get MeroX results into a format that is readable by Skyline we need to rely on a software converter that will translate the MeroX cross-linked peptide results into a standardized XML schema known as protein cross-linking (proXL) XML^3^. To ensure full compatibility with Skyline it is important that the **Scan** **column** in the **MeroX results** contains the **TITLE** information (displayed previously) as the **scan=INDEX** will be used to map the PSMs when Skyline creates the spectral library; example:

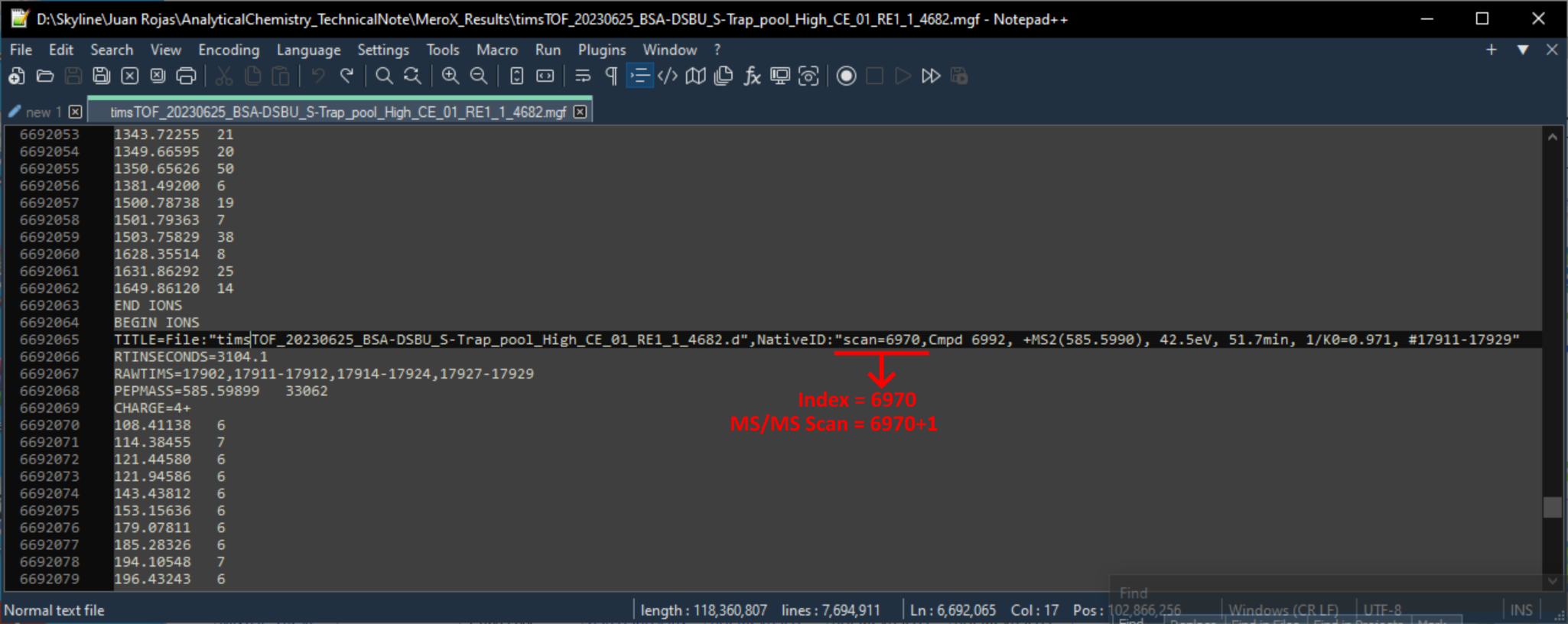

NOTE: The first MS/MS scan in the .mgf files has an index=0 which is why the example above corresponds to the title of the 6971 scan in the file.

To perform the results conversions do the following:

- Download *merox2ProxlXML.jar* java executable file from <https://github.com/yeastrc/proxl-import-merox> and place the java software in the same folder that contains the MeroX results
  - Requires Java to be installed in your PC; the workflow presented here was tested with Java Version 8 Update 351
  - For this tutorial material, the java software has already been downloaded and you can find the program in the “MeroX Results” folder
- Type **cmd** in folder directory and press enter

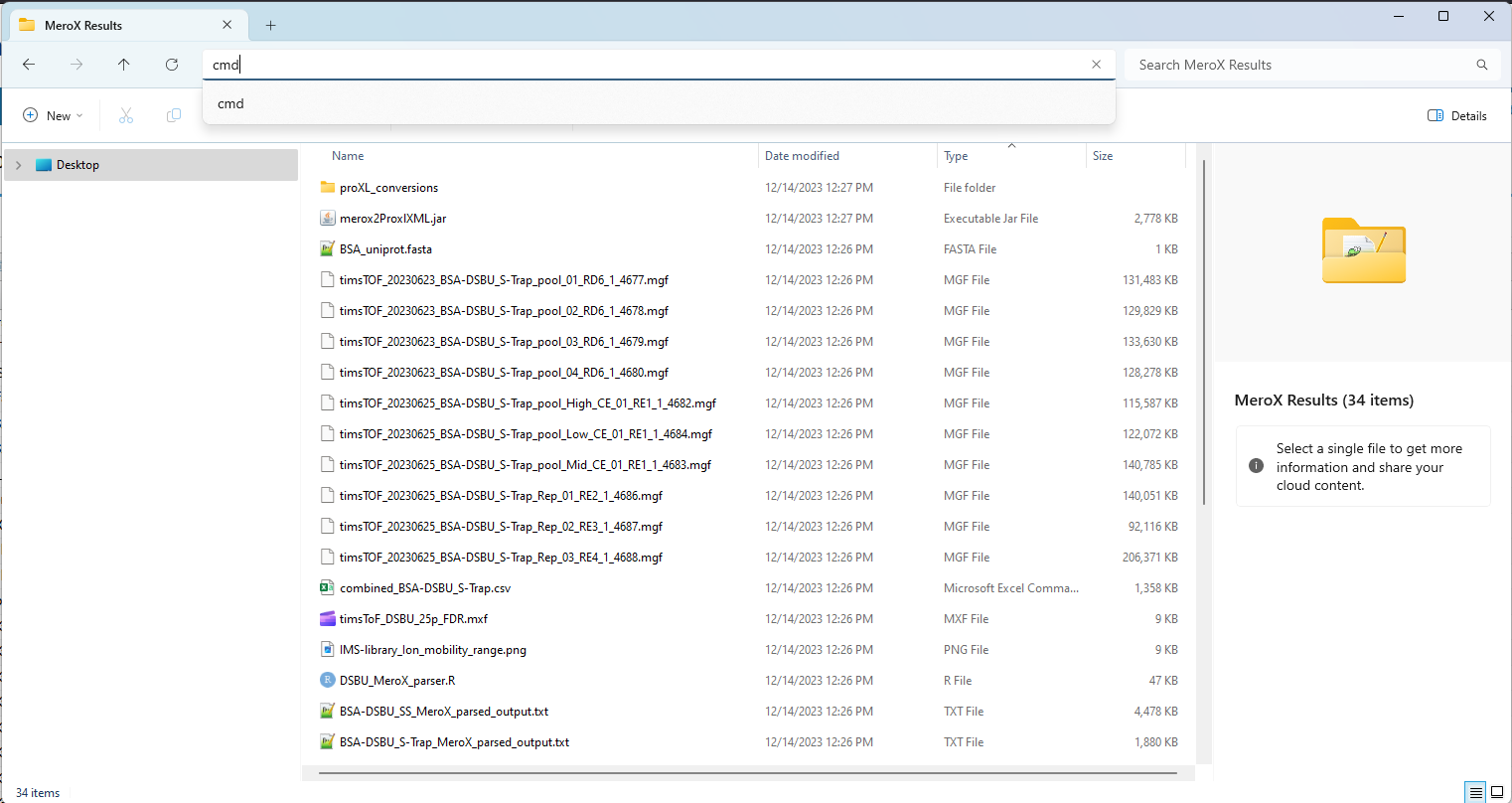

- After you have pressed **enter**, the command line window should have opened with the working directory already defined as the folder where you type **cmd**.

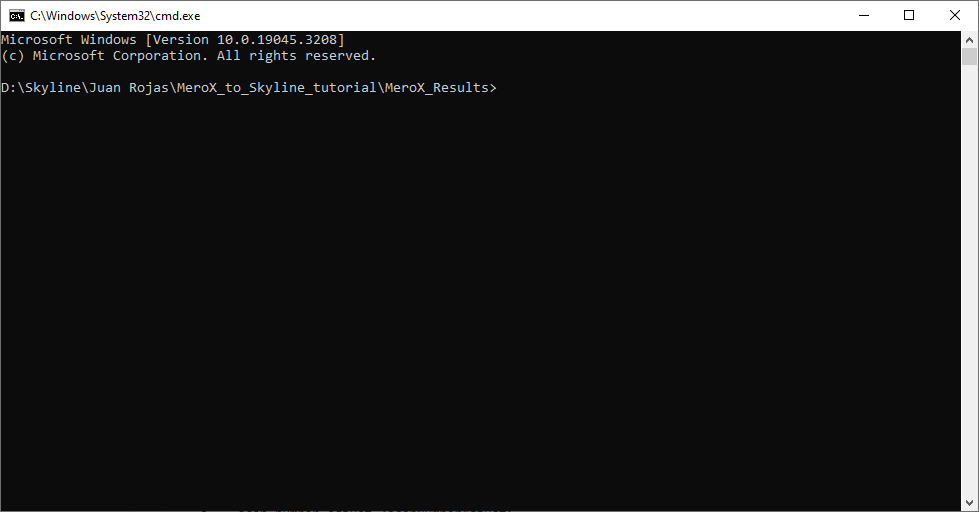

- Now open the **MeroX_to_proXL_conversion_commands.txt** in a text editor (e.g. Notepad or Notepad++) and copy the second line of text
- Back in the command line window, first **left-click** after “MeroX_Results>” and then **right-click** to paste the line of text copied before

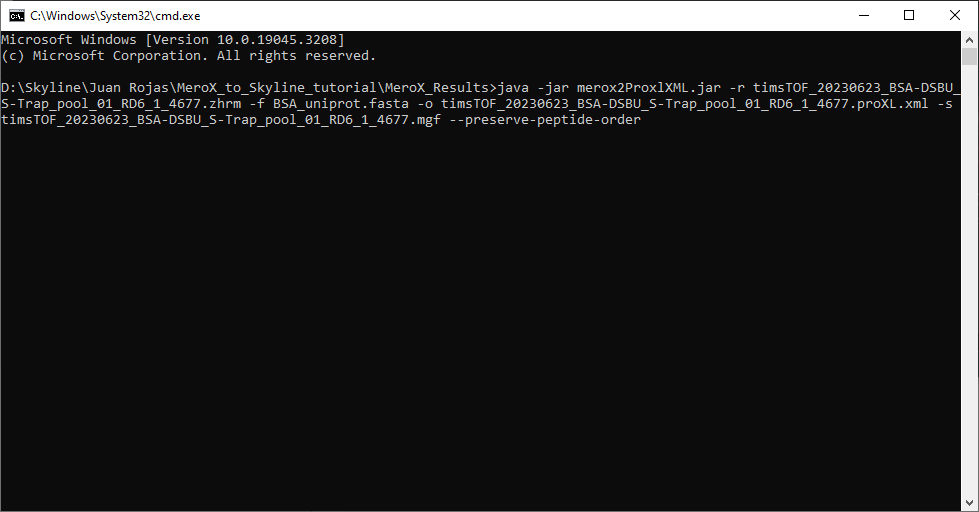

- The syntax of the commands used can be found in: <https://github.com/yeastrc/proxl-import-merox>
- Press **Enter** in your keyboard

Now you should have the first *.proXL.xml* file in the same directory as your native MeroX results:

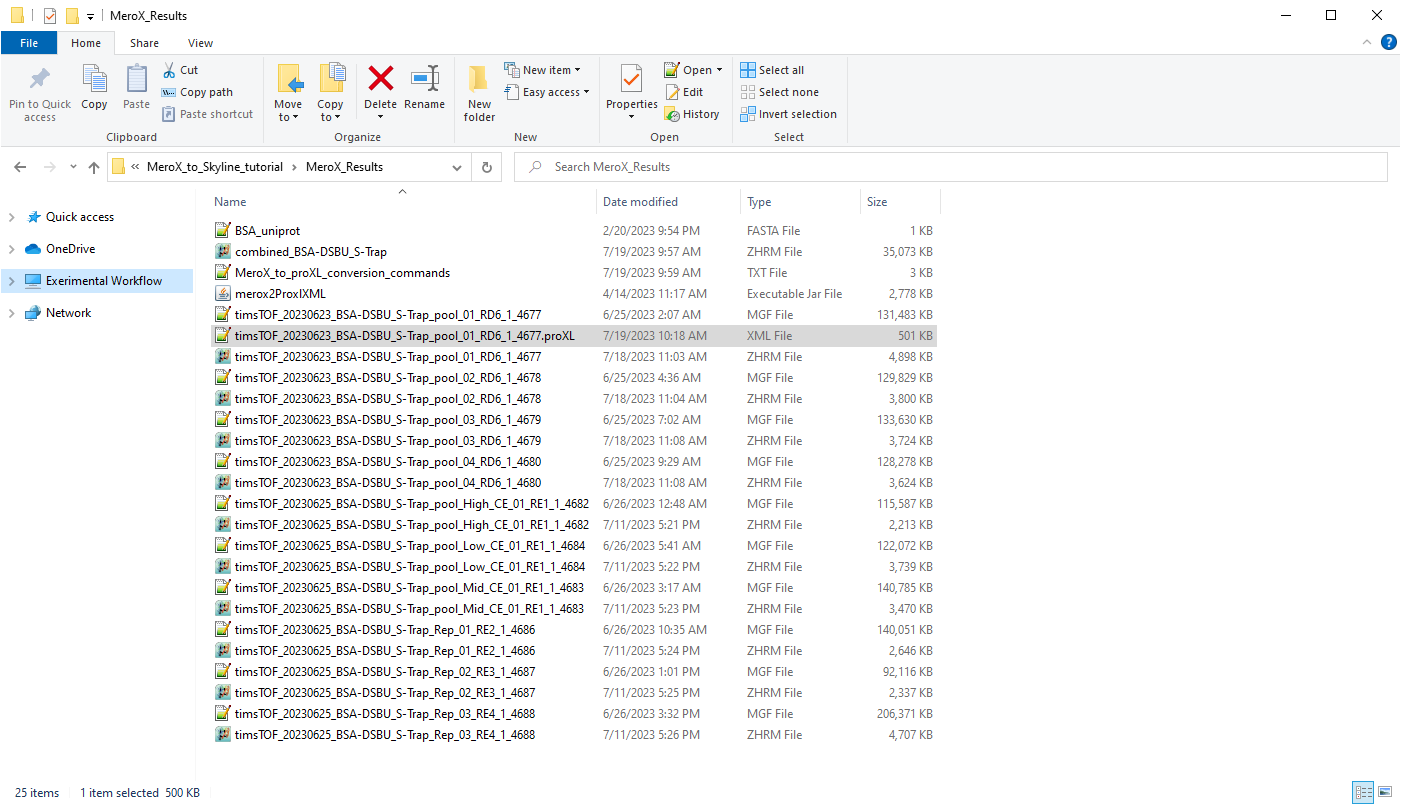

Perform the previous steps until you have one *.proXL.xml* file for each MeroX results file. In the end, you will have three files with the same name, but different file extensions corresponding to:

1) The MS/MS data stored in the *.mgf* file
2) The native MeroX results stored in the *.zhrm* file
3) The results converted to the ProXL XML database schema in the *.proXL.xml* file

#### Combined results library

Since version 1.2.0 the merox2ProXLML converter tool is capable of taking as input a combined MeroX results file. This improvement allows working with the global FDR estimation that MeroX performs when it combines results from multiple LC-(IMS)-MS/MS measurements.

Note that for this to work, the origin file name from annotated MS/MS scans should be in the TITLE header of each MS/MS spectrum stored in the .mgf files. The converter is able to extract the information associated to each matched spectrum. Therefore providing the scan file name is not necessary when using the tool from the command line, e.g.:

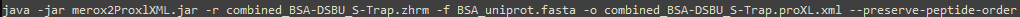

### Analyzing results in Skyline

#### Setting up the document

Before loading the raw data we need to setup the Skyline document with the correct settings and targets.

- **Open** Skyline-daily
- Select the **Proteomics interface**
- Click on **Blank Document**
- Remove any settings left from previous projects by clicking on **Default** under the **Settings** menu

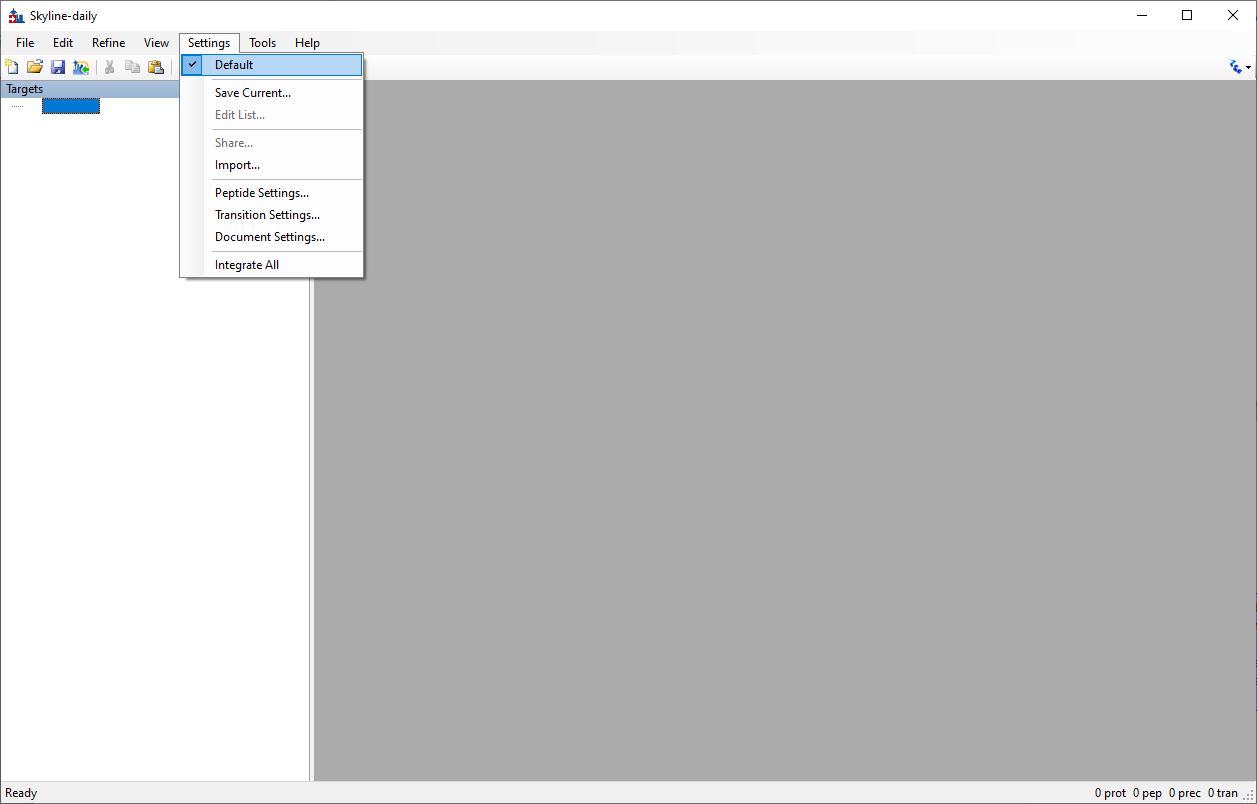

- First save the skyline by clicking on **Save** under the **File** menu (or **Ctrl+S**) and save the document in the same folder containing the raw MS files provided with this tutorial; name suggestion: *MeroX_BSA-DSBU_S-Trap.sky*

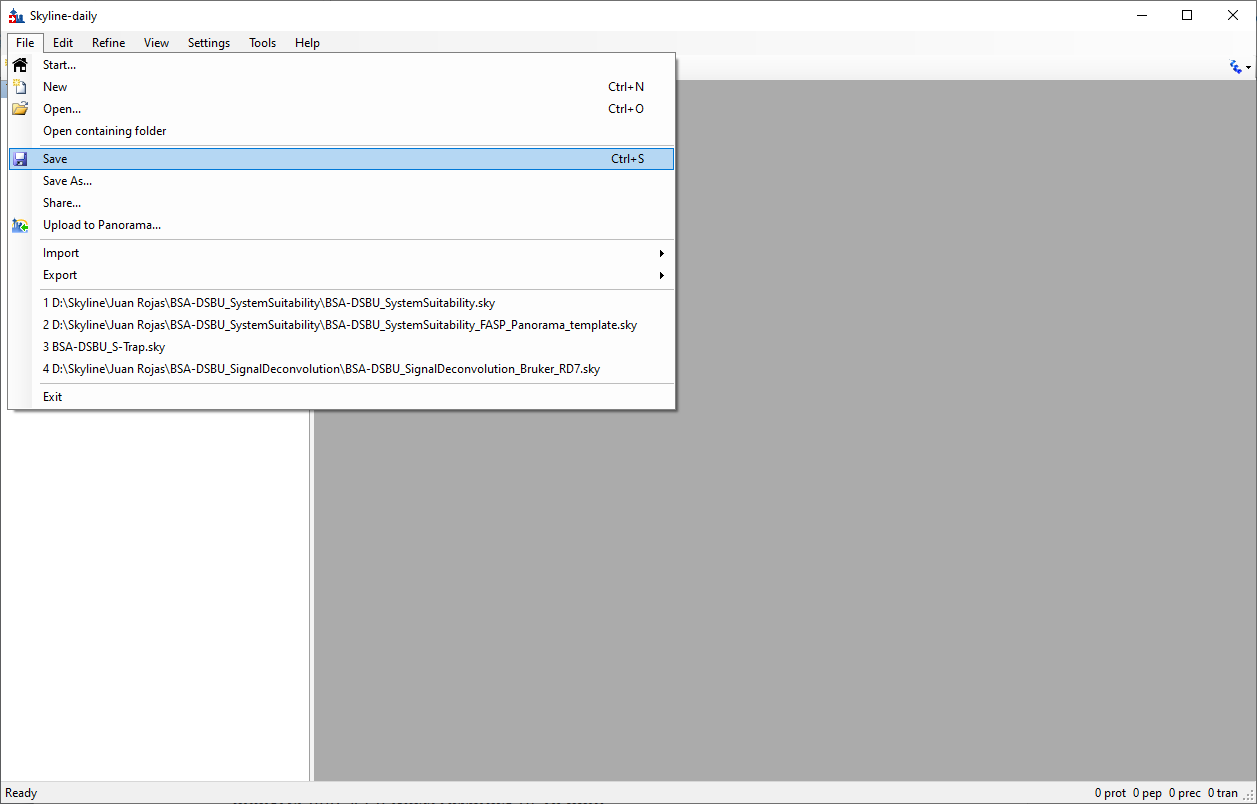

##### Transition settings

First let’s set up the settings for DDA data with MS1 filtering workflow:

###### MS data extraction settings

- Click on **Transition Settings…** under the **Settings** menu
- Select the **Full-Scan** tab
- On the **MS1 filtering** settings set the selection of **Isotope peaks included** to be decided by **count** and set to 3 peaks
- For the extraction window in the *m/z* dimension select **Centroided** under **Precursor mass analyzer** and set a **Mass Accuracy** of **10 ppm**
- For **Retention time filtering** allow Skyline to generate EICs by **Use only scans within 5 minutes of MS/MS IDs**

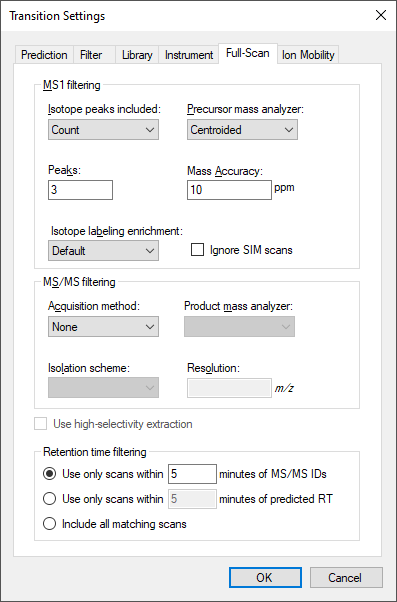

###### Target ions definition

- Now select the **Filter** tab
- We will consider **Precursor charges** ions ranging from **2-to-6** charged states and since we will only look add DDA data here we only want Skyline to calculate precursor **Ion type** (i.e **p**)

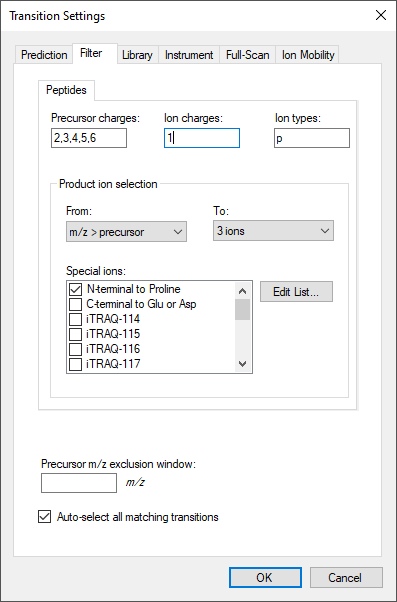

###### Targets error tolerance and selection criteria

- Now select the **Library** tab
- Here only change the **Ion match tolerance** to **0.05 *m/z*** to restrict the matches of the theoretical *m/z* values calculated by Skyline to those observed in the spectral library

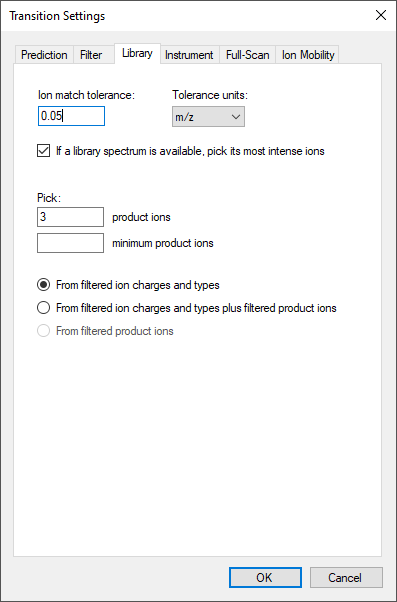

###### Ion mobility library

- Finally, select the **Ion Mobility** tab
- Under the **Ion mobility library** drop-down menu click on **<Add…>**

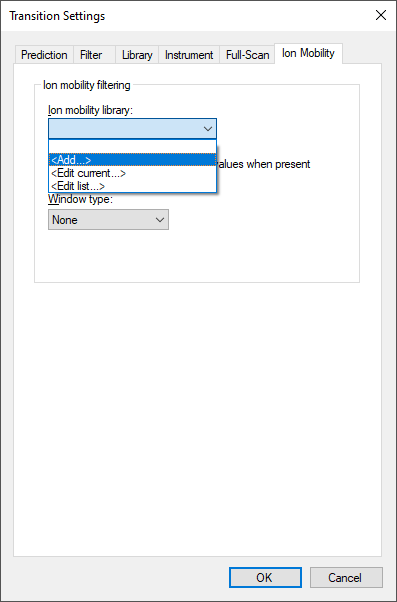

- Once you are in the **Edit Ion Mobility Library** click on **Create…** to create an ion mobility library (i.e., .imsdb) file
- Save the library as *MeroX_BSA-DSBU_166ms.imsdb* in the same folder where you saved the Skyline document
- Now to import the IMS library provided with this tutorial open **Excel** and **import** the file stored inside the **MeroX_Results** folder
- **Without** selecting the **column titles**, **select** all the cells from the **first 5 columns** and copy the values (**Ctrl+C**); that is 279 rows and 5 columns
  - We will ignore the ion mobility range for now

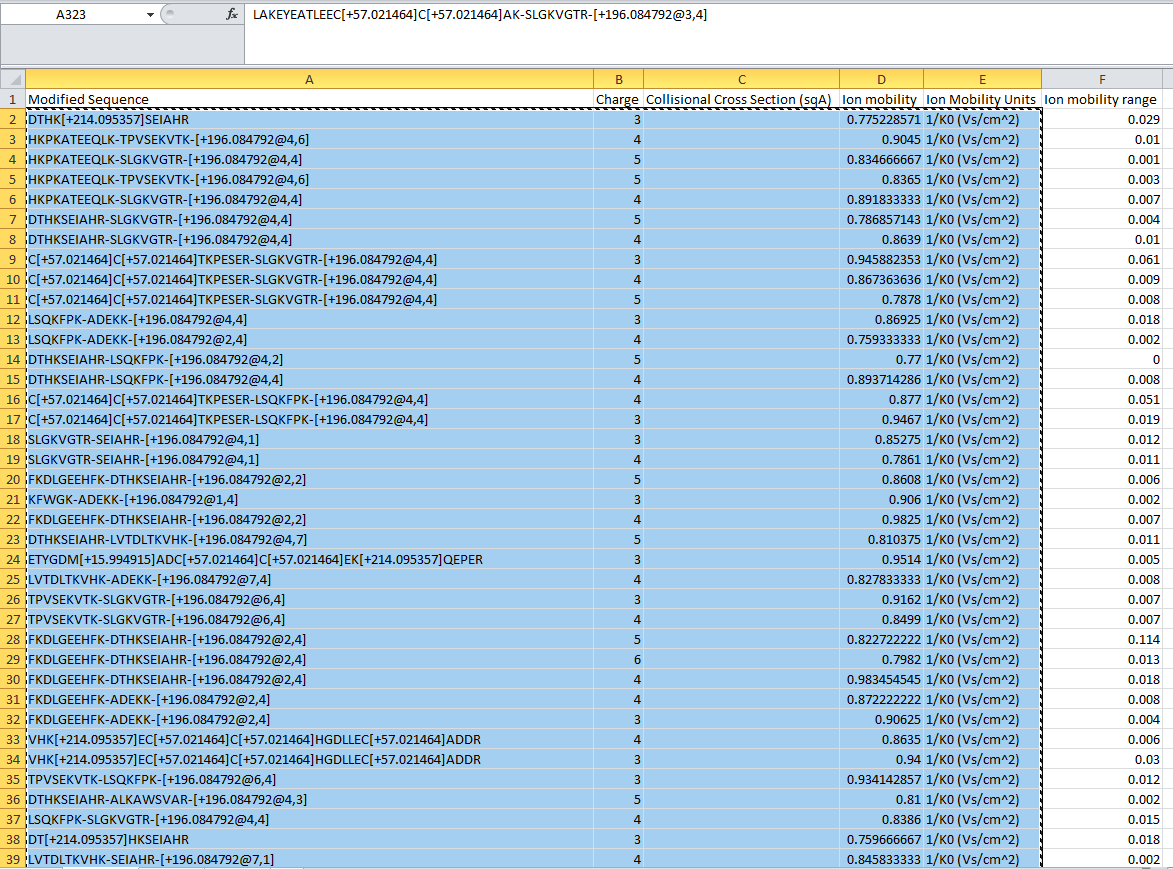

- Back on the **Edit Ion Mobility Library** under the **Measured peptides** table, left-click on the upper left corner box (highlighted in blue below) and paste the copied cells (**Ctrl+V**)

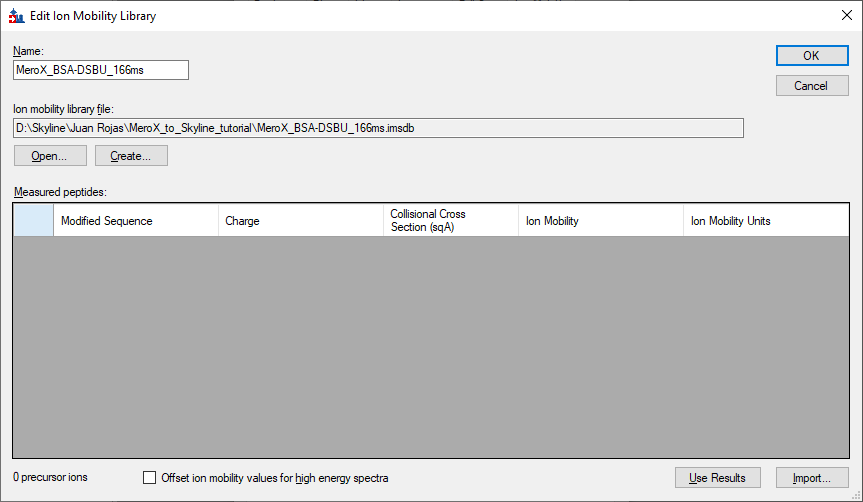

- Once pasted correctly, the populated table should look like this:

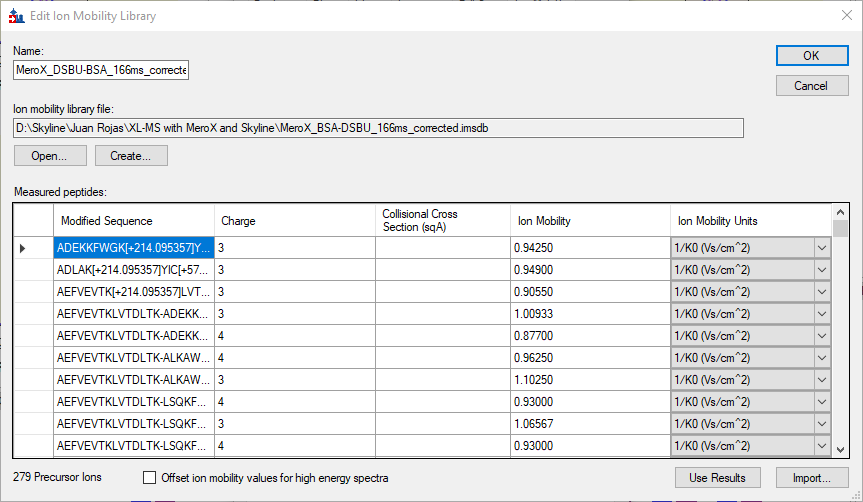

NOTE: You can also rely on Skyline to determine the center of the Ion mobility distribution once the EICs have been generated. This is done by using the **Use Results** button in the **Edit Ion Mobility Library**. However, if there are severe integration interferences or the wrong chromatographic peak was chosen by Skyline, then a wrong ion mobility value could be determined. You will get the best results of the **Use Results** function if refined Skyline documents are used. Here we have defined the Ion mobility library prior to EICs generation to facilitate automatic peak picking by Skyline.

- Click **Ok** and this will save the ion mobility library
- Once you are back in the **Ion mobility** tab select the **Window type** as **Resolving power** and set it to 30
  - The value you set in the **Resolving power** box is defined as follows in Skyline: “Determines window size for ion mobility filtering in chromatogram extraction (for example, drift times): For resolving power “R” at drift time “T”, the width “W” of the drift time filter window centered at T is W = 2T/R”

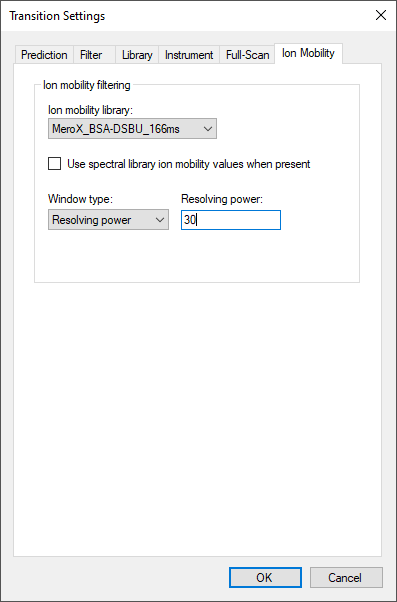

- Click **OK** and save the document (**Ctrl+S**)

##### Peptide settings

Now let’s define the settings required for Skyline to be able to compute the correct targets for EICs.

###### Modifications definition

- Click on **Peptide Settings…** under the **Settings** menu
- We have to define the cross-linker modification Select the **Modifications** tab
- Next to the list of **Structural modifications** click on **Edit list…**

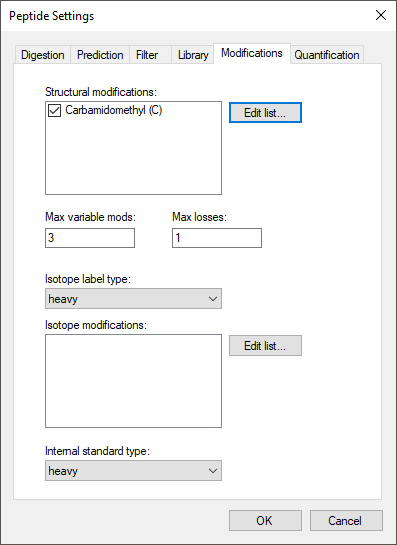

- Click **Add…** and define the **DSBU (XL)** modification as follows:
  - **Amino acid** specificity: K, S, T, Y
  - **Tick** the modification as a **Crosslinker**
  - **Chemical formula** set to only the final mass increase of a doubly reacted DSBU molecule: **C9O3N2H12**
  - Additionally, to compute the cross-linker MS-cleavable fragments set the following neutral **Losses** by clicking on **Loss >>** and then clicking on the
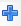
 to add the following mass losses:
    - **C4H7NO** with a **charge** loss of **0**
    - **C5H5NO2** with a **charge** loss of **0**
    - **C9O3N2H12** with a **charge** loss of **0**
- The mass losses from the defined cross-linker correspond to the losses that produce the fragments defined in the MeroX search settings, including the full loss/connection of the cross-linker from one of the cross-linked peptides

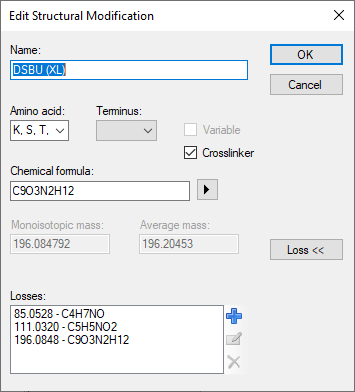

- Click **OK**
- Additionally, we will define the modification of a dead-end products when one of the sites has reacted with a peptide and the other with a water molecule so click **Add…** again
- Define the **DSBU (dead-end water)** modification as follows:
  - **Amino acid** specificity: K, S, T, Y
  - **Tick** the modification as a **Variable**
  - **Chemical formula** set to only the final mass increase of a doubly reacted DSBU molecule: **C9O4N2H14**

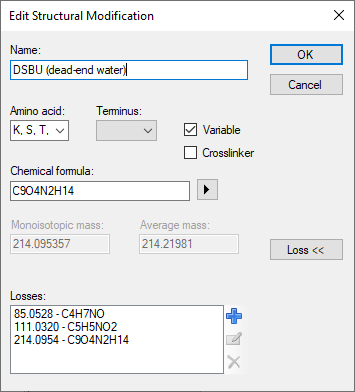

- Additionally, to compute the cross-linker MS-cleavable fragments set the following neutral **Losses** by clicking on **Loss >>** and then clicking on the
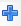
 to add the following mass losses:
  - **C4H7NO** with a **charge** loss of **0**
  - **C5H5NO2** with a **charge** loss of **0**
  - **C9O4N2H14** with a **charge** loss of **0**
- Click **OK** to return to the **Edit Structural Modifications** tab
- Now let’s add a final modification that is defined in the PTM library contained within Skyline. Click **Add…**
- Once in the **Edit Structural Modification** window click on the drop-down menu

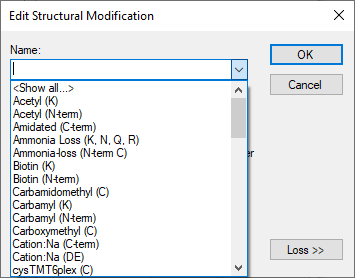

- Type **Oxidation (M)** and click on the option shown in the PTM list to autofill the definition of this PTM
- Click **OK** twice to return to the **Peptide Settings Modifications** tab and make sure that the defined and added modifications are ticked to make sure that Skyline considers them when Skyline interprets the peptides we would like to add

###### Maximal modification and losses

- Set **Max losses** to **2** (this will allow Skyline to consider secondary fragmentation events that produce the cross-linker MS-cleavable fragments) and that the following modifications are ticked

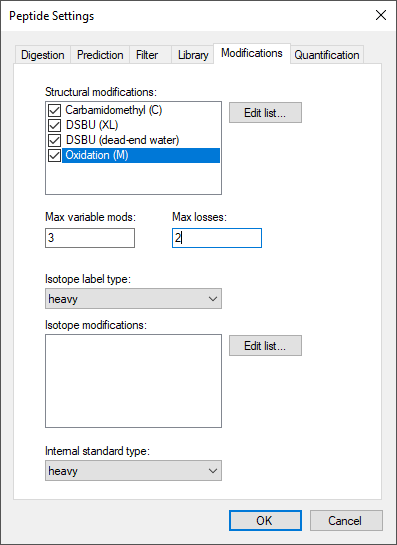

###### Spectral library creation

- Now select the **Library** tab
- Next to the **Libraries** list click on **Build…**
- **Name** the library as “MeroX_BSA-DSBU_S-Trap_q_0.05” and select the **Output Path** to be the same directory where your Skyline document is saved
- Select the **Data Source** as **Files** and make sure to tick **Keep redundant library**

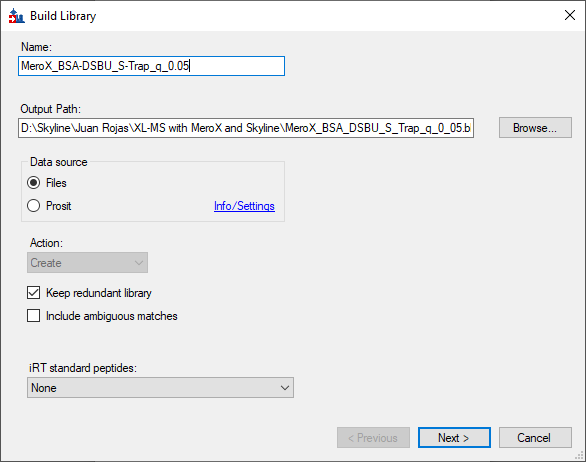

- Click **Next >**
- On the right side of the **Input Files** list click on **Add Files…** and browse inside of the “MeroX Results” folder and select the **10** *.proXL.xml* files that should be in this folder
- Click **Open**
- Once the results files are loaded into the **Input Files** list, left-click on the first cell of the **Score Threshold** score and make sure the value is set to **0.05**
  - This score corresponds to the q-value, defined as the minimal FDR threshold at which a PSM would be considered. Any PSMs with a q-value higher than 0.05 are excluded from the spectral library generation
  - The **Score Threshold** here is applied per results file and not with respect to the FDR estimation of the whole dataset as we did with the combined results previously

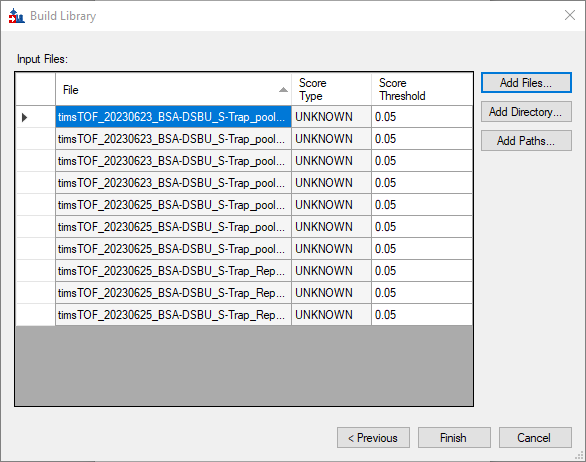

- Click **Finish** and the library creation should start

NOTE: it is important that the files containing the MS/MS spectra associated to each PSM (that is the *.mgf* files for this tutorial) are stored in the same folder as the *.proXL.xml* files

- When the library creation is finished a message saying “Library MeroX_BSA-… build completed.” should show up on the left-bottom corner of the main Skyline window
- Click **OK** and save the document (**Ctrl+S**)

###### Combined results spectral library

Note that in the example above, the q-value of 0.05 was selected as the filtering threshold for each file. However, this would apply an FDR estimation from a single LC-MS/MS run instead of using a global estimation. In experiments where there are very few true PSMs per file, a correct modelling of the FDR might not be possible. Therefore, we will showcase an alternative using the **combined_BSA-DSBU_S-Trap.zhrm** file that was created earlier.

As long as the .mgf formatting is kept as described here (so the file name is transferred to the MeroX scan column), now the ProXL converter tool is capable of using the combined MeroX results to create a single .proXL.xml file that contains the spectra matches from the whole dataset and uses the global FDR estimation performed by MeroX.

- Now select the **Library** tab again
- Next to the **Libraries** list click on **Build…**
- **Name** the library as “MeroX_combined_BSA-DSBU_S-Trap_q_0.05” and select the **Output Path** to be the same directory where your Skyline document is saved
- Select the **Data Source** as **Files** and make sure to tick **Keep redundant library**

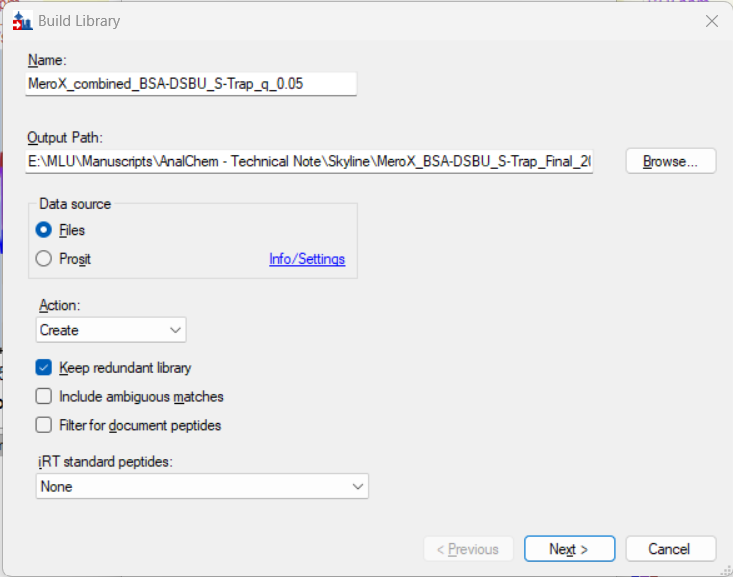

- Click **Next >**
- On the right side of the **Input Files** list click on **Add Files…** and browse inside of the “MeroX Results” folder and select the *combined_BSA-DSBU_S-Trap.proXL.xml* file
- Click **Open**
- Once the results files are loaded into the **Input Files** list, left-click on the first cell of the **Score Threshold** score and make sure the value is set to **0.05**
  - This score corresponds to the q-value, defined as the minimal FDR threshold at which a PSM would be considered. Any PSMs with a q-value higher than 0.05 are excluded from the spectral library generation

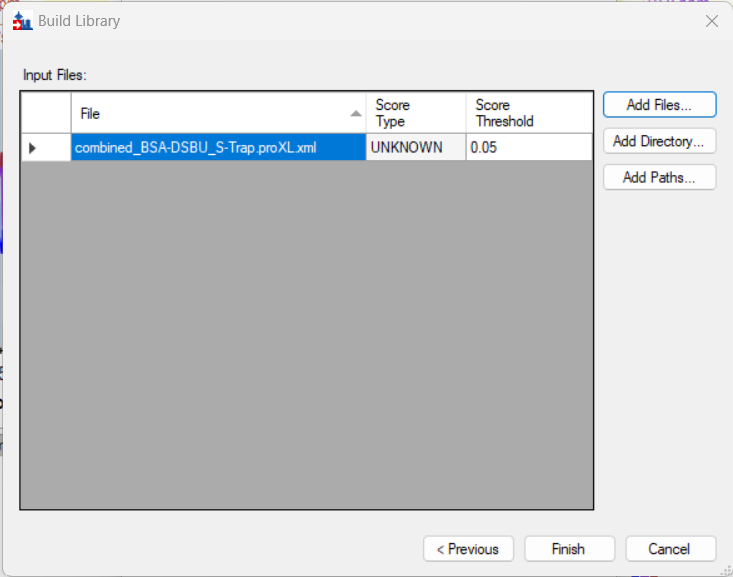

- Click **Finish** and the library creation should start

NOTE: it is important that the files containing the MS/MS spectra associated to each PSM (that is the *.mgf* files for this tutorial) are stored in the same folder as the *.proXL.xml* files

- When the library creation is finished a message saying “Library MeroX_BSA-… build completed.” should show up on the left-bottom corner of the main Skyline window
- Click **OK** and save the document (**Ctrl+S**)

#### Importing peptide ion targets

Now it is time to add the targets to the document. For this we have to first have a look at the spectral library we just created and add the targets from there.

- Under the **View** menu click on **Spectral Libraries**
- On the **Library** drop-down list make sure that the **MeroX_BSA-DSBU_S-Trap_q_0.05**
- You will notice that the **Add Modifications** window will pop-out showing some mass shifts allocated to specific amino acids that cannot be interpreted by Skyline.
  - These are PTMs that MeroX mapped to non-cross-linked peptides and do not correspond to the dead-end product with water that we defined earlier. For the purpose of this tutorial, these unknown PTMs are not relevant so we will ignore them
- Click **OK** to close the **Add Modifications** window to have access to the **Spectral Library Explorer**

Here you can visualize the annotated PSMs that passed the filter set during the library creation. Cross-linked peptides are visualized by the amino acid sequences connected by a dashed line with the cross-linking positions highlighted in **bold green**. Peptides that cannot be interpreted by Skyline, such as those containing the unknown mass shifts, will have the missing minimalistic spectrum symbol as shown below:

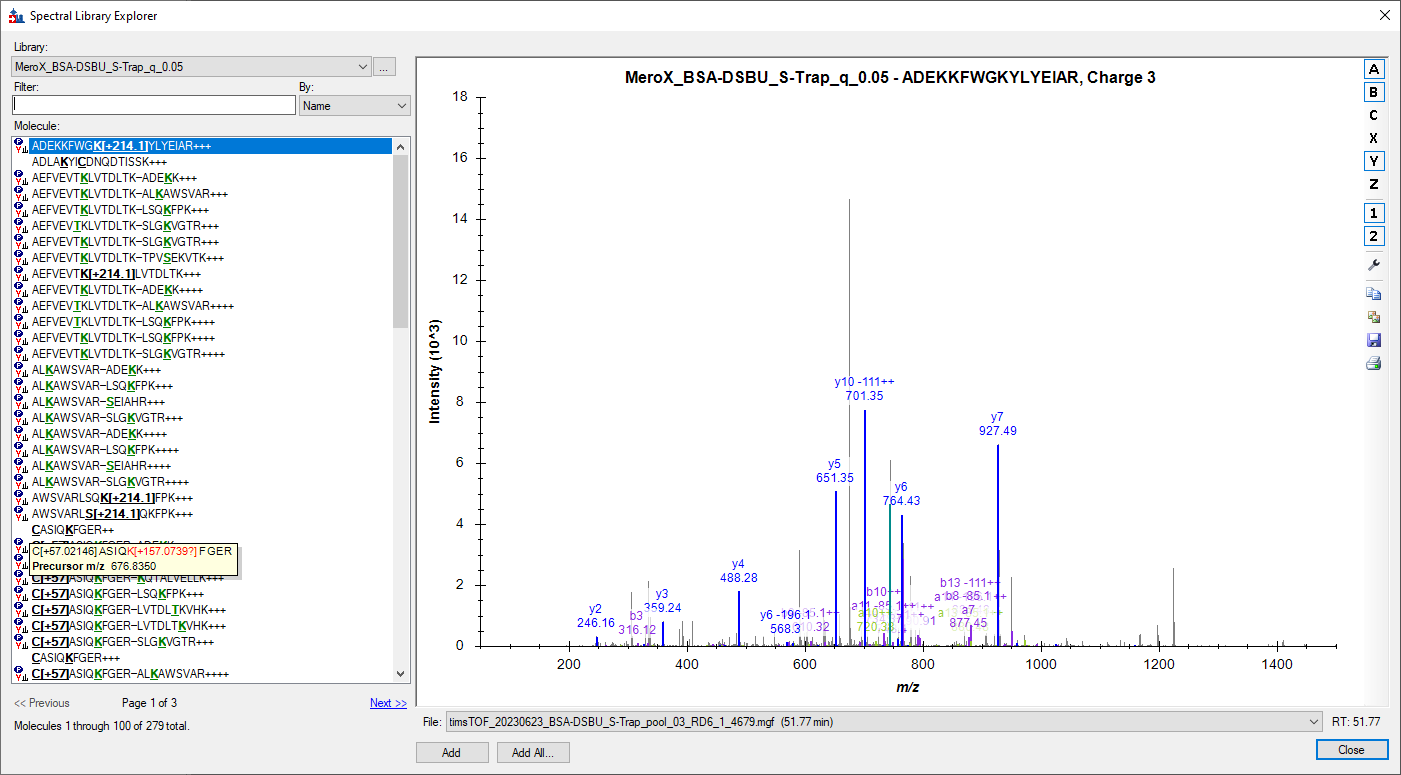

The explorer allows you to search the peptides in the library based on **Name, Precursor *m/z*, Charge, or Adduct** by clicking on the **By** drop-down list. Additionally, if the peptide in the library is supported by various PSMs and the **redundant library** was kept, these PSMs can be visualized one-by-one by clicking on the **File** drop-down list. Let’s look at an example:

- Make sure the **Name** filter is selected and type “ALKAWSVAR”
- Select the “ALKAWSVAR-ADEKK+++” peptide
- On the **File** drop-down list select timsTOF_20230625_BSA-DSBU_S-Trap_pool_Low_CE_01_RE1_1_4684.mgf (39.39)
- **Right-click** on the spectrum and make sure the following are **selected/highlighted**:
  - Ion Types:
    - N-Term: B
    - C-Term: Y
    - Losses: -85, -111, -196, -214, -64
  - Precursor
  - Ranks
  - Observed *m/z* values
  - Auto-scale Y-axis
  - Make sure that charges 1 and 2 are selected
- Zoom the spectrum in the *m/z* range between 200 and 1150

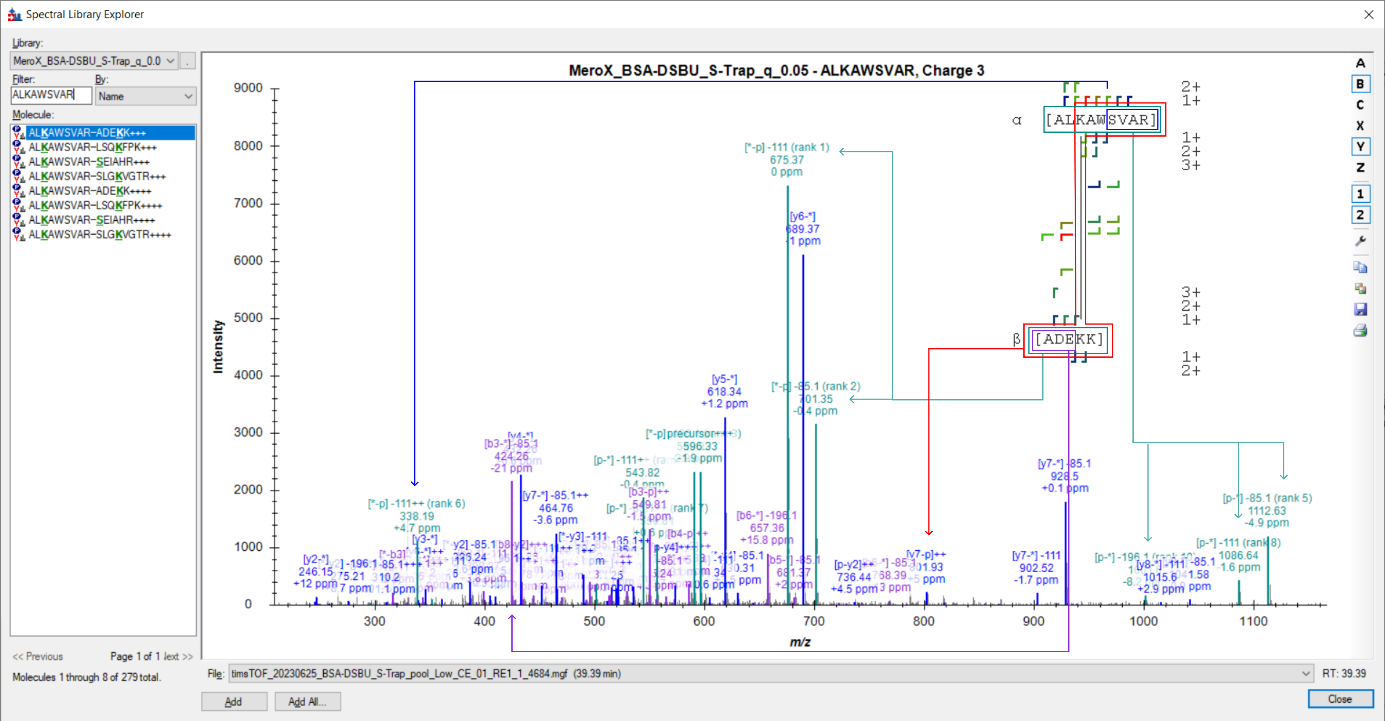

The way that fragment ions are annotated for cross-linked peptides in Skyline will be illustrated with the following examples:

- [y4-*]: corresponds to the singly charged y_4_-ion of the left peptide; the cross-linker does not play a part for this ion
- [*-b3]: corresponds to the singly charged b_3_-ion of the right peptide; the cross-linker does not play a part for this ion
- [p-* -85.1]-85.1: corresponds to one of the MS-cleavable fragments of DSBU where the full left peptide is observed with a loss of a part of DSBU (i.e. 85.0528 u)
- [*-p -111]-111: corresponds to one of the MS-cleavable fragments of DSBU where the full right peptide is observed with a loss of a part of DSBU (i.e. 111.0320 u); corresponds to the opposite
- [y7-p]++: corresponds to the doubly charged ion that contains the left y7-ion and the full right-side peptide still attached by the full DSBU cross-linker

NOTE: the MeroX cleavage annotation on the linked peptide sequences is not available in Skyline. Here the image was edited for illustration purposes. We advised to keep using MeroX for manual inspection of questionable PSMs as this tool allows linking directly the annotated ions to the matching fragment cleavage sites.

Now let’s move to importing the peptides to the Skyline document:

- Clear any text or numbers from the **Filter** box
- Click **Add All…**

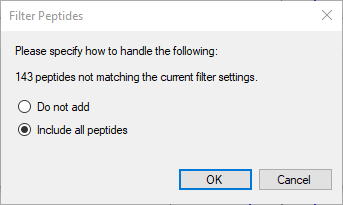

- If you get a window indicating that **X peptides not matching the current filter settings** then select **Include all peptides** and click **OK**
- After you get a window indicating the amount of targets that will get added to the document click **Add All**

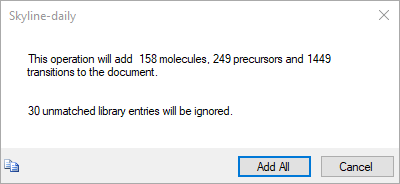

- Close the **Spectral Library Explorer**
- Under the **View** menu click on **Library Match** (or **Alt+1**) to still have access to the PSMs supported each imported peptide
- Select a peptide, right-click on the **Library Match** window and on the **Ion Types** options make sure the **Losses** of the defined modifications are selected

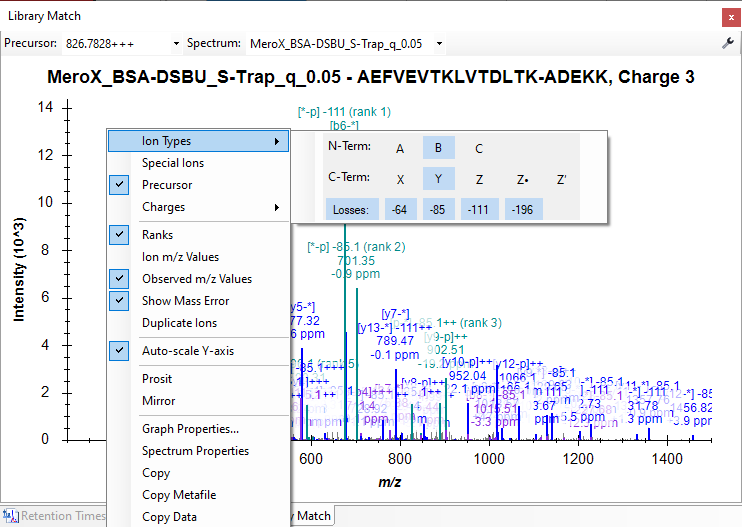

NOTE: at the moment Skyline will import all peptides into a single Library Peptides list since there is no established nomenclature on how to manage peptide-protein associations for this type of peptides. Nevertheless, if you would like to split the peptides into lists of interest this can be done manually by dragging the peptides into other peptide lists. It is recommended to do this before you import your raw data.

#### Importing raw MS data

Now that the settings, including the ion mobility library, and the targets have been defined we can import our DDA-PASEF results:

- Click on **File🡪Import🡪Results…**
- Select **Add single-injection replicates in files**
- Allow for **Many** **Files to import simultaneously** and make sure to tick on **Show chromatograms during import** (that way you can see some nice Skyline TV as known in the community)
- Click **OK** and select all the raw .d Bruker files provided with this tutorial
- Click **Open** and then the EICs should start being generated

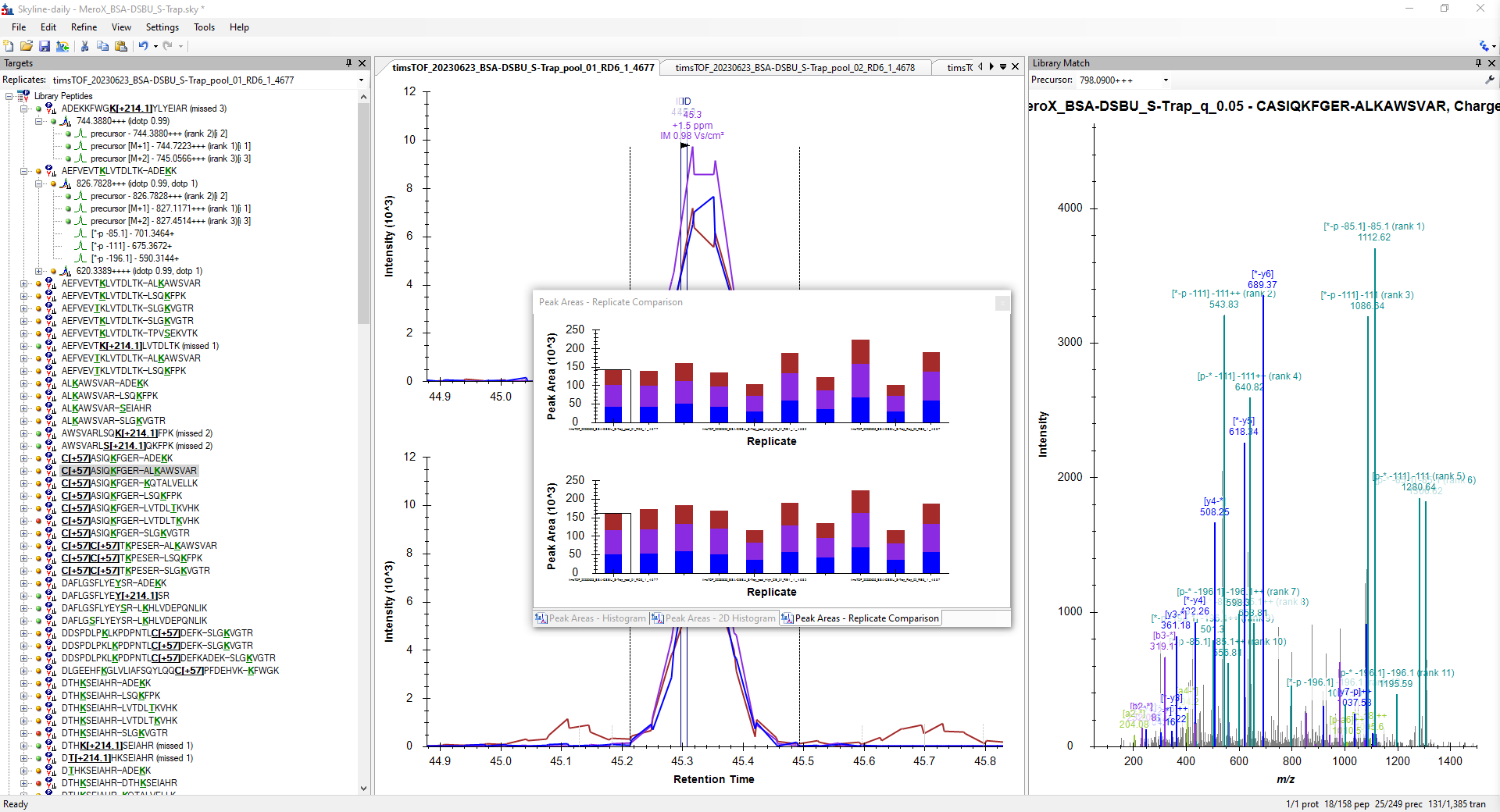

#### Manual correction of ion mobility library

Before having a look at some examples of cross-linked peptides, we have to first correct some of the estimated values in the ion mobility library due to the presence of gas phase conformers.

First, make sure the MeroX_IMS_Library.txt is loaded into an Excel sheet. Now, let’s have a look at two examples to have an idea of what would happen if these examples are not corrected:

Example #1: The ion that had the largest ion mobility range when estimating its average ion mobility was the [M+5H]^5+^ ion of the cross-link KVPQVSTPTLVEVSR-HKPKATEEQLK-[+196.084792@1,4] with a range of 0.195. Do the following to have a look:

- On the **Replicates** list make sure **timsTOF_20230623_BSA-DSBU_S-Trap_pool_01_RD6_1_4677** is selected
- Search for **K**VPQVSTPTLVEVSR-HKP**K**ATEEQLK on the target list
- Left-click on the left **plus** symbol to expand the precursor ions tree
- Left-click on **629.5543, +5** corresponding to the [M+5H]^5+^ ion

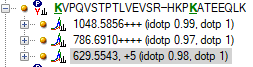

- Now on the chromatogram window click on the apex of the chromatographic peak to have a look at the raw ion mobility spectrum

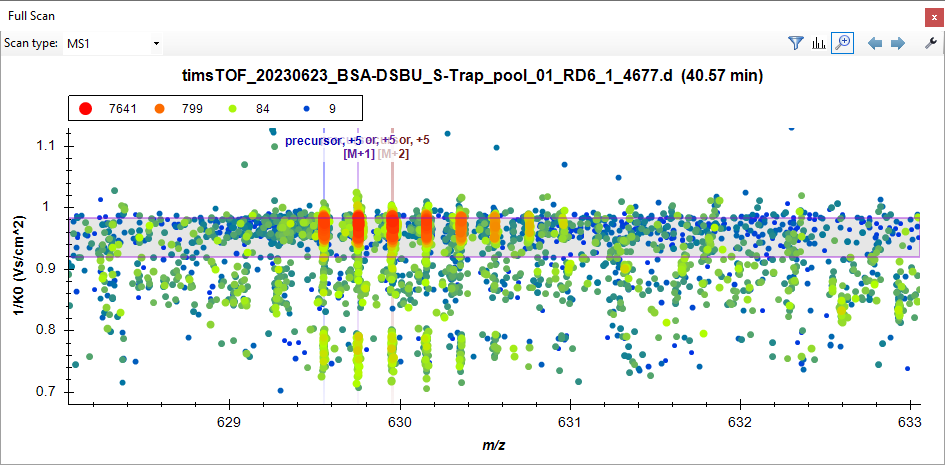

NOTE: Make sure that the **Show 2D Spectrum** icon is unticked so you have access to the 3D representation of the ion mobility spectrum with *m/z* set as the x-axis, the ion mobility as the y-axis, and the ion intensity as the z-axis (with a logarithmic scale color coding).

In this case the center of the ion mobility extraction window is at 0.9520 which is offset from the center of the most abundant conformer with a 1/K_0_ at around 0.968. There were 12 PSMs for the most abundant conformer and 1 for the lower intensity conformer at around 0.77, causing the average to be heavily weighted towards the most intense conformer. The extraction window managed to capture most of the main (top) ion population distribution, but you can see that the offset caused some of it to be missed. Now let’s correct the library entry for this precursor:

- Open an Excel sheet with the MeroX_IMS_Library.txt loaded in it
- Sort the entries first **by Modified Sequence** name and then by **charge**
- Find the row corresponding to **KVPQVSTPTLVEVSR-HKPKATEEQLK-[+196.084792@1,4]** with **charge 5**
  - The current ion mobility value should be **0.952**
- Now go back to the ion mobility spectrum, place your mouse icon left of the y-axis and at the approximate center of the ion population distribution
- Use the **middle mouse scroll-button** to zoom in the center until you can approximate the mobility value to the third decimal
  - We will set this value as **0.968**
- Go back to the Excel sheet and replace the **Ion mobility** entry for the discussed precursor

You could add an additional entry in the IMS library corresponding to the conformer around 0.77 for this charge state, but at the moment Skyline will only integrate one gas phase conformer per peptide isomer so it would be ignored. For the purpose of this tutorial just choose the most intense conformer (first criteria) with the highest 1/K_0_ (secondary criteria) as the library entry.

(Optional) We find it good practice to double check the corrections to the IMS library after the results are re-imported. You can leave a precursor note on the precursor ions for which you corrected the ion mobility library entry. **Right-click** on the selected precursor, **left-click** on **Edit Note**, and write a text tag that you can search afterwards in the **Document Grid** (e.g., re-IMS).

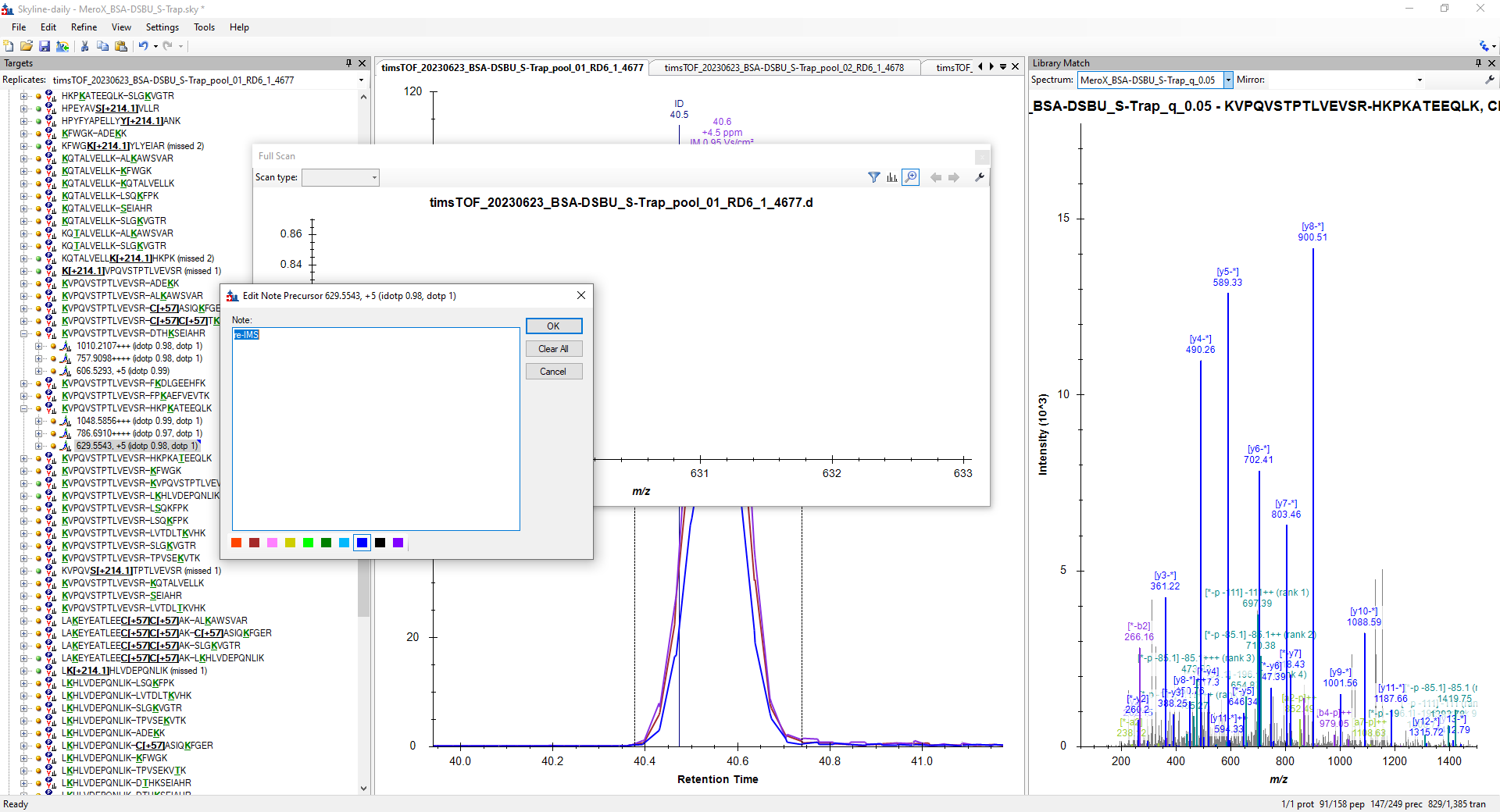

Example #2: The [M+4H]^4+^ ion of the cross-linked peptide KVPQVSTPTLVEVSR-DTHKSEIAHR-[+196.084792@1,4] has a range of reported mobility values of 0.098. Do the following to have a look:

- On the **Replicates** list make sure **timsTOF_20230623_BSA-DSBU_S-Trap_pool_01_RD6_1_4677** is selected
- Search for **K**VPQVSTPTLVEVSR-DTH**K**SEIAHR on the target list
- Left-click on the left **plus** symbol to expand the precursor ions tree
- Left-click on **757.9098, ++++** corresponding to the [M+4H]^4+^ ion
- Now on the chromatogram window click on the apex of the chromatographic peak to have a look at the raw ion mobility spectrum

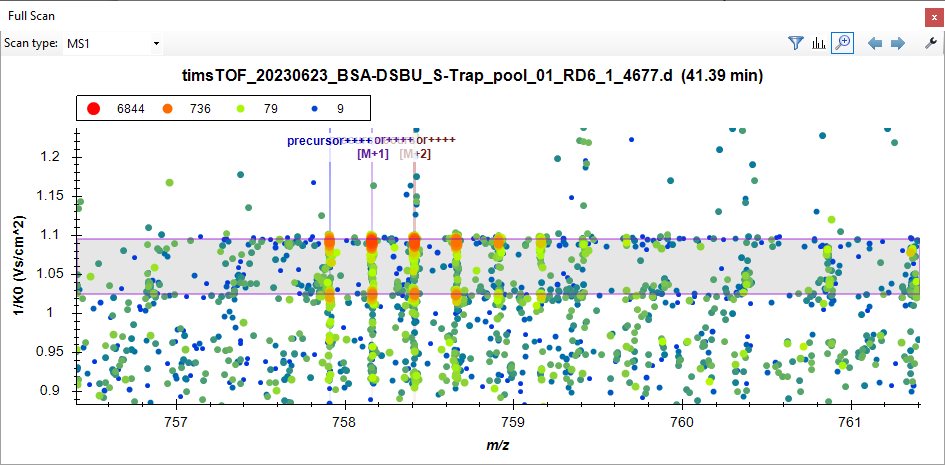

In this case the simple average estimation failed in approximating the center of either of the two gas phase conformers and caused an incorrect integration of either of the examples. There were 7 PSMs with a 1/K_0_ around 1.01 and 10 PSMs with a 1/K_0_ at around 1.09 causing the average to be somewhere in the middle of both ion populations. Now let’s correct the library entry for this precursor:

- Open an Excel sheet with the *MeroX_IMS_Library.txt* loaded in it
- Sort the entries first **by Modified Sequence** name and then by **charge**
- Find the row corresponding to **KVPQVSTPTLVEVSR-DTHKSEIAHR-[+196.084792@1,4]** with **charge 4**
  - The current ion mobility value should be **1.060647059**
- Now go back to the ion mobility spectrum, place your mouse icon left of the y-axis and at the approximate center of the ion population distribution
- Use the **middle mouse scroll-button** to zoom in the center until you can approximate the mobility value to the third decimal; choose the most intense conformer with the largest 1/K_0_
  - We will set this value as **1.092**
- Go back to the Excel sheet and replace the **Ion mobility** entry for the discussed precursor

Assuming there were enough points for all gas phase conformers then these could be clustered and the average could be extracted from each cluster of conformers, but unfortunately some of the conformers were only sampled once in this dataset (as in the first example shown). Therefore, we have shown you how to correct these values manually.

For new samples or re-measurements with new IMS conditions you would repeat the process above for precursor ions that have ion mobility ranges above the binning window (Δ1/K_0_ = 0.025 for this data set) used during *compound creation* in DataAnalysis to double check if errors introduced by averaging would affect integration. However, for this tutorial we have provided a corrected IMS-library (i.e., *MeroX_IMS_Library_corrected.txt*) with the contents of this tutorial. Load the corrected library:

- Click on **Transitions Settings…** under the **Settings** menu
- Go to the **Ion Mobility** tab
- On the **Ion mobility library** drop-down menu click on **<Add…>**
- Set the **Name** as **MeroX_DSBU-BSA_166ms_corrected** and click on **Create…**
- Save the new IMS-library in the same folder as the Skyline document
- Load the *MeroX_IMS_Library_corrected.txt* file into an Excel sheet
- **Copy** the cells of first 5 columns
  - IMPORTANT: avoid copying the column titles
- Back on the **Edit Ion Mobility Library** under the **Measured peptides** table, left-click on the upper left corner box (highlighted in blue below) and paste the copied cells (**Ctrl+V**)
- Click **OK** twice until you are back on the main Skyline window
- Save the document (**Ctrl+S**)
- Click on **Manage Results…** (**Ctrl+R**)

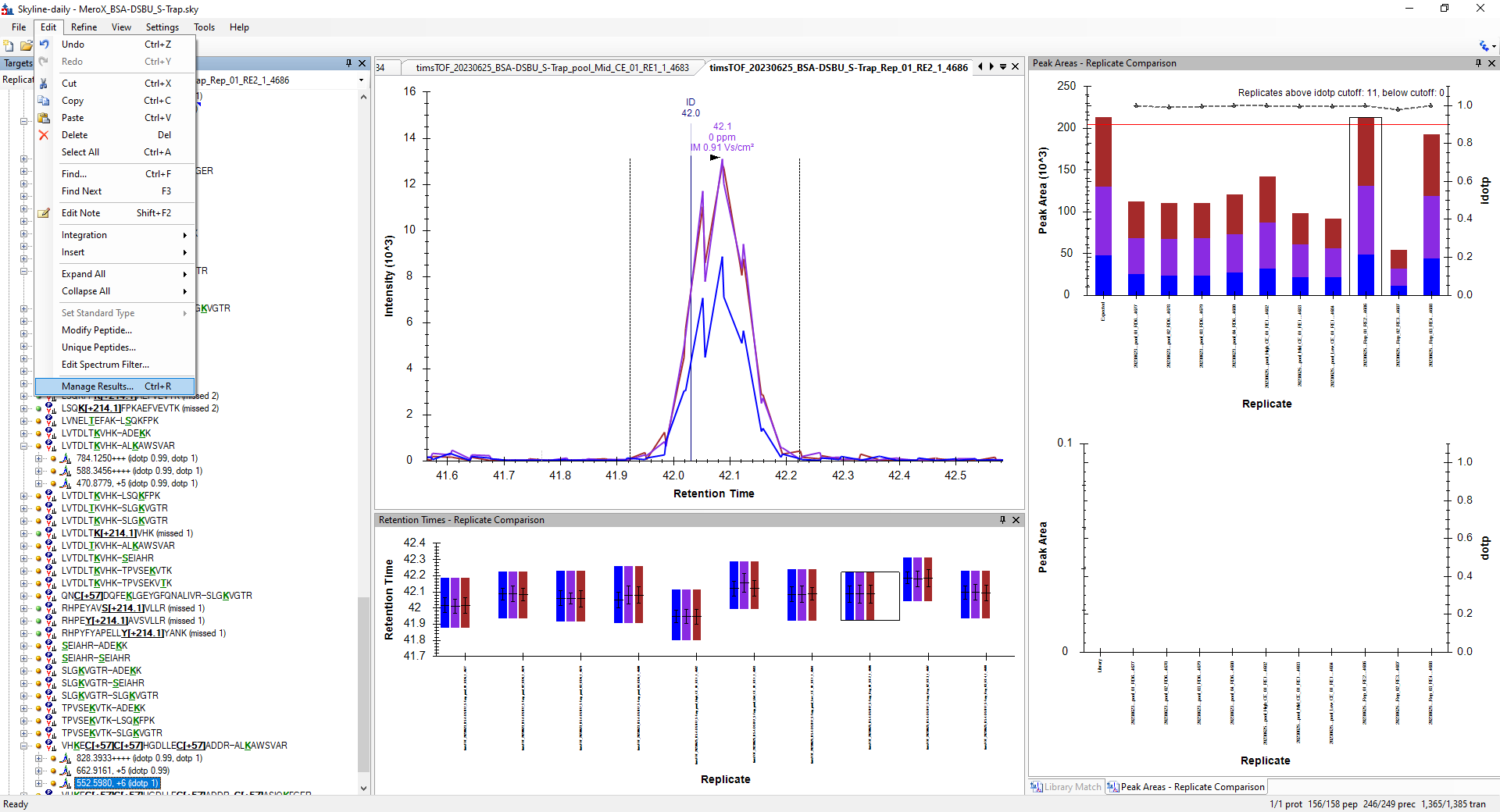

- **Select all Replicates** and then click on **Re-import**
- Once an asterisk appears left to the replicates name, click **OK** to re-generate the EICs

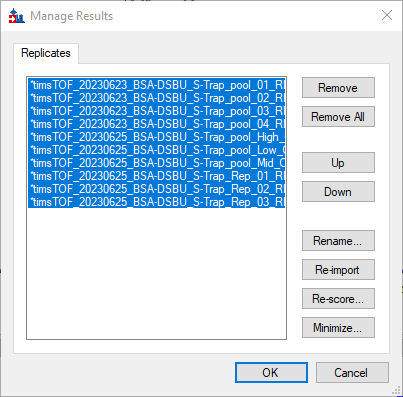

- Once the chromatograms have been re-generated save the document

#### Fitting XL-peptides to retention time calculator

One of the challenges of identifying cross-linked peptides is that the peptides generated by adding a cross-linker reagent exist in substoichiometric quantities and tend to be just a small portion of the peptide mixture that is analyzed by LC-MS. However, the fact that cross-linked peptides are only observed upon the addition of reagents can be used to identify unique low intensity peptide features within a peptide digest when compared to reaction negative controls (i.e. same matrix without cross-linker addition). Additionally, under reproducible chromatographic conditions these negative controls can be used as an additional layer of verification of false positive IDs by checking for the presence of “true IDs” in a matrix where these should not exist.

The samples presented in this data set were spiked with Pierce iRT peptides. This will help in correcting for any small deviations in the chromatography of the samples measured. Let’s add first the iRT peptides:

- Click on **Peptide Settings** under the **Settings** settings menu
- Select the **Prediction** tab
- Under the **Retention time predictor** list select **<Add…>**
- On the **Edit Retention Time Predictor** window name the predictor as “Pierce_BSA-DSBU”
- On the **Calculator** list select **<Add…>**
- On the **Edit iRT Calculator** window **name** the predictor as “Pierce_BSA-DSBU_iRT-Lib” and click on **Create…**
  - This will create the iRT peptides database file (*.irtdb*) that contains the indexed retention times of your peptides
- **Save** the database with the name “Pierce_BSA-DSBU_iRT-Lib” on the same folder containing the Skyline document
- Back in the **Edit iRT Calculator** select **Pierce (iRT-C18)** from the **iRT standards** list
- Tick the **Store redundant iRT values**

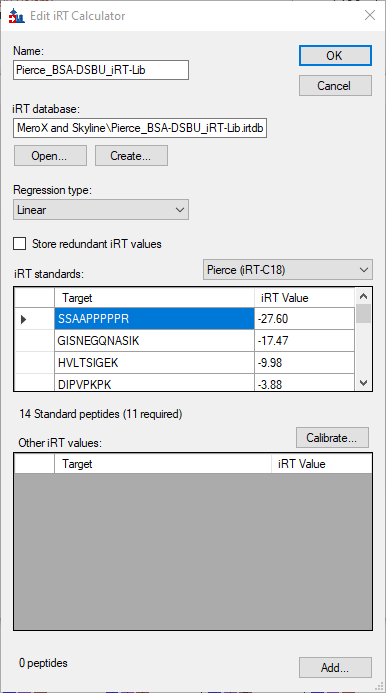

- Click **OK**
- Back on the **Edit Retention Time Predictor** set the **Time window** to **5 min** and then click **OK** twice until you are back to the main Skyline window
- Skyline will recognize that you do not have the iRT standards in the document and will ask you if you would like to add them. Set the **Maximum transitions per peptide** and click **Yes.** This should immediately add the first three isotopes of the respective precursor ions, i.e. monoisotopic ion (M), the ion with one C_13_ isotope (M+1), and the ion with two C_13_ isotopes.

Now you should have a new set of targets in the document corresponding to the **Pierce standards**. Make sure the following precursor ions and corresponding three first precursor isotopes are selected:

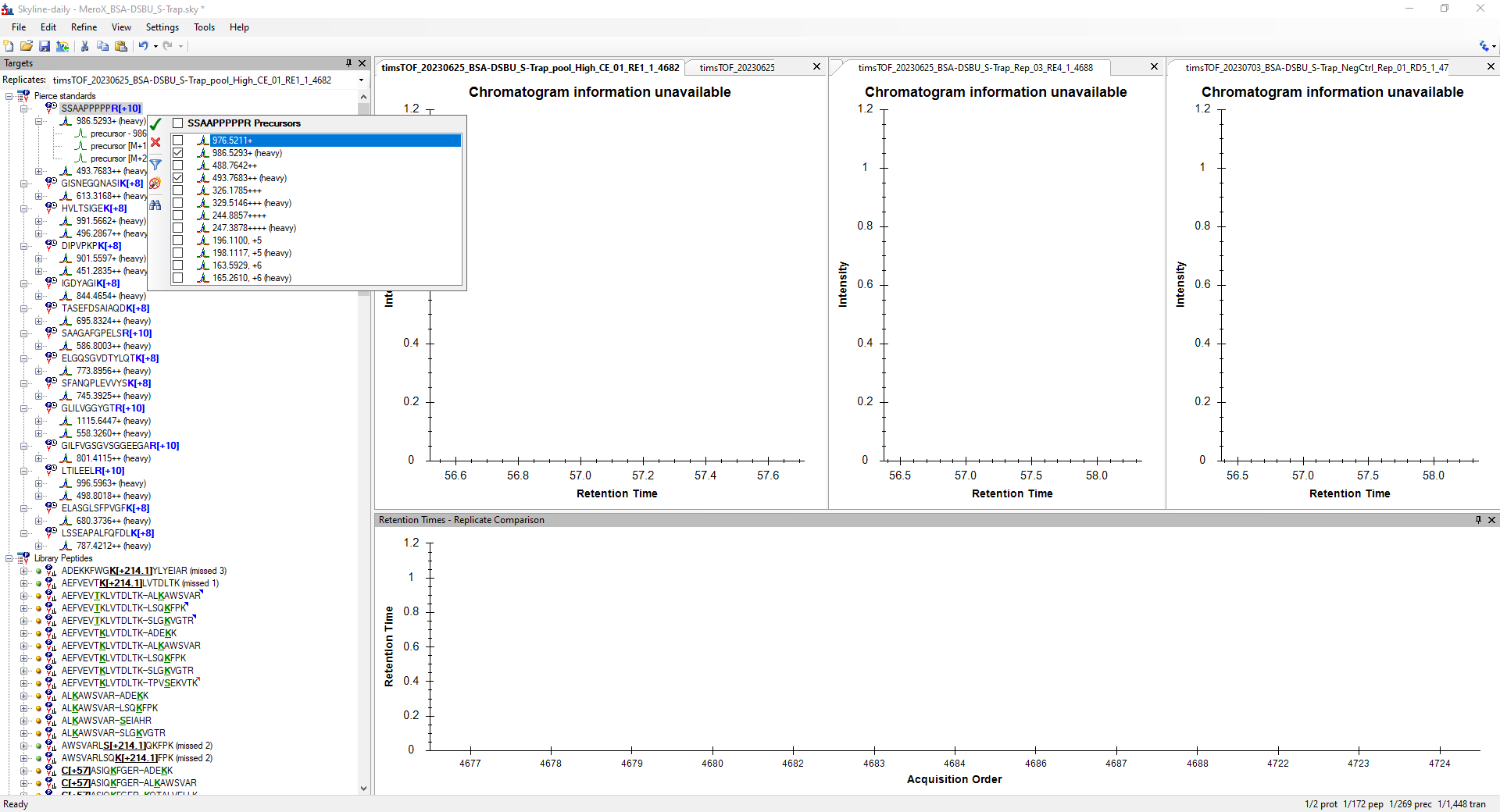

NOTE: If the targets you get are not the same as those in the figure, you can edit the targets by clicking selecting a peptide or precursor and pressing the **spacebar**.

Let’s re-generate the EICs to include these precursors:

- Click on **Manage Results…** (**Ctrl+R**) under the **Edit** menu
- Select all files and click **Re-import**

The retention times of these peptides were determined externally by measuring only the Pierce Peptides. The detectable precursors of these peptides were not sampled during the analysis of the BSA-DSBU digests. To maximize detection of cross-linked peptides, the MS method was designed to exclude precursor ions with charge states lower than 3 to avoid oversampling unmodified peptides. Thus, the singly and doubly charged precursor ions of the Pierce Peptides were not sampled during DDA and could not be identified based on their MS/MS spectra.

To make sure that you use the same retention times defined by the external measurements we have provided the corresponding peak boundaries for these peptides in the tutorial dataset. To enforce these peak boundaries do as follows:

- Open on the **File** menu
- Click on **Peak Boundaries…** within the **Import** options

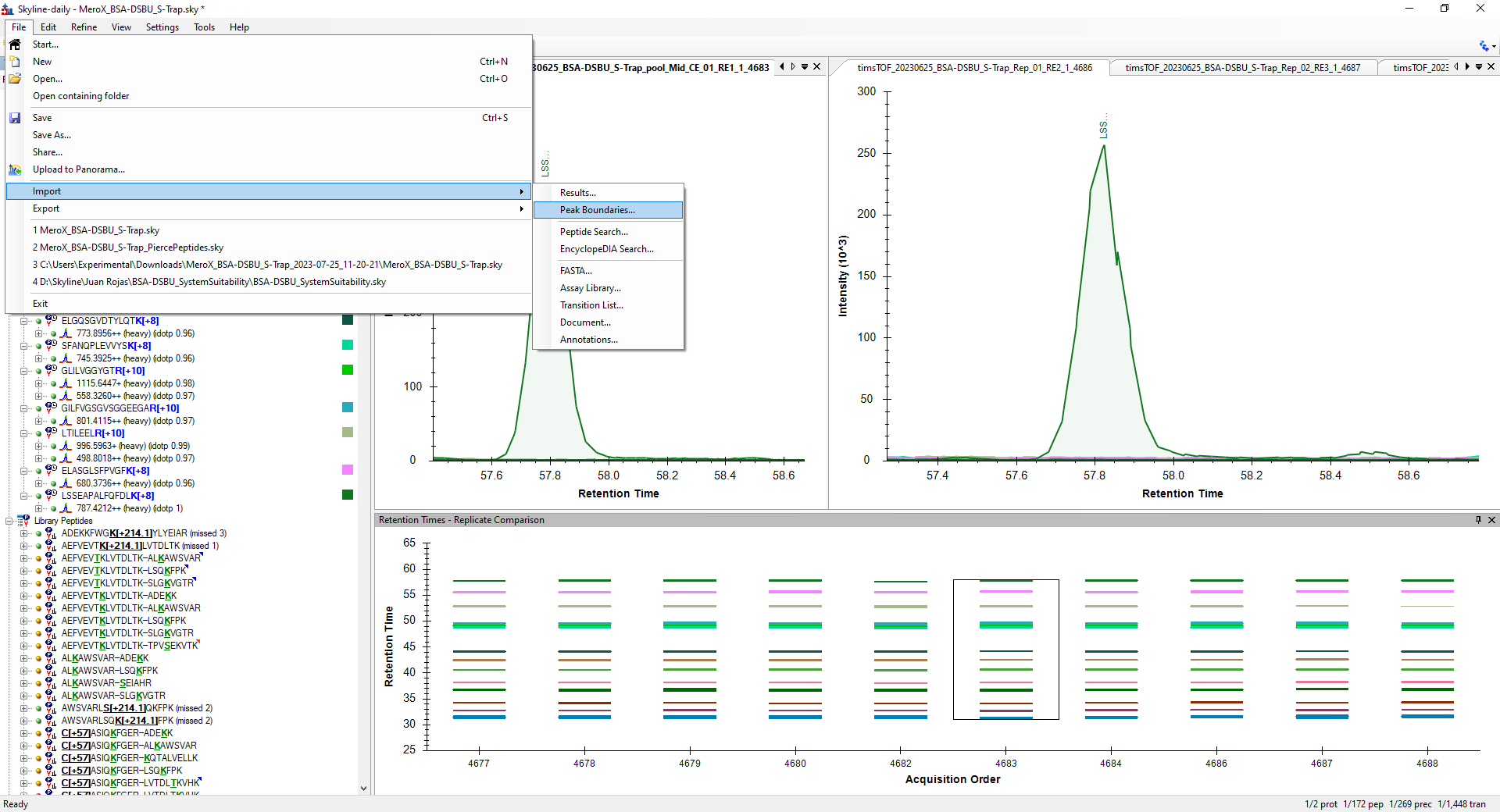

- Select the “Pierce_Peptides_Peak_Boundaries.csv” file included with the tutorial material
- **Right-click** on the chromatogram window and make sure **Retention Time Prediction** is chosen

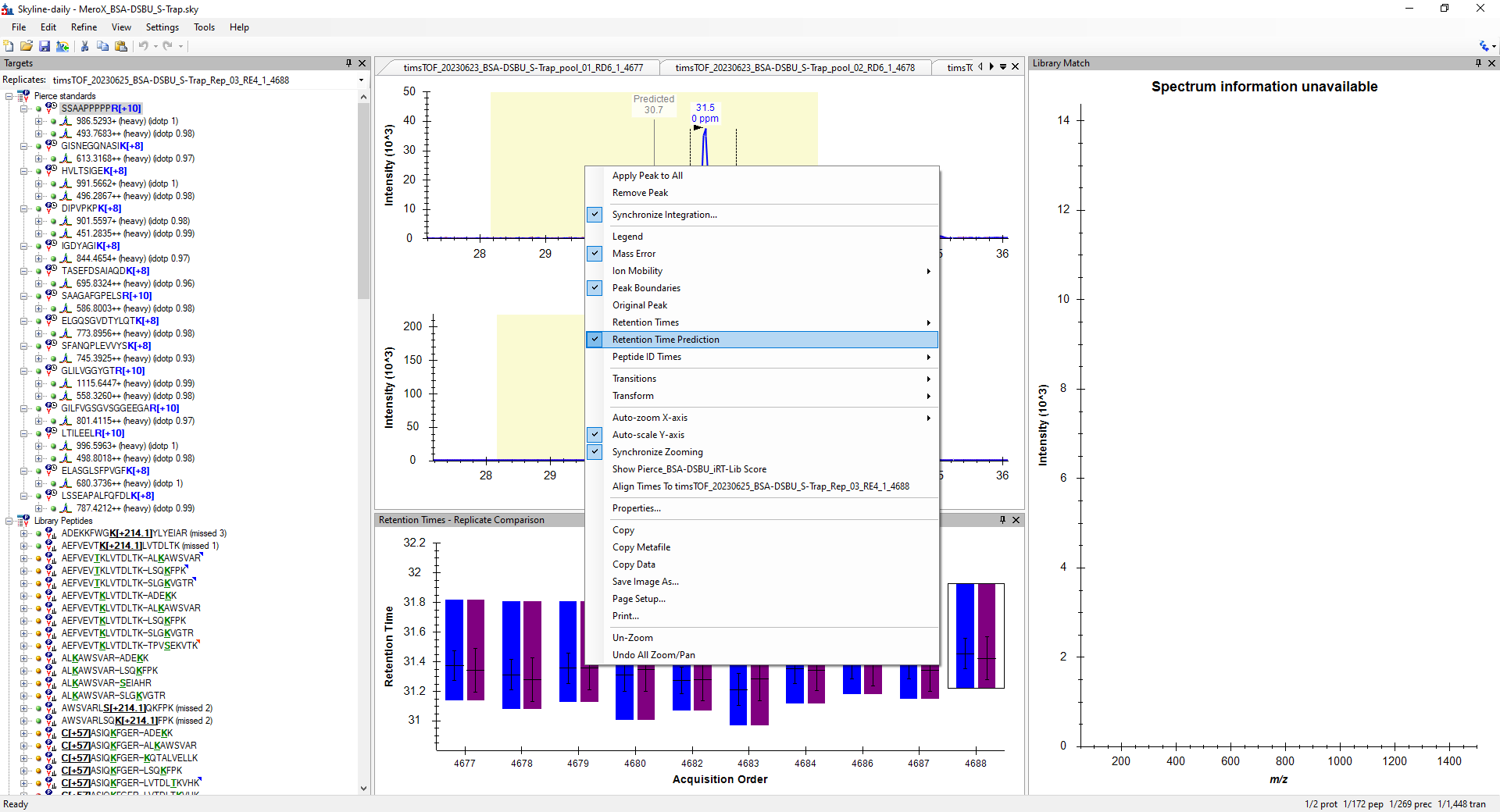

Let’s have a look at the re-imported data. If you click on the iRT standard peptides you will notice that a yellow background is displayed. This background corresponds to the retention time window prediction that we selected before. However, if you click on any other peptide that are not the iRT peptides this window yellow background will not be present. This is because these peptides have not been indexed yet on the retention time calculator. To fit the rest of the peptides to the retention time calculator do as follows:

- Click on **Peptide Settings** under the **Settings** menu
- Go to the **Prediction** tab
- Click on the **calculator icon**
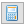
 and then click on **Edit Current…**
- Once you are on the **Edit iRT Calculator** window click on **Add…** button in the bottom right and then select **Add Results…**

Now you should have the **Add iRT Peptides** window with a table containing each of the replicates and the result of attempting to fit the results to the calculator.

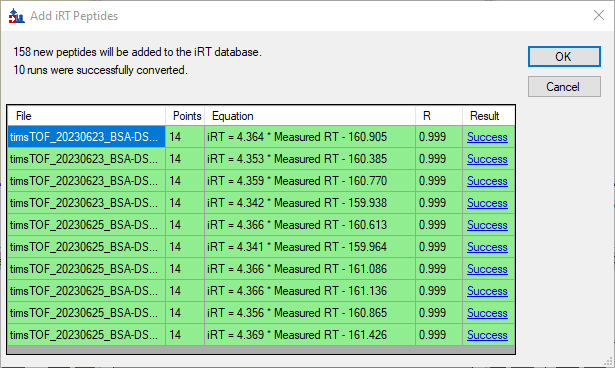

The example above was done with a refined version of the Skyline document you have created. For the purpose of this tutorial we will click **Cancel** so you can work with a more extended iRT database that was created with a larger dataset analyzed with the same chromatographic conditions.

NOTE: Here we are not attempting to predict the retention time of the identified peptides based on intrinsic properties of the molecule. Instead, we are fitting the empirically determined retention times into retention time index defined by the standard peptides used. Therefore, entries to this iRT database should be managed carefully and additions to the database should be entered once the chromatographic peak selection has been revised to ensure best performance.

- After clicking **Cancel** go back to the **Edit iRT Calculator** window and click on **Open…**
- Select the “Pierce_BSA-DSBU_Col20230608.irtdb” and click **Open**
- Once the **Other iRT values** has been populated click **OK** twice until you are back to the main Skyline window

Now, if you click on a cross-linked peptide, e.g. **C[+57]**ASIQ**K**FGER—ADE**K**K, you will notice that the yellow background and the predicted retention time is visible. Let’s have a look at how all peptides fit to the retention time prediction at the same time:

- Click on the **View -> Retention Times -> Regression -> Score To Run**

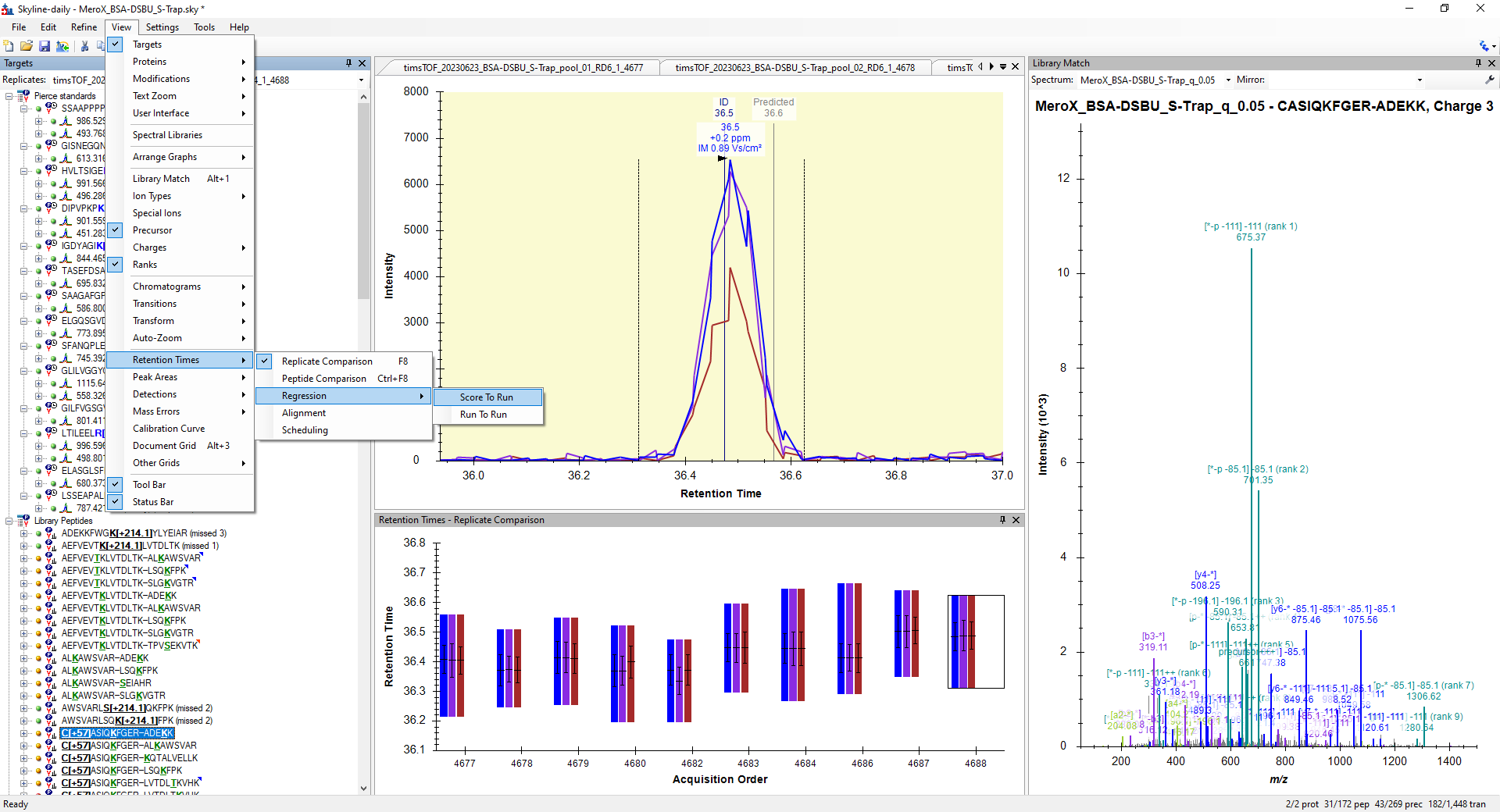

- Once **Retention Times - Score To Run Regression** window opens, **right-click** on it and make sure that the **Calculator** selected is **Pierce_BSA-DSBU_iRT-Lib**

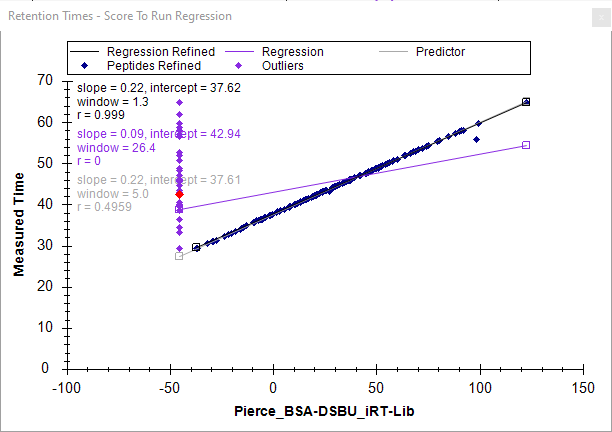

All peptides in the document that matched an entry in the calculator have a dark blue entry, while those that are not present in the iRT database are purple diamonds and are only displayed with their measured time. Before we have a look at some of the missing peptides in the iRT database let’s make sure we are all working on the same document by downloading a refined Skyline document that has been prepared for this tutorial.

- Go to <https://panoramaweb.org/XL-MS_MeroX_Skyline.url>
- Click on the **download** next to the of the “MeroX_BSA-DSBU_S-Trap_Refined” Skyline document in the **Targeted MS Runs**

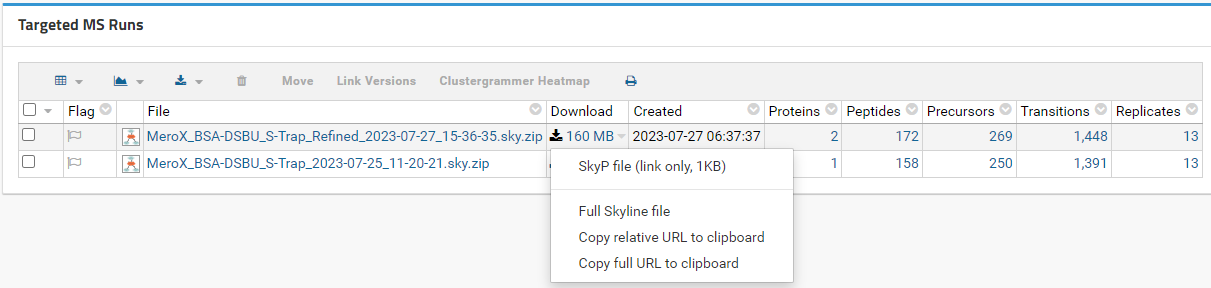

- Click on **Full Skyline file**
- Extract the contents of the compressed folder into the directory containing the raw MS Files

Now you have a Skyline document with adjusted peak boundaries (where necessary) and includes the Negative Control samples where the peak boundaries were forced onto the expected retention times of the cross-linked peptides.

#### Using chromatography and ion mobility features to identifying false positive IDs and resolve cross-linked peptides isomers

Fragmentation spectra of cross-linked peptides is inherently very complex due to the combination of peptide fragments generated from the linked peptide chains. There will be instances where only relying on the fragmentation spectra might not be enough to unequivocally identify the cross-linked peptides and their linkage sites by the automatized decision making of database search engines. Some of these instances include, but not limited to: 1) co-fragmentation of isobaric, co-eluting ions produce chimeric spectra difficult to identify, 2) low intensity fragment ion spectra generated from low intensity precursor ions peptides generated from transient interactions are not easily detected in the conditions implemented, 3) presence of cross-linked peptide isomers that produce fragment ion spectra that support more than one linkage position. Ultimately, the MSMS spectra is only part of the information collected for peptides analyzed by liquid chromatography coupled to ion mobility spectrometry and tandem mass spectrometry (LC-IMS-MS/MS). Skyline allows visualizing within the same platform many intrinsic properties of the separated peptides, including the collected fragmentation spectra, that allows adding additional layers of verification for cross-linked peptides. Here will show three examples of an ideal scenario, an ambiguous scenario, and a challenging scenario that would require re-sampling with targeted methods in additional experiments to verify the XL-peptide ID.

##### The Good

F**K**DLGEEHFK—SLG**K**VGTR

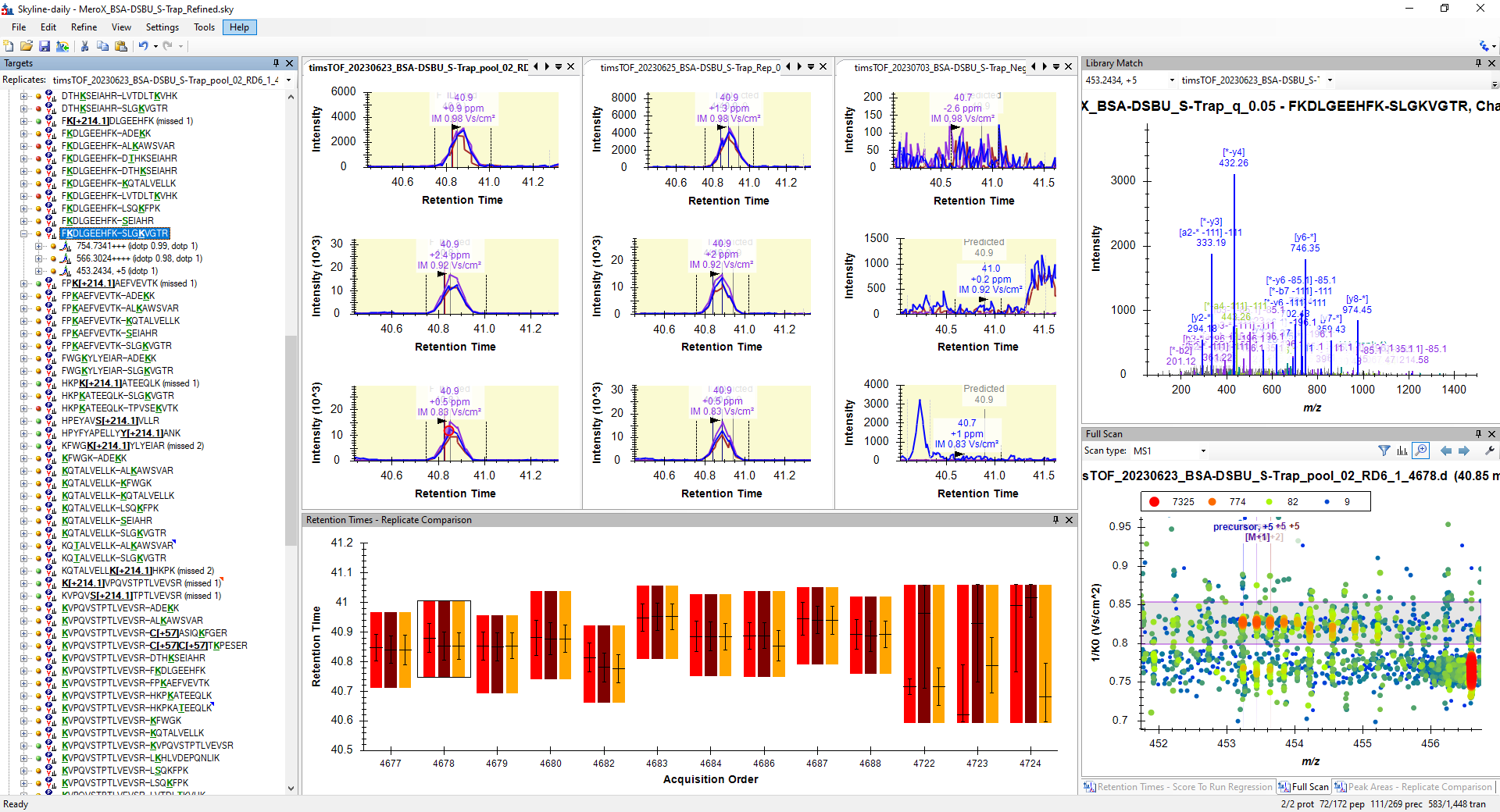

The XL-peptide F**K**DLGEEHFK—SLG**K**VGTR-[+196.084792@2,4] was identified by DDA-PASEF with three distinct precursor ions with interpretable MS/MS spectra for each. These can be scanned through the **Library Match** window (shown on the top-right). As you scan through this Skyline will point out to the sample from which these were collected; for this dataset they are usually around the apex since DataAnalysis does chromatographic peak picking during the MS/MS data binning. It is good to check the origin of the MS/MS IDs and make sure they match to a sensible chromatographic peak shape. The overlapping chromatographic traces of the precursor ions further support the identity of a true peptide feature. For each precursor, three isotope ions are automatically displayed in the chromatographic traces and a relative ratio between each isotope's integrated areas and their expected elemental composition isotopic distribution is calculated. This ratio is referred to as the **isotopic dot product** (**idotp**) and is displayed to the right of the precursor ions in the target list; there is no clear threshold for this value, but values below 0.9 should be investigated for what might cause deviations from 1. The **idotp** is useful to identify wrong precursor monoisotopic peak selection or detection of integration interference from co-eluting, isobaric species.

In the bottom right, is a display of the ion mobility spectrum of the [M+5H]^5+^ ion with *m/z* at 453.2434 at the peak apex of the “S-Trap_pool_02” instrument technical replicate. Here we can see how ion mobility filtering during MS/MS spectra binning facilitated the ID of this precursor. The quadrupole isolation width was set to 2 m/z from the center of the selected precursor. In the presented example this isolation width would have included at least part of the fragment ions generated from the neighboring precursor ion with monoisotopic *m/z* around 454.2. However, these ions have enough of a difference with respect to their ion mobility that the contributions from the neighboring precursor ion could be filtered out and avoided creating a chimeric MS/MS spectrum.

At the bottom of the screen you can see the chromatographic peak boundaries for all replicates included in this document, including the peak boundaries enforced on the negative controls (last three). The stable chromatographic conditions help in confirming the detection of the XL-peptide in all XL replicates,despite some of its precursors having missing PSMs. At the same time, the chromatographic stability can be confirmed in the negative controls by using the iRT peptides and accurately predict the expected retention time of the indexed XL-peptides in the negative controls. By enforcing the peak boundaries around this predicted retention time we double check that proposed XL-peptides are not a false ID produced by wrongly assigning other peptides from the digest matrix. For this XL-peptide example, the sequence coverage was complete and the peptide’s precursor ions intensities were overwhelmingly above the background signal that was easy to believe in its identity without this last validation step; you can check this by looking at the **Peak Areas - Replicate Comparison** graph. However, we have found this additional quantitative criterion useful to support the identity of XL-peptides that are borderline detectable that we can then target on tailored PRM experiments to confirm their identity.

##### The Confusing

**C[+57]**ASIQ**K**FGER—LVTDL**T**KVHK

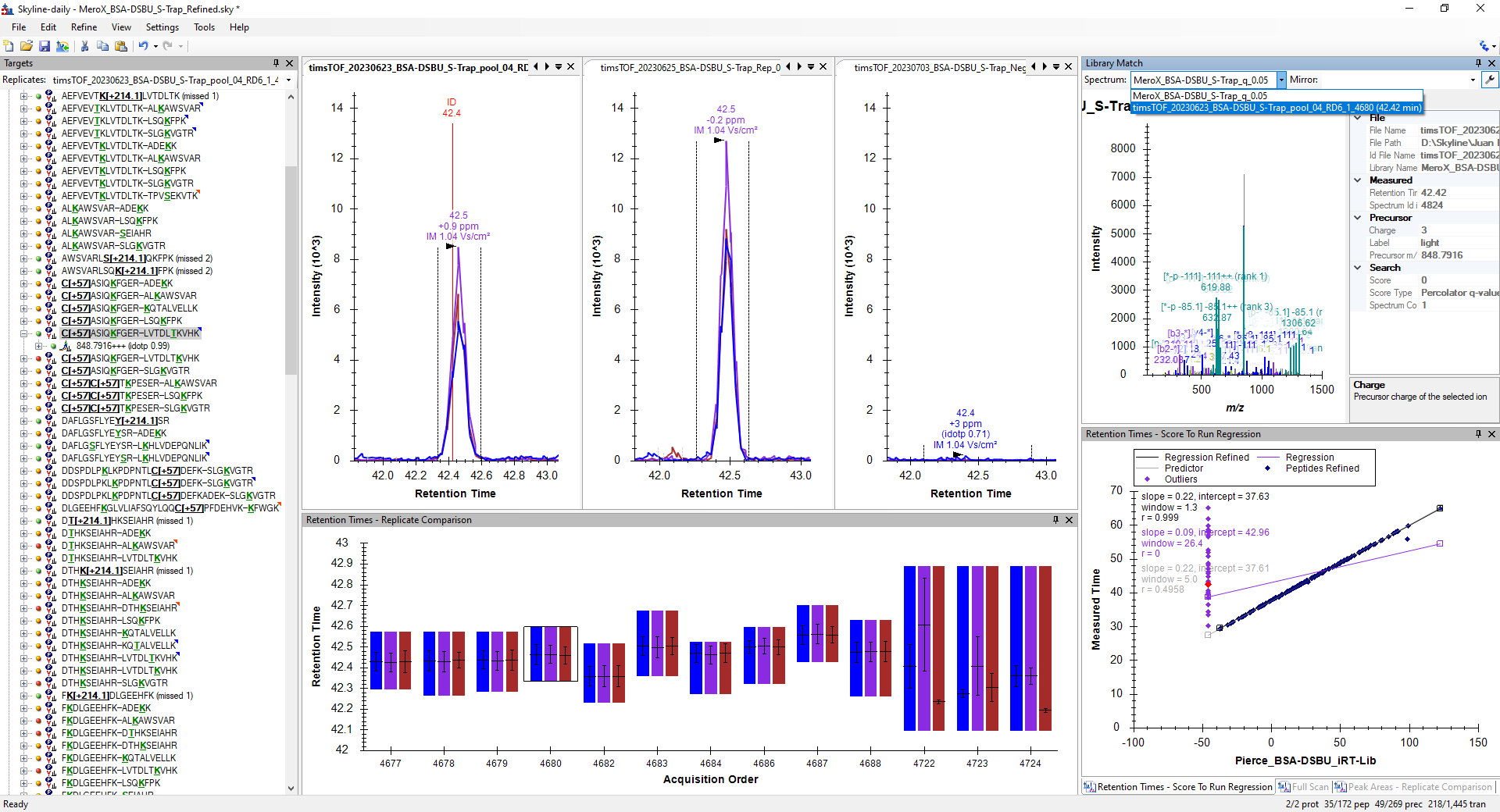

The XL-peptide C[+57]ASIQKFGER—LVTDLTKVHK-[+196.084792@6,6] was identified by DDA-PASEF with only one PSM from the [M+3H]^3+^ precursor ion. This can be seen by looking at the list of Spectrum matches in the Library Match window or by checking on the details of the spectrum by clicking on the wrench
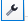
symbol on the top-right corner of the Library Match window. However, the peptide feature is unique from the XL samples indicating it was generated during the DSBU reaction and the peptide sequence coverage is almost complete. This is an example of peptide identification where the cross-linked sequences are correct, but the cross-linking site was incorrectly assigned for this PSM. Close inspection of this PSM in the MeroX GUI showed that equal probability of assignment was assigned to the T6 or K7 amino acid of the right peptide. The spectrum quality does not allow unequivocally assigning the linkage site of one of the connected peptides. However, this peptide is also a good example of being able to have a look at the whole combined dataset instead of being hyperfocused on a single MSMS spectrum. If you look closely at the targets list, you will see that right below is the C[+57]ASIQ**K**FGER—LVTDLT**K**VHK-[+196.084792@6,7] XL-peptide. This peptide is supported by three charged states and a total of 24 PSMs. The apex retention times are the same and this can be further highlighted by clicking on both targets while holding **Ctrl** pressed:

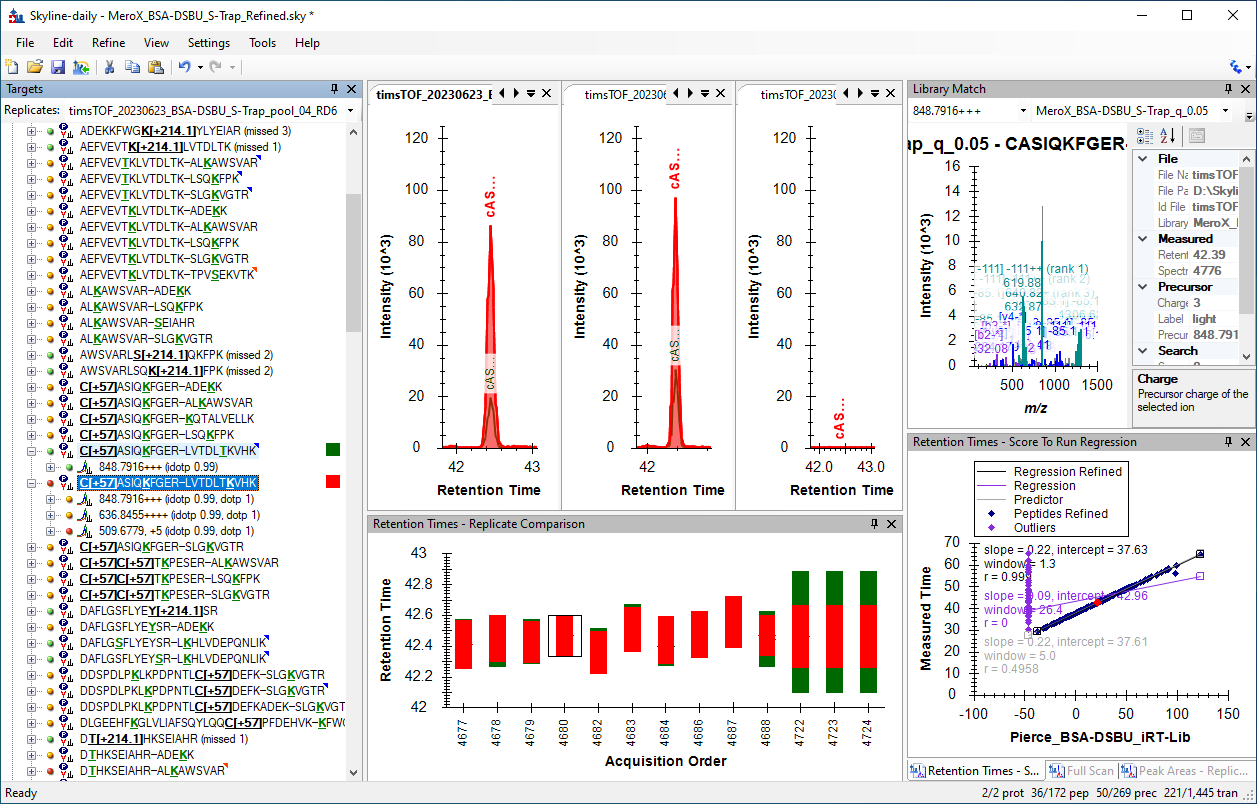

The other MSMS spectra acquired for these precursors allowed identifying correctly the XL-site of the right peptide. You can also notice that there is no predicted time for C[+57]ASIQKFGER—LVTDLTKVHK-[+196.084792@6,6]peptide in the chromatogram window and corresponds to one of the missing peptides in the iRT database (highlighted red diamond on the **Retention Times - Score to Regression** window). We have curated the iRT database provided to avoid having wrong entries so that problematic MSMS spectra like the one in this example can be more easily identified in future measurements.

##### The Ugly

L**K**HLVDEPQNLIK—LVTDLT**K**VHK

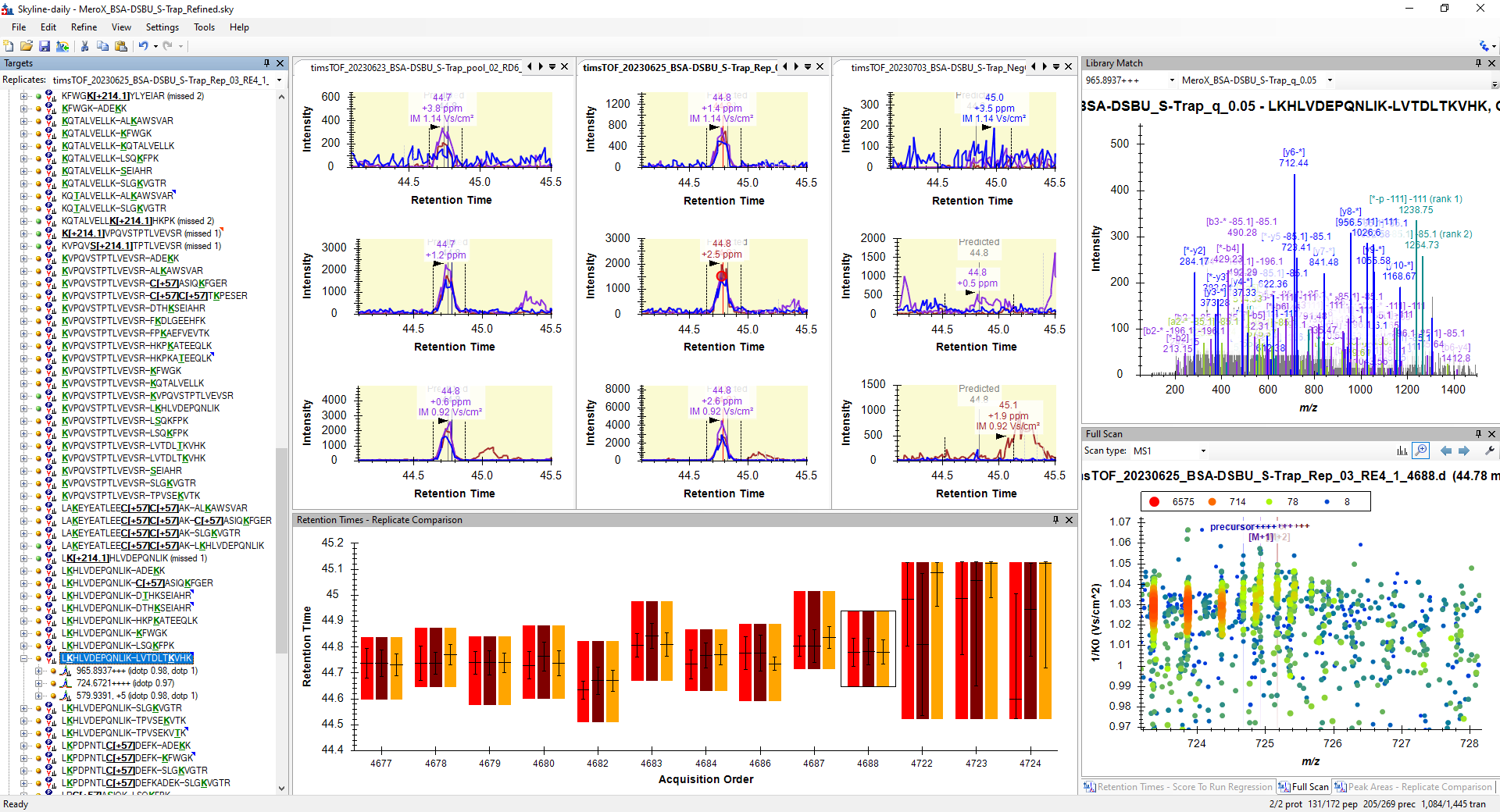

Finally let’s have a look at a challenging example with XL-peptide L**K**HLVDEPQNLIK—LVTDLT**K**VHK. This peptide was identified with two different precursor ions, the [M+3H]^3+^ and the [M+5H]^5+^ ions with a single PSM each. We found it curious that the [M+4H]4+ ion was not identified. Therefore it was manually included to generate EICs for this charge state as well. This is why in the refined document you will find all three charge states in contrast to those generated from the DDA results. By inspecting the ion mobility spectra at the apex of the [M+4H]^4+^ ion we can see the most likely reason for the missing ID, severe co-fragmentation from a co-eluting species. The high resolving power of the timsTOF allowed separating these species in the *m/z* dimension avoiding any integration interference. However, the ion mobility of these ions is too similar, preventing filtering out interfering fragment ions using the ion mobility dimension that most likely resulted in uninterpretable chimeric spectra.

Despite the fragment ions having lower S/N ratio than the other examples presented, closer inspection of the [M+3H]^3+^ ion PSMs in the MeroX GUI allows locating the XL sites and provides enough sequence coverage to verify both linked peptides. Additionally, the co-elution of multiple charged states with well fitting **idotp** makes us believe this peptide ID is correct, albeit borderline detectable. We would keep this peptide in a filtered Skyline document for generating a PRM-PASEF method with Skyline (**File -> Export -> Method…**) for subsequent measurements that ensure collecting as much fragment ion current from this precursor as possible. This would be recommended for verification of the lowest scoring PSMs identified with the initial DDA-PASEF results.

### MeroX and Skyline Data Integration

The final part of this tutorial will illustrate how we combined the data exported from MeroX and Skyline to perform a quality check of the LC-TIMS-MS/MS method used to acquire the data. For this we used custom R scripts that are documented in an accompanying R Notebook (RNotebook - XL-MS with MeroX and Skyline). This notebook has been provided in a RStudio format (.Rmd) and additional formats (.pdf, .html). The notebook and required data to execute it can be found in the “**Supplementary Files: R Script and Data Integration**“ folder found in XXXXXXXX.

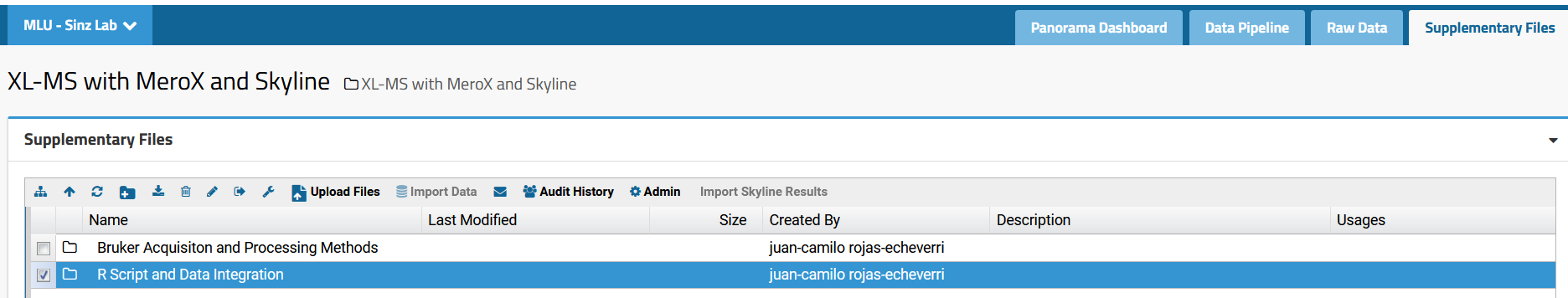

This portion of the tutorial can be followed with the corresponding R code used to generate some of the images presented here and if you are familiar with this programming language we would encourage you to read this portion in the interactive R Notebook mentioned above. Otherwise, you can read the main discussion around the images generated here.

#### MS/MS binning parameters selection

The **Processing and binning MS/MS scans with Bruker Compass DataAnalysis** section of the tutorial discusses how selection of the correct binning parameters with respect to retention time, *m/z*, and 1/K_0_ is important to improve quality of MS/MS spectra while not inducing chimeric spectra of distinct gas phase ion populations. Both chromatographic peak width and dispersion of ions in the gas phase are abundance dependent. The *m/z* of the detected ions is more robust and very small *m/z* drifts are expected within the time frame of chromatographic peak elution as long as detector saturation is not exceeded.

In an ideal scenario, with the MS data pre-processing applied to this data set, there should have been one PSM per gas phase conformer for each file. This PSM will be a summed and peak picked spectrum combining all fragmentation events for a specific precursor across multiple PASEF frames.

For the initial selection of these parameters, we find it to be more useful to start with broad ranges. When chimeric events are encountered then investigate which parameter tightening might help in reducing the binning of multiple gas phase ionic species. Below we discuss some criteria to keep in mind when selecting these parameters.

##### Drift in the *m/z* dimension

For the timsTOF Pro instrument used to acquire the dataset presented here, a *m/z* drift tolerance of **0.015** was used as a permissive, but reliable value that allowed recognizing fragmentation events of the same precursor ions. This worked for the 3-to-50 90 min gradients used here. Shorter gradients, where more crowded peptide elutions are expected, might benefit from tightening this threshold.

##### 1/K_0_ width

The initial binning range on the 1/K_0_ dimension was chosen by manually inspecting some of the mid to low intensity cross-links identified in previous measurements and was set to **0.025** to help resolving gas phase conformers. This can be illustrated with the **[M+3H]^3+^** ion of the **C[+57.021464]C[+57.021464]TKPESER-SLGKVGTR-[+196.084792@4,4]** cross-linked peptide. The 3D ion mobility spectrum at the chromatographic peak apex in the replicate timsTOF_20230623_BSA-DSBU_S-Trap_pool_01_RD6_1_4677 shows the presence of two distinct gas phase conformers.

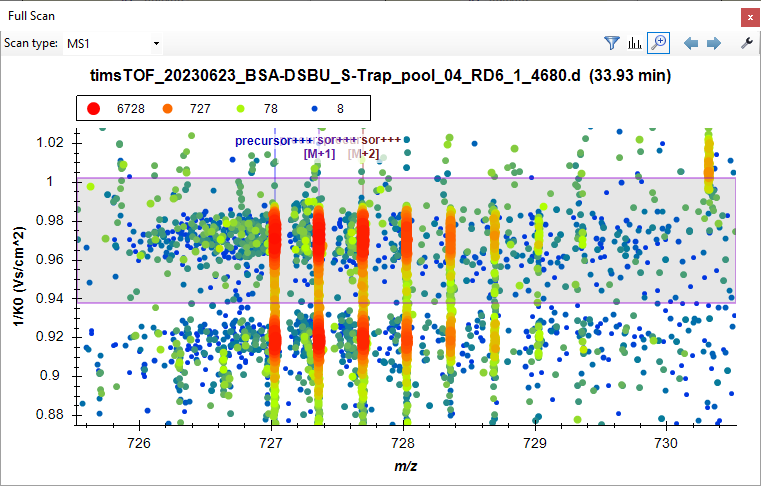

You will notice that there are two PSMs assigned to the same file with the exact retention time. These correspond to the conformers with 1/K_0_ at 0.920 and 0.959, respectively. The fragmentation pattern recorded was very similar, but not identical as displayed by dot product of the mirrored spectra of both spectra (image generated from the **Library Match** window in Skyline).

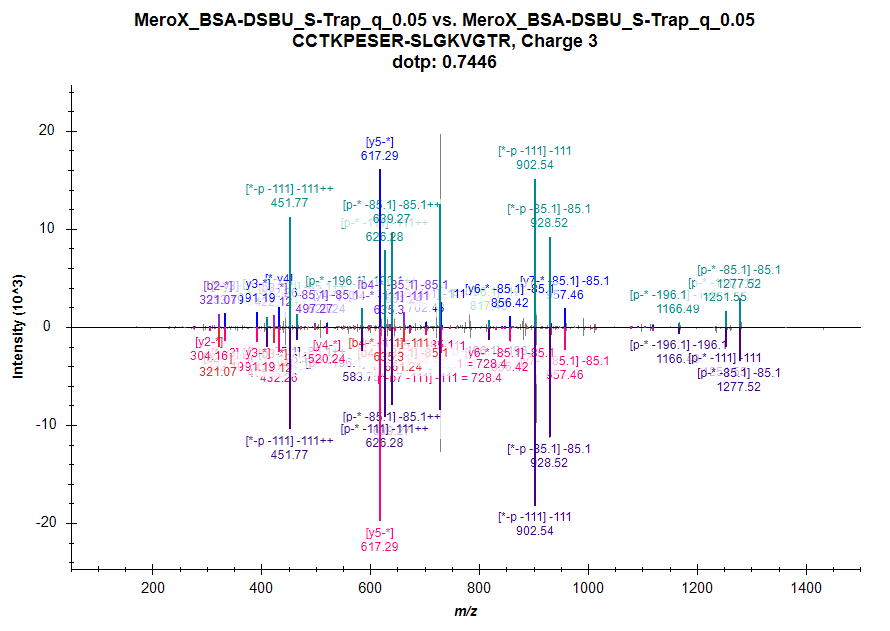

At first, we thought that the initial 1/K_0_ binning tolerance used might have been too strict for the most abundant ions, which have the widest ion mobility distribution. Let’s have a look at the **[M+5H]^5+^** ion of the **KVPQVSTPTLVEVSR-HKPKATEEQLK-[DSBU (XL)@1,4]** cross-linked peptide. You will notice that the 3D ion mobility spectrum at the chromatographic peak apex in the replicate **timsTOF_20230623_BSA-DSBU_S-Trap_pool_04_RD6_1_4680** shows an ion mobility distribution of 0.05-0.06.

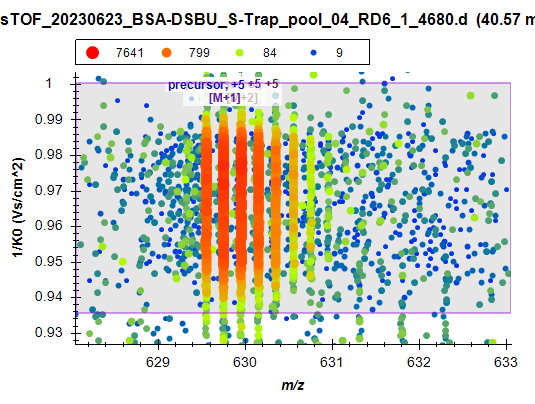

In the library match for this precursor ion, you will notice that there are two IDs assigned at the same time. Close inspection of the supporting spectra showed that the annotated 1/K_0_ for the supporting spectra were 0.976 and 0.950, which initially suggested fragment ion signal splitting due to the tight range of 0.025 for binning in the 1/K_0_ dimension. However, the ion mobilogram of this precursor ion (generated in Bruker DataAnalysis) across the chromatographic peak revealed that the split was due to the detection of two ion populations with very similar ion mobility, effectively detecting the shouldering ion mobility corresponding to the center of an additional gas phase conformer:

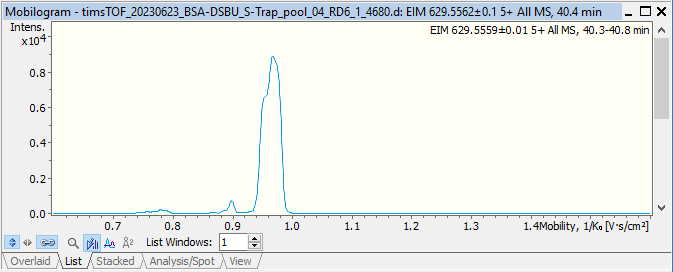

Even if there was fragment ion signal splitting, these would occur for the most abundant precursor ions where summation of fragment ion spectra across fewer PASEF frames would still give interpretable spectra.

Based on these evaluations across the detected cross-links, we concluded that binning in the ion mobility dimension with a 1/K_0_ range of 0.025 was a safe choice for separating information from gas phase conformers while increasing the S/N of fragment ions.

##### Chromatographic peak width

Chromatographic peak width is perhaps the most variable parameter across LC-MS systems of the three considered above for recognition of MS/MS events pertaining to the same precursor ion. The initial value was set based on prior measurements where we checked for the chromatographic peak width of mid to low intensity precursor ions. However, this parameter can be checked with some of the outputs provided by Skyline: **Max full width at half maximum (FWHM)** and the peak boundaries defined by the **Min Start Time** and the **Max End Time** which are calculated from the EIC of each precursor for all replicates in the document. A custom Skyline report (**Precursor Areas (XL).skyr**) that exports the above mentioned values can be found in the **“Supplementary Files: R Script and Data Integration”** of <https://panoramaweb.org/XL-MS_MeroX_Skyline.url>.

Here we use the Max FWHM calculated by Skyline which corresponds to the FWHM of the widest/most intense **transition** (i.e. ion target); in the case of DDA data this corresponds to the most intense precursor isotope. We use the curated peak boundaries as a proxy estimate of the baseline width of the chromatographic peak. We used these values to plot histograms of the recorded FWHM, the estimated baseline width, and an approximation of the baseline width base on the FWHM, that is FWHM x 3.

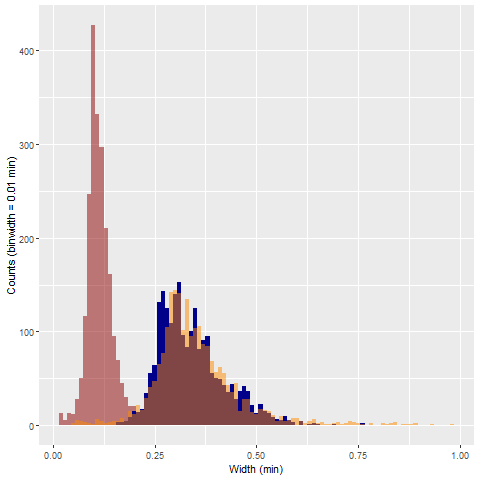

Here the orange distribution corresponds to an estimate of the baseline width using the FWHM. FWHM is a metric that is more easily and more accurately estimated by chromatographic peak picking software such as Skyline. We can see that an approximation using 3 times the FWHM has a very good overlay over the peak boundaries obtained from a curated Skyline document. Therefore, as a **rule of thumb** one could grab the **maximal FWHM** detected for all peaks and **multiply by three** to get a good initial value for the binning across the RT dimension parameter.

The histograms show that the 0.75 min binning range across the retention time (RT) dimension might have been excessively wide. After inspection of these results, we can conclude that the RT binning parameter could be reduced to ~0.675 min to selectively capture all MS/MS events associated to a specific precursor while minimizing interference of closely eluting isobaric species.

NOTE: this parameter is specific for the chromatographic conditions used to acquire the data of this tutorial. We recommend that these values should be adapted to each data set.

#### Quantitation considerations

##### Points across the peak vs. Precursor intensity

An ideal design of a DDA method that can be used for quantitative measurements is that which allows to obtain enough precursor ion measurements across the chromatographic peak of a peptide while using all the remaining time to obtain interpretable fragment ion spectra. Skyline allows us to assess how successful the MS method was at achieving this by using the **Points Across Peak** calculated in Skyline for all transitions/ion targets.

NOTE: Particularly for the case of cross-linked peptide analysis, we care to optimize these settings only for these peptides. Cross-linked peptides are only a small fraction of the lowest intensity peptides identified in a cross-linked peptide digest, especially for samples that have not been enriched in any way such as the BSA-DSBU digest analyzed in this tutorial. Since the Skyline document only contains the information of DSBU modified peptides, the information of high abundant unmodified peptides is omitted. However, if you work with analysis that includes both unmodified and cross-linked peptides you should make sure that the analysis is not biased by the peak widths of unmodified peptides.

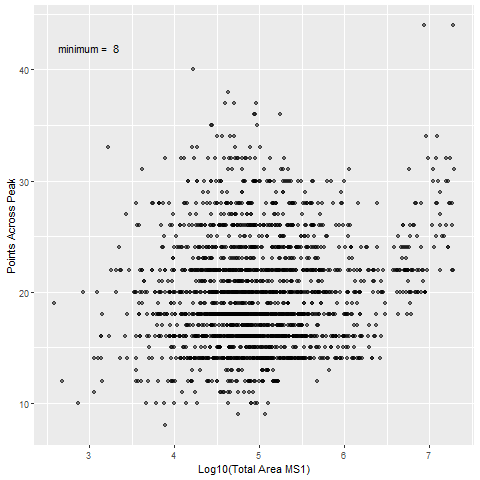

Having at least 9 points across the chromatographic peak is a good target to avoid the introduction of significant deviation to the estimated peak area due to insufficient sampling of the chromatographic peak^8^. With the two-dimensional histogram displayed above we can see that the method used for acquisition, which consisted of two MS1 TIMS ramps and 10 PASEF ramps of 166 ms each, ensured that most cross-linked peptides were profiled with 10 or more points across their respective chromatographic peaks. Therefore, we concluded that with the current chromatographic system the DDA-PASEF method was appropriate for performing MS1 quantitative comparisons of the measured cross-linked peptides.

The use of shorter gradients would require shortening the DDA cycle time, for example by reducing the amount of PASEF ramps per cycle to ensure enough points are captured. However, this would come at the cost of losing some depth in profiling.

##### Reproducibility of precursor ion peak areas

The DDA results here can be inspected for quantitative comparison of precursor ions. Here we will have a look at the reproducibility of the peak area integrated for the precursor ions associated to the identified cross-linked peptides. This will be presented with the coefficient of variation (CV%: standard deviation/mean x 100) of the respective areas for three group sets: 1) technical replicates after conditioning of instrument with 4 prior injections of a similar matrix (S-Trap Pool (quant)), 2) technical replicates after two blank measurements (S-Trap Pool (conditioning)), and 3) digestion sample preparation replicates (S-Trap Reps).

From a single BSA-DSBU cross-link reaction, three aliquots were split and processed in parallel with a S-Trap digestion protocol. Each of these bottom-up proteomics sample preparation replicates are referred to as S-Trap Reps. After reconstitution in aqueous 0.1% (v/v) trifluoroacetic acid and 30% (v/v) acetonitrile, equal aliquots were taken to make a pooled sample that we refer here as S-Trap Pool. The individual replicates and pools were frozen at -80 °C until required for measurement.

###### Conditioned S-Trap Pool CV%

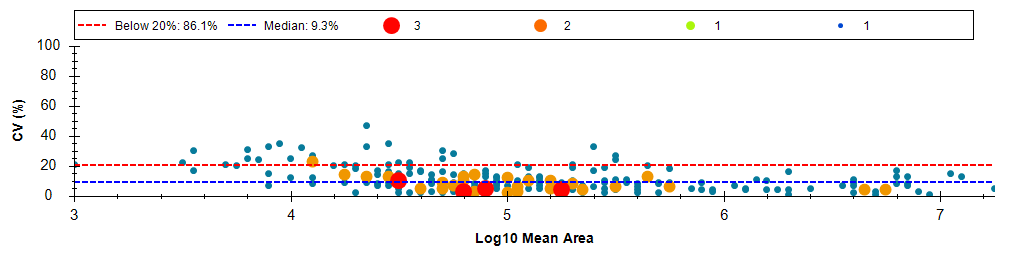

NOTE: This three-dimensional histogram was generated with Skyline after grouping the samples with respect to the groups defined above.

The CV% across instrument technical replicates (i.e., injection of same sample) showed that the majority of identified precursor ions could be measured with good reproducibility, indicating a suitable LC and MS method for measurement. This result is in contrast to recent evaluations of DDA reproducibility for cross-linked samples^9^ where DDA data was characterized as not suitable for analysis of quantitative cross-linking mass spectrometry experiments. The authors stated they did not perform peak boundary adjustment of their DDA data, so closer evaluation of the DDA data provided would have to be revised as this conclusion was achieved without the manual curation that we have encouraged through this tutorial. However, DDA-PASEF allows for faster cycle times without sacrificing depth on profiling and might be the true reason for the stark contrast.

###### Non-conditioned S-Trap Pool CV%

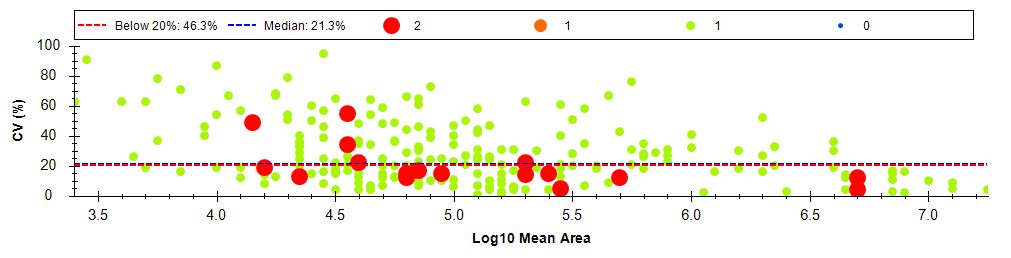

After the initial technical measurements were done, a blank run was performed, followed by instrument standby for about 8 hours. A new sequence was started with an initial blank and three additional injections of a fresh aliquot of the same digest. The CV% histogram shows a reduction in the quantitative reproducibility of the same peptides from the same sample which illustrates the importance of conditioning LC-MS system before extracting quantitative comparisons. Generally, we use three LC-MS injections prior to quantitative measurements, but this should be assessed per system and sample type. Knowing these limitations a priori helped us in using these “conditioning” injections for qualitative purposes. These replicates were used to expand the library of identified cross-linked by using different collision energy profiles (discussed in a later section).

###### S-Trap Replicates CV%

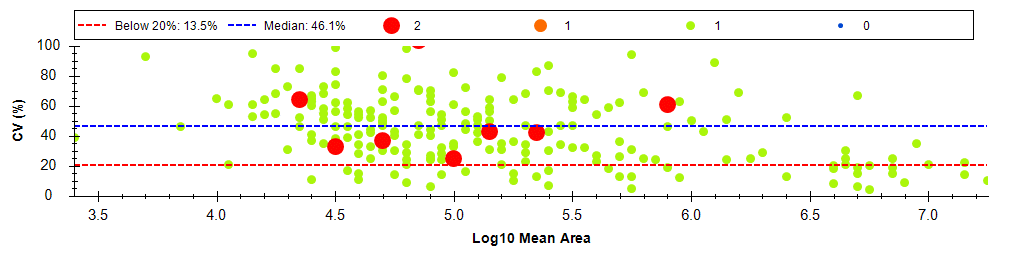

The reproducibility of the peptides across digestion replicates was lower than in the other two cases. By contrasting the larger variation of the sample preparation replicates with the instrument performance we could identify issues in the sample preparation reproducibility that we plan on addressing in future experiments. Based on the reviewed literature, the variability introduced by sample preparation has not been addressed with the level of detail that we can see with the EICs generated through Skyline. Most comparisons have remained centered at qualitative evaluation of cross-linked peptide identifications which is overshadowed by the data incompleteness issues introduced by DDA. However, as the focus of quantitative cross-linking mass spectrometry gains importance, evaluation of quantitative reproducibility affected by factors spanning from cross-linking reaction up to measurement will have to be addressed. With the current support for cross-linked peptides, Skyline presents itself as an ideal platform to perform these needed evaluations.

#### Design of DIA-PASEF windows

Once you have a curated list of cross-linked peptides you can use these results to tune a DIA-PASEF method that would allow for targeted detection of these peptides in an untargeted acquisition method. To minimize the cycle time of the DIA-PASEF method we will explore the ion mobility distribution of the identified precursors so that the detection boundaries are reduced to capture only the peptides of interest.

##### Ion mobility vs. m/z

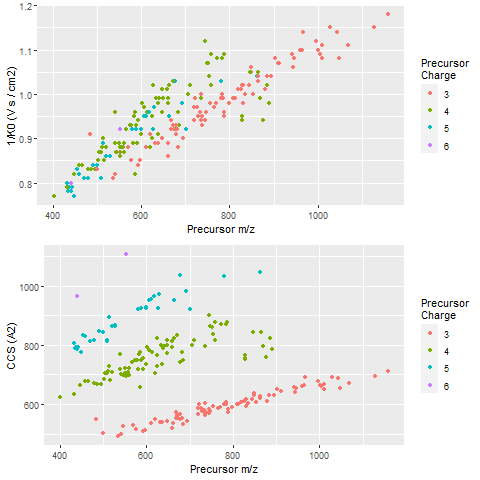

With the plots above we can appreciate the distribution of the identified ions and can use the following values to define the DIA-PASEF windows:

```
## [1] minimum m/z =  401.475697
## [1] maximum m/z =  1158.989187
## [1] minimum 1/K0 =  0.77
## [1] maximum 1/K0 =  1.18
```

These values would be useful to design a single DIA-PASEF *m/z* vs 1/K_0_ window scheme. First, let’s try to set some windows without the consideration of IMS and then contrast it with the same window scheme but shorter times enabled by ion trapping and separation in the TIMS cells.

###### Quadrupole windows scheme without IMS:

To retain a similar duty cycle as that of the DDA-PASEF method, (2 x MS1 x 166 ms) + (10 PASEF Ramps x 166 ms) ~ 2 s, while using the same TIMS trapping and separation time, we can split the quadrupole scanning range of 800 amu units into 80 m/z windows (10 PASEF ramps x 166 ms) windows while allowing two TIMS ramps for two Precursor scans (2 x MS1 x 166 ms) to keep the same sensitivity for precursor ions as that of the DDA-PASEF method. Inspection of the extreme values in Skyline, shows that the scanning range could be further reduced to a *m/z* range of 460 to 1075. The ions outside of this range correspond to cross-linked peptides that are supported by other precursor ions of similar or better intensities. This would further reduce the quadrupole windows to ~60 *m/z*.

**Quadrupole windows scheme with IMS:**

PASEF allows us to improve the quadrupole window scheme by knowing that specific ions are released from the TIMS cell **only** at specific points of the TIMS ramp. Let’s have a look at the averaged ion mobility spectrum across the relevant elution range of our peptides (20-70 min) generated with the timsTOF Pro instrument control software (timsControl).

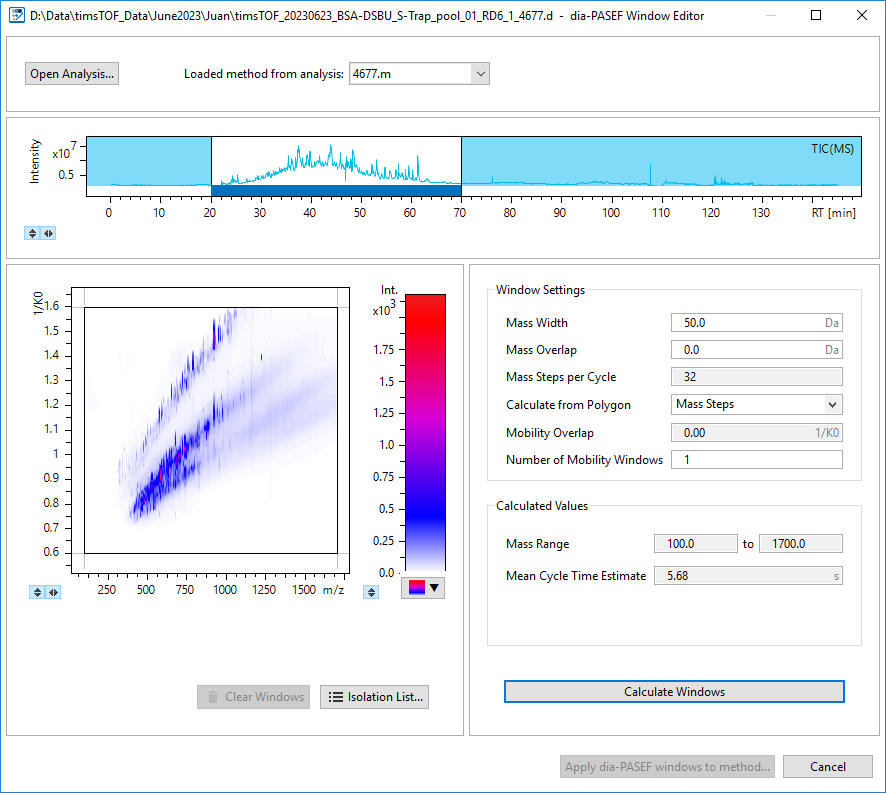

Here you can see that each ion is detected at only finite *m/z* and 1/K_0_ ranges. This can be used to maximize the use of each TIMS ramp by positioning the quadrupole at specific timepoints of the TIMS ramp only in positions where ions are expected to be released at that point. This concept is the key principle of PASEF^10^. This allows us to schedule multiple quadrupole scanning regions per TIMS ramp as illustrated below. Each color of the *m/z*-1/K_0_ isolation ranges correspond to a single PASEF ramp.

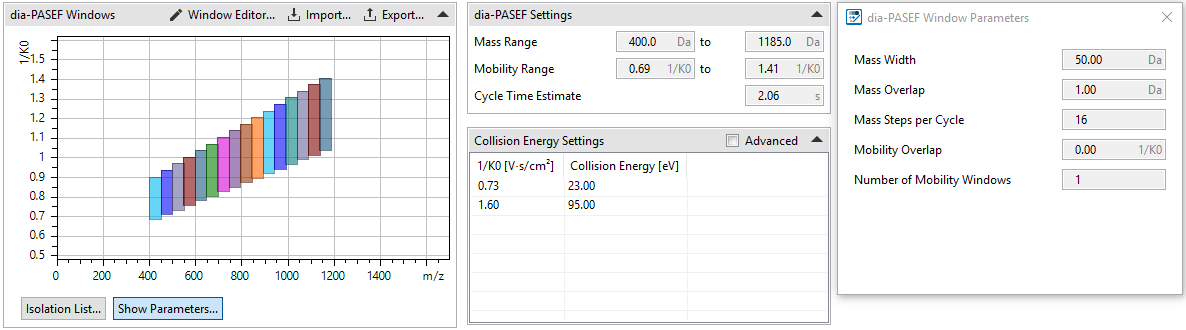

Here one precursor TIMS scan and 16 mass step cycles of 50 m/z each could be achieved in 2.06 s thanks to the DIA-PASEF scheduling. These quadrupole isolation windows of 50 *m/z* are still larger than those used in non-TIMS DIA workflows or fast gradient methods with TIMS-TOF instruments. However, cross-linking experiments still rely on “long” gradients (e.g. 3-35% 90 min gradients) and the additional dimension of ion mobility allows us to work with these wider windows since co-fragmentation can be partially resolved using ion mobility filters in Skyline while still allowing for higher ion accumulation times in the TIMS cells, boosting method sensitivity. The effects on cross-linked peptide detection has been evaluated recently^11^ and it showed that from a range of 10-to-50 *m/z* windows, only marginal differences were observed with respect to detected peptides most likely due to more selective and confident association of fragment ions to their respective precursor ions when smaller windows are used.

Although currently not supported by the acquisition and data analysis software, in the future the quadrupole isolation windows could be adapted to the distribution of ions across the chromatographic gradient, as demonstrated recently for other LC-MS/MS platforms^12^. As larger, more hydrophobic peptides elute later, the relevant ion population shifts towards larger *m/z* values as the gradient progresses.

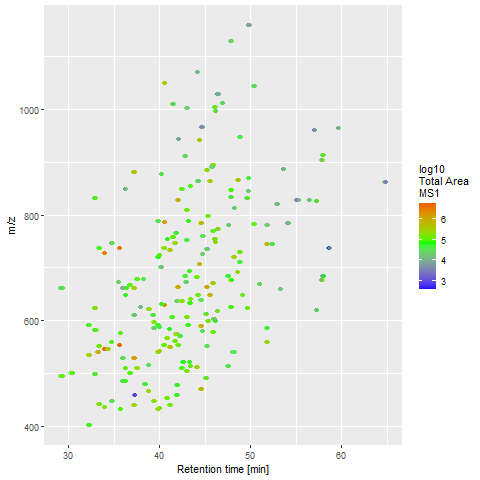

#### Evaluating rate of missing PSMs

Despite its limitations, DDA still provides great quality fragment ion spectra ideal for the annotation of peptides with complex fragmentation patterns such as those of cross-linked peptides. For those peptides that were detected with the lowest intensity precursor ions, the identification was only possible by concentrating all the fragment ion current generated for this precursor into a single summed MS/MS spectrum. This was further facilitated by the selective and efficient ion current sampling of the DDA-PASEF acquisition mode. However, this worked only for measurements where the precursor ions happened to be in the targets that the acquisition machine selected automatically. Even for instrumental replicate measurements, small deviations in intensity recorded and drifts in chromatography causes that the selected targets are not the same between replicates, particularly for the lower intensity ions. This is part of what is known as the data incompleteness of DDA data where fragment ion evidence is not recorded or identified on all measurements.

Let’s have a look at how many cross-link IDs were obtained from each replicate and the intersection of this. At the same time, let’s compare this to the complete list of identified cross-linked peptides using UpSet plots^13^.

We will do these comparisons only for the first four technical replicates measured at the same collision energy profile. These measurements were done after the chromatographic system had already been conditioned by four injections of a similar digest matrix to minimize variation in chromatography. In an ideal scenario, these would be identical measurements that would give the same results.

Keep in mind that when comparing results for cross-linking data you have to define at which level the overlap is evaluated. That is protein-protein, peptide1-peptide2 with or without site specificity, or the precursor ion of peptide1-peptide2 cross-link? The overlap will vary depending to the level of specificity and we’ll exemplify that here from highest overlap to lowest at the precursor level. Protein-protein overlaps are not relevant for this data set since it comes from a single protein cross-linking experiment.

##### UpSet: Technical Replicates - Cross-linked Peptides (No site specific)

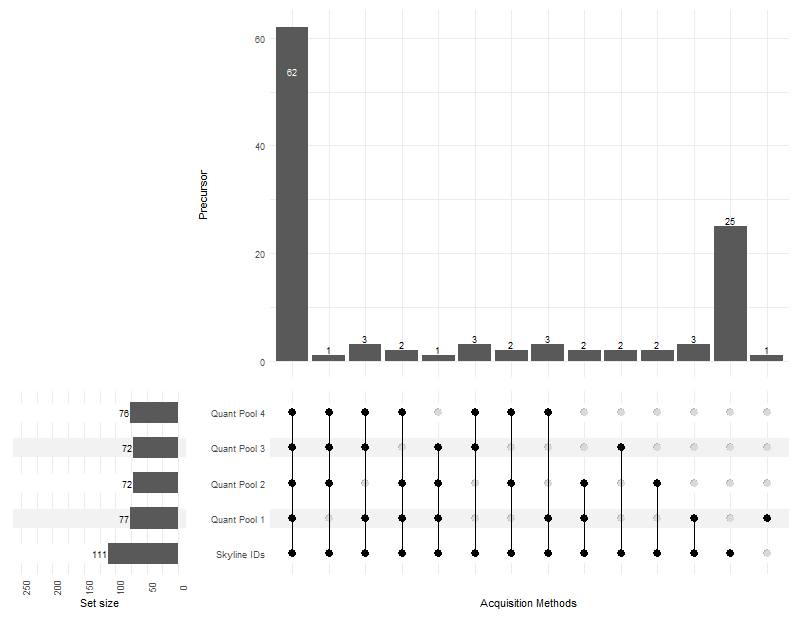

Here we can see that there were a total of 111 peptide1-peptide2 (without considering linkage specificity) combinations that remained in our final list of cross-linked peptides in Skyline. On average there were ~74 peptides identified per sample, however they were not all exactly the same across replicates. Only 62 peptide1-peptide2 combinations were found in all four replicates while the unique combinations are displayed by the connection matrix and the amounts displayed in the bars. The ID cross-section shows that most of the remaining cross-links were supported in more than one replicate, but there were still 7 peptide1-peptide2 combinations that had supporting information in a single replicate. Even in this summarized version of cross-link IDs that do not consider the linkage position or the specific precursor ion that supports their ID it is already possible to visualize the effects of DDA in identification reproducibility if you were to rely solely on the MS/MS spectra.

There were 25 peptide1-peptide2 combinations that were not identified in either of these technical replicates. The PSMs supporting these cross-links come from the other measurements acquired with different collision energy profiles or the individual sample preparation replicates; this will be discussed further below. There was a single peptide1-peptide2 proposal from the Quant Pool 1 replicate that passed the 1% FDR that was not included in the final list of cross-linked peptides. This corresponds to the DTHKSEIAHR-DTHKSEIAHR cross-link proposal which was deemed to not have PSM that allowed sufficient sequence coverage by fragment ions that were distinctly above the noise threshold.

Now let’s dive into evaluation of more specific comparisons while we map some metadata to the UpSet plots.

##### UpSet: Technical Replicates - Cross-linked Peptides (site specific)

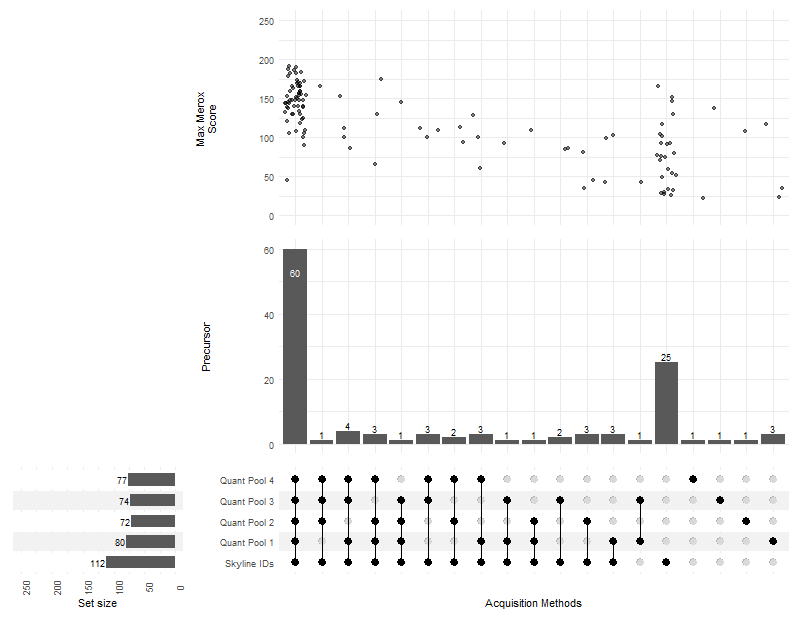

Now you see that when linkage position specificity is included a lower overlap is observed for this curated dataset. Ultimately, a decision of the linkage position was decided in Skyline by comparing all the PSMs available for a respective peptide1-peptide2 combination and deciding upon a single chromatographic peak that should have this linkage position. In this dataset you can have a look at **FKDLGEEHFK-DTHKSEIAHR-[+196.084792@2,2]** and **FKDLGEEHFK-DTHKSEIAHR-[+196.084792@2,4]** where both linkage are supported by multiple PSMs each and where detected as peptides eluting at different times. This final step of refinement can be translated back to the original MeroX results used to create the spectral library by making sure the PSMs reported correspond to the correct chromatographic peak where MeroX might struggle in assigning the correct linkage sites for MS/MS spectra containing fragment ions with low S/N.

From the new mapped metadata, we can appreciate the MeroX score distribution of the overlaps. As expected, those cross-linked peptides identified in all replicates tend to have the highest MeroX score distribution. At the same time, you can see that some of those cross-link IDs that come from single replicates also show up with a low-to-mid score. Finally, some of the IDs that were excluded from Skyline (right side columns) still had good MeroX scores (i.e. Score < 70). If you look closely into these cross-links in the final Skyline document, you will see that the peptide1-peptide2 combinations are duplicated and that actually a different linkage position was chosen to be kept in Skyline.

Finally, let’s have a look at the overlaps of identified precursor ions. This is perhaps the most useful comparison for MS method optimization since it allows mapping the intensity detected for these precursors and the distribution of charge states for which collision energies would be tuned.

##### UpSet: Technical Replicates - Precursor ions

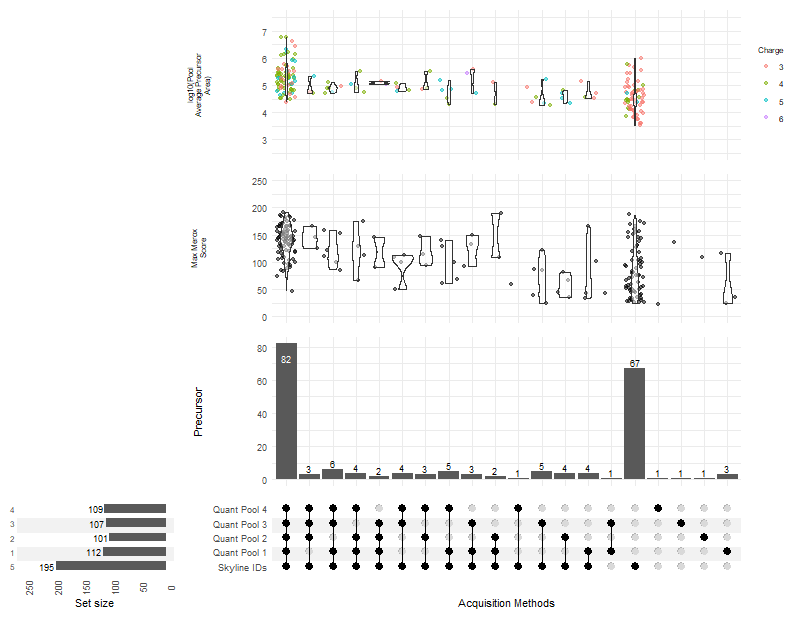

When comparing data at the precursor level the overlap is even smaller, but helps us in understanding some of the reasons for this which are discussed below.

##### Effects of DDA intensity bias on missing IDs

The last UpSet plot showed the logarithmic transformation with base ten of the average precursor area (of the first 4 technical replicates) for each of the identified precursors. This shows us that a great portion of the precursor ions identified in all replicates corresponds to the most intense ones (based on their integrated areas) which are the most likely ones to be selected by the instrument for fragmentation. On the other hand, identified precursors that have missing PSMs in other replicates are ranging on the lower end of the integrated areas distribution.

The degree of the intensity bias is dependent on the co-elution of analytes around the targets of interest. So let's see the retention times of the identified precursors and their integrated areas while we contrast it to the total ion current (TIC) and base peak ion chromatogram (BPC) of one of the technical replicates (Quant Pool 1). Here we use the BPC-to-TIC difference as a proxy measurement of how crowded is the *m/z* dimension due to the co-elution of multiple species.

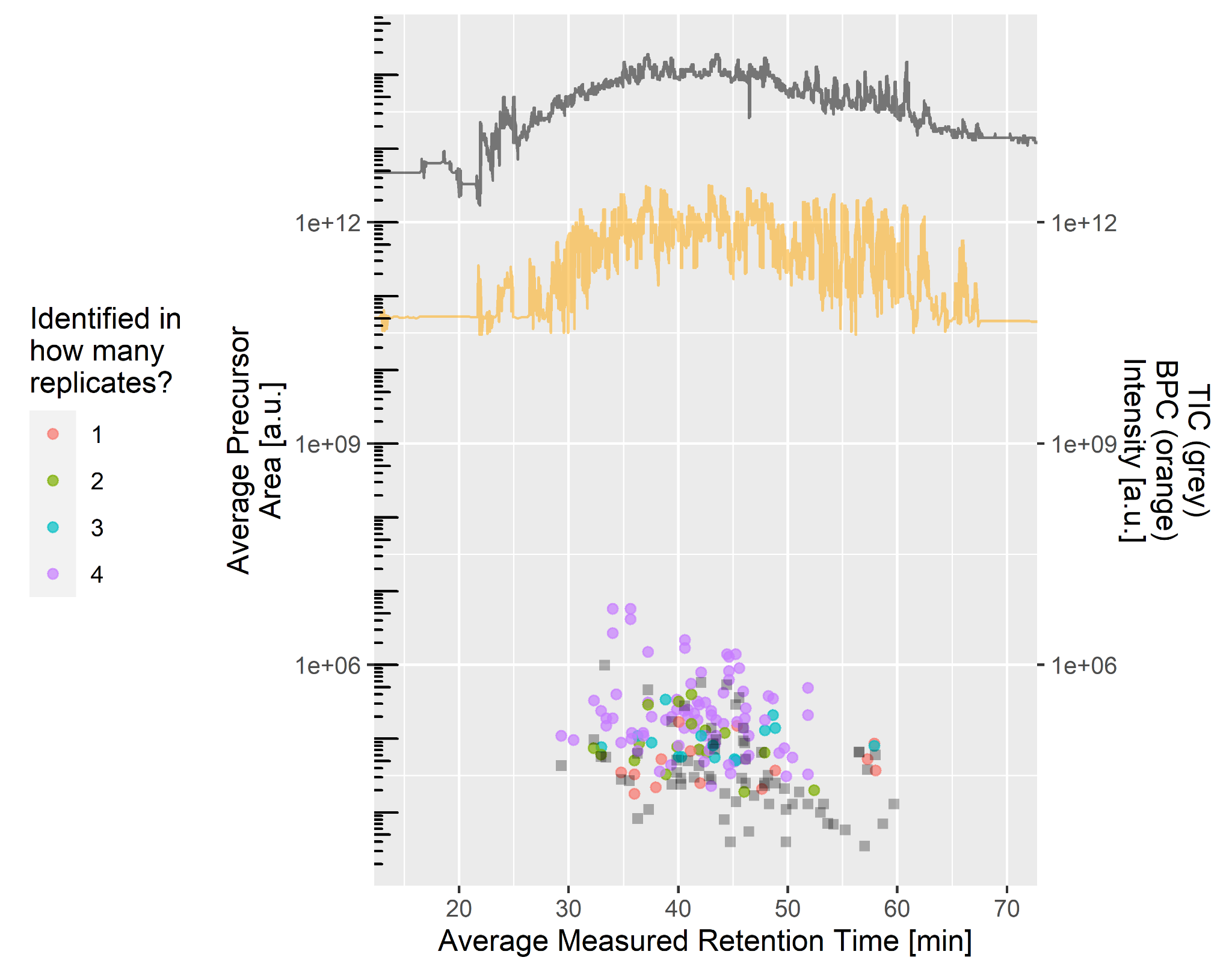

NOTE: for visualization purposes the y-axis is displayed with a logarithmic transformation of base 10, but we also provide the code for the linear axis representations.

Again, here we can see that the most abundant precursor ions are those identified in all 4 replicates. However, you can see that some intense precursors were still not identified in all replicates (green and blue dots). By overlaying the data to the TIC and BPC you can see that all of these precursor ions were identified in a region where the maximal TIC was recorded, corresponding to where most analytes co-elute. The large gap between the TIC and BPC shows that the total ion current is not dominated by a single ion. So, despite the ions being intense enough to provide identifiable spectra, these were most likely not chosen in the replicates with the missing IDs. Also, as observed in the UpSet plots, those precursors identified in a single replicate are in the lower range of detectable ions.

Interestingly, most of the missing IDs from the first 4 technical replicates (illustrated as grey squares) are on the lower range of detectable ions, suggesting, not surprisingly, that intensity is the main restriction for identifying ions in all replicates. However, below we discuss an additional factor that contributes to this, collision energy optimization, which explains the missing IDs in all replicates of ions that were in the mid-to-high range of detectable ions.

##### Effects of collision energy on missing IDs

Additionally, color coding the precursor ions based on their charge states shows us that most of the precursor ions that were not identified in these technical replicates are mainly triply charged ions. The collision energy (CE) profile used for these technical replicates was a collision energy profile used for general proteomics applications in timsTOF Pro instruments labelled as low CE Profile in the **Collision Energy Profile** table.

NOTE: In timsTOF instruments, the collision energy is selected based on the ion’s inverse ion mobility (1/K_0_) which in turn has a dependency on its *m/z*, i.e. within a same charge state ions the ions with the largest *m/z* have the largest 1/K_0_. Ions with the largest 1/K_0_ are those that travel farthest into the TIMS cell and are the first ones to be released from the TIMS cell into the quadrupole so the collision energy profile is set from largest CE to lowest.

| Settings | CE Start (eV) | CE End (eV) | 1/K0 Start  (Vs/cm^2^) | 1/K0 End  (Vs/cm^2^) |
| --- | --- | --- | --- | --- |
| Low CE Profile | 59 | 23 | 1.60 | 0.73 |
| Mid CE Profile | 75 | 23 | 1.60 | 0.73 |
| High CE Profile | 95 | 23 | 1.60 | 0.73 |

Preliminary analysis of the initial replicates showed the charge state distribution of the identified precursors trended towards high charge states (4+ or higher). This was despite the detection of lower charge state ions for the same peptides when the corresponding EIC traces were aligned in Skyline. Therefore, the subsequent measurements included in this data set explored collision energy profiles where the CE applied to higher *m/z* ions was increased by using a CE profile used previously^14^ and an intermediate CE profile. The inclusion of measurements (not displayed in the UpSet plot displayed above for simplicity) with a systematic increase to the CE applied resulted in the addition of new PSMs of lower charge state ions. When comparing the charge states of the same analyte, lower charge states require a higher collision energy to induce efficient fragmentation than higher charge states. Some of these PSMs provided redundant information to some already identified cross-links while also providing new cross-link peptides as visualized in the first UpSet plots. On the other side, using higher collision energies for some of the precursors ions identified before showed a decrease in the quality of the fragment ion spectra due to over-fragmentation; loss of characteristic cross-linker MS-cleavable peptide doublet ions, complete loss of any precursor ion, and noisy spectra despite detecting intense precursor ions in the MS scan. There is no ideal fragmentation energy setting for all possible peptides and their respective charge states. The data shown here suggests that a larger precursor identification range might be achieved with spectral libraries generated from measurements that used multiple collision energy profiles.

Moreover, with the visualization of the EIC traces in Skyline is possible to detect the dominant charge state of a specific cross-linked peptide. This knowledge in combination with a multiple CE spectral library allows creating selective, non-redundant PRM-PASEF methods where only the most intense precursor ion can be selected using the collision energy that generate the most reliable PSM.

#### What to do with single replicate PSMs with good MS/MS spectral score?

A common practice in the high-throughput analysis of cross-linking mass spectrometry data has been to exclude identifications that were only found in one replicate measurement. This is enforced as an attempt to exclude false positive identifications that arise from random matches to the *m/z* vs intensity matrix of recorded MS/MS spectra. As these matches are random, they are very unlikely to be replicated. More recently, results filtration with a false discovery rate (FDR) using a target-decoy approach has been implemented in cross-linking workflows after these have demonstrated their usefulness in other bottom-up proteomics fields. However, the strength of a reliable FDR control depends on the amount of data points of “true hits” so that the identification score (e.g. MeroX score) of these can be used to contrast it to the score distribution of artificial decoy peptide sequences. For single or few protein systems (as is common in initial *in-vitro* experiments) where few interaction points are expected this approach loses power in correctly identifying a score threshold to discriminate false positives with defined error rate. This problem is further increased when filtering is done per LC-MS/MS file instead of whole dataset generated by replicate measurements and even more so when MS/MS summation is performed as described in this work.

So how can we reconcile the approach followed in this work that improves the S/N ratio of detected fragment ions at the expense of reducing matches per file to (ideally) single PSMs? What should we do with reliable cross-link identifications that show evidence of MS-cleavable ions of both peptide chains, full sequence coverage of these, and localization of cross-link sites, but were identified in a single replicate?

As you have already noticed from the EICs of the final list of cross-linked peptides in the final Skyline document, all peptides presented here showed matching peptide elutions across all replicates. MS/MS fragment ion spectra is only part of the information collected for peptides separated and analyzed by LC-IMS-MS/MS. Skyline is an ideal platform that allows pooling and visualizing several characteristic features of these peptides:

- Precursor ion exact *m/z*; based on their matched elemental composition and charge state
- Isotopic distribution; again based on their matched elemental composition and charge state
- Consistent elution time and order; defined by the peptide-specific interactions to the chromatographic system employed and for which drifts can be accounted with iRT peptides
- Ion mobility of detected precursors; defined by the collisional cross section of the ion which in turn has a dependency on its *m/z*
- Fragment ion spectra; defined by the cross-linked peptide sequences, cross-linker, and collision energy employed
- (Optional) Uniqueness of signal from cross-link reaction; random matches to wrongly sequence MS/MS spectra of unmodified peptides can be quickly be excluded by running negative controls

These were the critical parameters evaluated for each cross-linked peptide to confirm or discard its detection the samples measured. Although it is possible to train chromatographic peak picking models that consider these parameters (and more), currently this is not possible for cross-linked peptides since Skyline still needs to implement the calculation of decoy cross-linked peptides. But eventually this can also be implemented to do peptide feature detection FDR control where PSMs become part of the decision making and not the only limiting

### Conclusions

Congratulations! You have completed your first cross-linking mass spectrometry data analysis with DDA-PASEF, MeroX, and Skyline. You learned how to process raw DDA-PASEF data to generate IMS-filtered spectra into text peak lists compatible with most database search engines. You learned how to perform XL-peptides database search with MeroX and how to standardize these results with proXL for follow-up analysis and spectral library creation with Skyline. You learned how to import these DDA identified XL-peptides targets into Skyline. You learned how to import custom ion mobility libraries and how you can manually edit this to account for errors in the mobility values selected. And, finally, you saw some examples where the complex information collected in LC-IMS-MS/MS experiments can be explored within a single software platform to accept or reject the identifications proposed by spectral-centric database search software.

The inclusion of ion mobility information to your data and the consideration of indexed retention times increases the complexity of the analysis. However, we found that many low abundant precursor ions could only be identified due to the efficient ion current sampling available through PASEF and decomplexing what would have been chimeric spectra without the use of the additional IMS dimension. Even with these enhancements many cross-links remained uniquely identified in some replicates. For these latter cases and IDs obtained with only a particular collision energy profile, the alignment of results using a calibrated retention time calculator helped in dealing with the known limitations of DDA data due to its stochastic nature.

The data presented here is not without its flaws, as you might have noticed with the quantitative reproducibility of the sample preparation replicates, but by using the deep inspection tools that Skyline provides we were able to recognize these issues and show how you could use the same tools to evaluate your samples. Future experiments will address improvement of sample preparation reproducibility. On the other hand, we have shown here how it is possible to improve the qualitative aspect of DDA data with a peptide-centric analysis where we used the retention time, ion mobility and MS/MS ID alignment to carefully align peptide detection across multiple replicates.

Additionally, we encourage the cross-linking community to adapt Skyline to their analysis. Although we showed how data processed only with MeroX could be analyzed, the modular aspect of spectral library creation from Skyline allows for analyzing the same data with multiple cross-linking database search engines as long as their outputs are converted to proXL format. This multi-search engine approach has already proven to be an effective way of accurately controlling the false discovery rate of search engines^7^. Skyline is an ideal platform that allows combining results from multiple search engines and can be used to further filter the targets that have to be evaluated manually or increase the reliability of fully automatic data processing. Moreover, once your analysis is done, the results can easily be shared in a transparent way to the community with the use of Panorama^15^.
